# Reframing Psychiatry for Precision Medicine

**DOI:** 10.1101/2020.07.19.210971

**Authors:** Elizabeth B Torres

## Abstract

The art of observing and describing behaviors has driven diagnosis and informed basic science in Psychiatry. In recent times, studies of mental illness are focused on understanding the brain’s neurobiology but there is a paucity of information on the potential contributions from peripheral activity to mental health. In Precision Medicine, this common practice leaves a gap between bodily behaviors and genomics that we here propose to address with a new layer of inquiry that includes genes’ expression on tissues inclusive of brain, heart, muscle-skeletal and organs for vital bodily functions. We interrogate genes’ expression on human tissue as a function of disease-associated genes. By removing genes linked to disease from the typical human set, and recomputing the genes’ expressions on the tissues, we can compare the outcomes across mental illnesses, well-known neurological conditions, and non-neurological ones. We find that major neuropsychiatric conditions that are behaviorally defined today (e.g. Autism, Schizophrenia, Depression) through DSM-observation criteria, have strong convergence with well-known neurological ones (e.g. Ataxias, Parkinson), but less overlap with non-neurological ones. Surprisingly, tissues majorly involved in the central control, coordination, adaptation and learning of movements, emotion and memory are maximally affected in psychiatric diagnoses along with peripheral heart and muscle-skeletal tissues. Our results underscore the importance of considering both the brain-body connection and the contributions of the peripheral nervous systems to mental health.

## 1. Introduction

Modern medicine is at an inflexion point [1], whereby advances in computational methods, wearable sensing technology and open access to Big Data are reshaping the ways in which we inform basic science and rapidly translate our knowledge to actionable treatments. Psychiatry is one of those medical fields that is rapidly evolving, while adapting traditional models to help advance their main goal of helping patients improve quality of life. Along those lines, Computational Psychiatry [2] a nascent sub-field within Psychiatry, is merging methods from Computational Neuroscience with clinical approaches, through successful collaborations. These new developments are bound to open new frontiers in therapeutic treatments. And as part of a more general effort in the medical field, Precision Medicine (PM) [1] has emerged as a new platform to combine expertise from multiple layers of the knowledge network, to ultimately design personalized targeted treatments **(Figure 1A).** Integrating the personalized concept of PM with the new advances in Computational Psychiatry could give us a new way to approach mental illness and help patients cope with lifelong changing needs.

**Figure 1.**
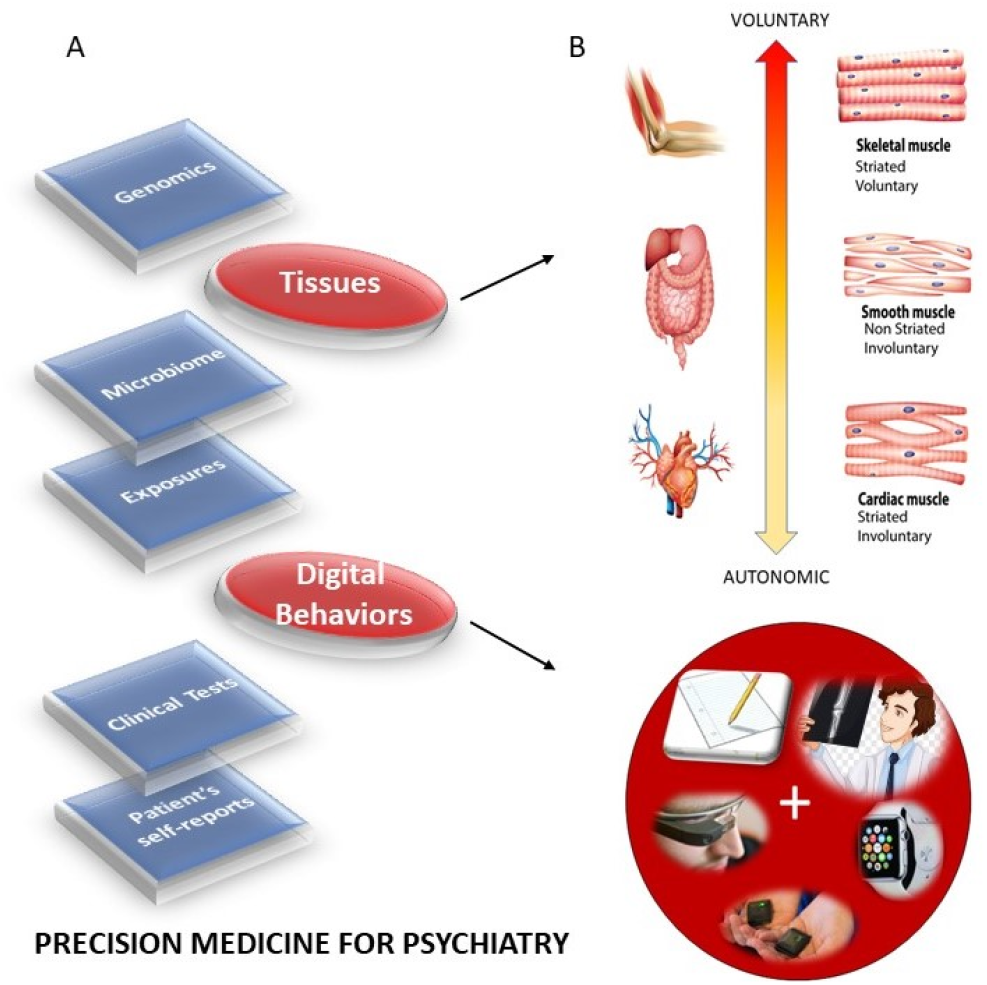
Roadmap to implement the Precision Medicine Model for diagnoses and treatments of mental illnesses. (A) The PM’s interconnected knowledge network can contribute information about the individual’s medical history, behaviors, environment, microbiome, and genetic makeup. Importantly, the new proposed layer of *digitized behaviors* leveraging the wearable biosensors revolution can transform medicine by creating truly personalized assessments. Additionally, the layer of behaviors can be connected to the nervous systems functioning via fundamental levels of neuromotor control that span along a phylogenetically orderly taxonomy. (B) This taxonomy is based on levels of maturation in autonomous neuromotor control, linked to three fundamental muscle types: autonomic (by cardiac muscles); involuntary (by smooth muscles) and voluntary (by skeletal muscles.) By linking the fundamental muscle types to the levels of control in the nervous systems, digital behaviors can then be mapped to bodily autonomy, bodily autonomy mapped to muscle types and muscle types to genes/proteins. Any measure of treatment’s effectiveness for mental illnesses can then map back to improvements in observable behaviors embedded in activities of daily social life.

The task ahead is challenging because there is no proper roadmap to connect the layers of the knowledge network in PM and produce personalized diagnoses and measures of treatment outcomes that truly separate disease progression from treatment’s effectiveness, according to age and development. Part of the problem is that most brain science has focused on experimental assays and methods that curtail natural movements. As such, our knowledge about the dynamics of natural behaviors is very limited, particularly in reference to those aspects of behavior that remain hidden to the naked eye of the clinician-trained to observe specific expected behavioral landmarks of a Psychiatric disorder, conceived exclusively as a mental illness. In so doing, the clinically trained eye may miss important information that is perhaps common across different disorders of the nervous systems and rather relevant to help improve the patient’s quality of life. For example, motor coordination and volitional control are critical ingredients of autonomy in any natural behavior underlying activities of daily living. Yet, these are not considered part of these diagnostics criteria for mental illnesses such as Autism, Schizophrenia and Depression-as per the Diagnostics Statistical Manual (DSM-5) [3] (and see Supplementary Material.)

Research on the underlying neurobiology of mental illnesses has revealed their associated genetics and/or helped characterize patterns of brain activity in response to external stimuli (while curtailing naturalistic bodily motions to avoid instrumentation artifacts in imaging data or in EEG, MEG, etc.) This central approach to brain science has left us with a paucity of information about the possible contributions to mental illness from the peripheral nervous systems, and from vital organs important for autonomous living. The peripheral activity, however, continuously feeds back to the brain via afferent (body-to-brain) channels and is in turn, dynamically updated through efferent (brain-to-body) activity, self-generated by the system itself. This recursive loop, whereby re-entrant information that is partly self-produced by the organism and partly influenced by external environmental conditions, would provide important clues about truly evolving dynamics and stochastic (variability) across all-natural behaviors. Approaching the problem through this lens could bring a new quantifiable layer of granularity to basic research. This would include the design of age-appropriate metrics reflecting the development of the organism as it ages and as it copes with a disorder. The micro- and macro-motion data from the nervous systems biorhythms is the low hanging fruit that we can easily attain by leveraging the wearable sensors revolution. And because these quantifiable digitized activities and signals therein are partly self-generated, self-monitored and self-corrected by and within the nervous systems, this quantitative approach has the potential to switching us from a purely correlational science to a science that is based on causal relations between nervous systems activities and external / contextual stimuli. In this sense, the new proposed approach to mental illness is amenable to intervene and modify the system with well-informed, near-optimal means capable of improving its performance.

Micro- and macro-motions that underly all aspects of human behavior depend on the intactness of fundamental tissues, many of which have already been characterized in genomics. Here we combine these underlying aspects of behavior with the current genomics knowledge, to inform Psychiatry of possible ways to improve quantification of nervous systems activities. Ultimately, we seek to build accommodations and support for the patient population, while reconceptualizing mental illness as a physically quantifiable disorder of *the nervous systems*.

The nervous systems already offer a taxonomy of function and control that is phylogenetically ordered and well organized along several axes. Some of these axes are accessible today with non-invasive means and as such, we can obtain signals and build computational models to understand mechanisms and translate them to actionable societal solutions. One possible orderly structure is suggested in **Figure 1B**, where we propose to map levels of neuromotor control (voluntary, involuntary, and autonomic) to fundamental types of muscles (skeletal, smooth, and cardiac) linked to commonly sampled tissues in genomic datasets. Combining information about genes expression on tissues that involve key components of the central nervous systems (the brain and the spinal cord), key organs for vital bodily functions (including smooth muscle lining internal organs), muscle skeletal tissues and nerves, and cardiac tissues (for autonomic heart functioning), we explore the effects of removing disease-associated genes, on the overall remaining genome expression on these tissues. As a first step in this exercise, we reasoned that the genes associated to a given disorder ought to be important in the functioning of certain systems, which in turn depend on certain tissues. We also reasoned that such stochastic variations and combinations could be measured relative to the presence of all genes and to the absence of genes across neurological or non-neurological conditions.

*What is the tissue distribution of genes’ expression in neuropsychiatric disorders like Autism, Schizophrenia and Depression, in relation to well-characterized neurological conditions? Is there convergence in the remaining genes’ expression on the tissues, upon removal of the genes associated to that disease? Furthermore, how would the genes’ expression change across the tissues in non-neurological conditions like various forms of cancer, immunodeficiencies, endocrine systems deficiencies and so forth? How would it change in acquired disorders like PTSD, currently diagnosed through observation?*

Take Autism for example. Autism is an umbrella term for a very heterogeneous set of neurodevelopmental disorders, but no gold standard criteria includes core neurological symptoms that could help us create early accommodations and support for the nascent nervous systems of the infant (during pre-cognitive stages of neurodevelopment.) The rule of thumb is to assume that the child has odd, socially inappropriate behaviors and that they should be modified through operant and cognitive conditioning techniques-often translated from lab animals to human babies, without any type of collaboration with other fields studying infant development. Current methods of diagnoses and treatments in autism are not based on normative neurodevelopmental data charts to understand age-dependent departures from typical neurodevelopment. Without any systematic way to build age-appropriate metrics, to capture highly non-linear, stochastic patterns and rates of change in the (rather accelerated) infant neurodevelopment, entire generations of infants, children and adolescents have been exposed to such means of behavioral treatments and no information can tie these back to the underlying genomic pool of this population.

In Schizophrenia, delusions, avolition and catatonia are at the core of the disorder, but as in Autism above, no criteria in the DSM highlights the profound somatic sensory-motor issues that have been found in patients [4]-even without the use of psychotropic medications, known to alter motility. Interestingly, historical accounts of Psychiatry (in pre-Freudian times) show the reliance on motor aspects of the behaviors that defined several mental illness from a neurological perspective [5].

Depression is also currently treated purely as a mental illness, but it may be important to understand potential contributions to various forms of depression, from the peripheral nervous systems and from the body in general. Genetic information may give us a way to link tissues affected in these neuropsychiatric conditions with those affected in neurological ones, for which treatments and interventions of various forms may be effective. These may be in the form of drugs, or in the form of physical, mindfulness and occupational therapies aimed at helping support the person’s bodily autonomy and overall increase the chances for independent living.

We here offer a new lens to help balance psychiatric with neurological criteria derived from genomic information specific to each disorder. In a first (crude) step of many to come, we start by comparing across well-known neuropsychiatric and neurological conditions, the results from eliminating the genes associated with each disorder and quantifying the degree of convergence in the maximally affected tissues, in relation to those resulting from eliminating the genes associated with non-neurological ones. We focus our discussion on possible ways to continue this path of inquiry and highlight current caveats for future improved iterations of the proposed methods.

## 2. Materials and Methods

We combine the data sets from genes associated with mental illnesses, with well-known neurological disorders and with illnesses that are not directly associated with the nervous systems. We also include genes associated with manifestations of acquired Post Traumatic Stress Syndrome Disorders (PTSD). Among mental illnesses defined by the DSM-5, we include Autism, Schizophrenia and Mental Depression of different types, (*e.g*. general, bipolar and unipolar). Among neurological conditions we include Ataxias (e.g. cerebellar, spino-cerebellar, progressive, and gait) and Parkinson’s disease. Among non-neurological disorders we include colon cancer, breast cancer, diabetes, congenital heart disease, hematologic neoplasm, and various auto-immune ones (lupus systemic erythematosus, psoriasis, and irritable bowel syndrome.)

We use as reference the genes, genes’ expression, and tissues from the GTEx Portal human RNA-Seq (Transcripts Per Million TPM)^1^ specifically using the files denoted in the Appendix B. In Autism, we use the genes scoring module of the Simons Foundation Autism Research Initiative (SFARI) scored according to evidence from the literature. We also use Ataxia genes, the X-genes and the FX-genes taken from various literature reviews [6, 7]. Furthermore, we use genes associated to mitochondrial disorders [8] and genes identified in Parkinson’s disease, taken from [9–14]. Besides the Autism SFARI genes and the genes reported in literature reviews, we take the genes associated to Autism, Schizophrenia and Depression reported in https://www.disgenet.org/home/ along with other genes from the above mentioned non-neurological disorders. The latter will inform us of fundamental differences in genes’ expression between these diseases and those which affect the neuromotor control and basic functioning, as mediated by interactions between the brain and the peripheral nervous systems (including the autonomic nervous system.)

The SFARI Autism categories that we used were those reported as of 03-04-2020. Quoting from their site:

- CATEGORY 1 Genes in this category are all found on the SPARK gene list. Each of these genes has been clearly implicated in ASD—typically by the presence of at least three de novo likely-gene-disrupting mutations being reported in the literature—and such mutations identified in the sequencing of the SPARK cohort are typically returned to the participants. Some of these genes meet the most rigorous threshold of genome-wide significance; all at least meet a threshold false discovery rate of < 0.1.
- CATEGORY 2 Genes with two reported de novo likely-gene-disrupting mutations. A gene uniquely implicated by a genome-wide association study, either reaching genome-wide significance or, if not, consistently replicated and accompanied by evidence that the risk variant has a functional effect.
- CATEGORY 3 Genes with a single reported de novo likely-gene-disrupting mutation. Evidence from a significant but unreplicated association study, or a series of rare inherited mutations for which there is not a rigorous statistical comparison with controls.
- SYNDROMIC The syndromic category includes mutations that are associated with a substantial degree of increased risk and consistently linked to additional characteristics not required for an ASD diagnosis. If there is independent evidence implicating a gene in idiopathic ASD, it will be listed as “#S” (e.g., 2S, 3S). If there is no such independent evidence, the gene will be listed simply as “S”.

The GTEx dataset is as of 06-05-2017 v8 release. For every gene in the Autism, Ataxia, X, FX, Mitochondrial diseases, Parkinson’s disease, and the non-neurological diseases, we first confirmed the presence of the gene in the GTEx data set and then incorporated it to the analyses.

The genes from the DisGeNet portal were found by interrogation of their data set under disease type and saving the outcome to excel files containing all pertinent information. All sample files used in our analyses are provided under Supplementary Materials.

### Count Normalization

The GTEx matrix of RNA-seq genes along the rows (*56,146*) x the tissues (*54*) along the columns was transposed (*54 x 56,246*), such that we expressed each tissue as a function of the gene expression denoted by the count (TPM). Each individual count value was then normalized using equation (1)

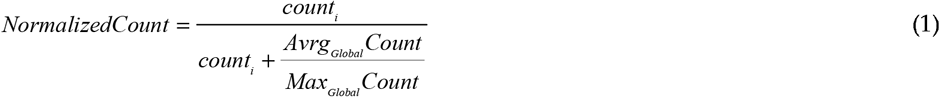

Here *count* is the count value of the *gene_i_, Avrg_Global_Count* is the overall average of the matrix of values taken along the columns and the rows. *Max_Global_Count* is the maximum count value, also taken globally across the matrix values. Figure 2 shows the original count numbers (Figure 2A) and the normalized version (coined Micro-Movement Spikes MMS) in Figure 2B. Figure 2C shows the MMS derived from the fluctuations in counts normalized by equation (1) while Figure 2D shows the histograms of the peaks (marked in red dots) for different tissues and genes scored by SFARI.

**Figure 2.**
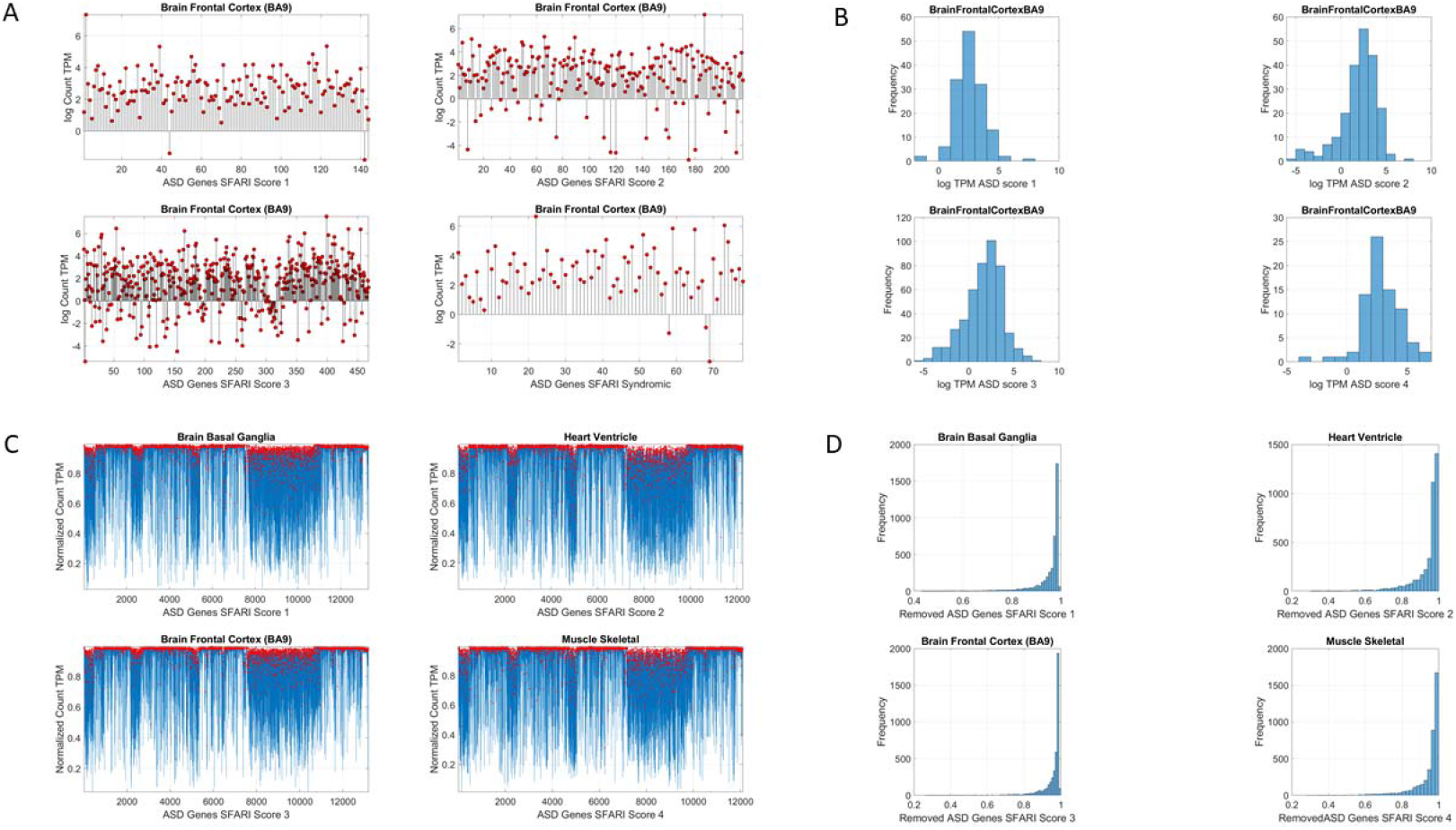
Analytical methods. (A) Sample raw data consisting of log count (TPM) for different scored genes expressed in the brain frontal cortex (Brodmann Area 9.) (B) Histograms of the log Count TPM for each case in (A). (C) Upon removal of the SFARI Autism genes, micro-fluctuations’ spikes in the normalized count, with deviations taken relative to empirically estimated mean, global averaged count, and global maximal count in Equation (1). (D) Histograms of the normalized micro-fluctuations’ spikes.

### Genes Removal

For each of the disorders of interest in Tables 2 (Mental Illness and Neurological) and Table 3 (Non-Neurological), we remove from the human GTEx data the set the genes associated with each condition, as reported in the various databases (SFARI, DisGeNet https://www.disgenet.org/home/ and the literature meta reviews.) We then treat the resulting count series as a random process. We use the exponential distribution to characterize it and to assess the differential expression across the tissues relative to non-removal in the original human genome.

**Table 2.** Genes distributions used in the removal process and literature sources.

| Neurological | Number of Genes | Source |
| --- | --- | --- |
| Parkinson | 17 | Lit Review [9, 11–14] |
| Ataxia Autosomal Recessive | 70 | Lit Review [6, 7] |
| Ataxia Autosomal Dominant | 46 | Lit Review [6, 7] |
| X-Chromosome | 6 | Lit Review [6, 7] |
| Fragile X (SFARI scores) | 73<sup>1</sup> (17,11,30,15) | SFARI Genes Module |
| Fragile X Syndrome | 194 | DisGeNet |
| FXTAS | 62 | DisGeNet |
| Ataxia | 813 | DisGeNet |
| Cerebellar Ataxia | 421 | DisGeNet |
| Gait Ataxia | 159 | DisGeNet |
| Progressive Cerebellar Ataxia | 134 | DisGeNet |
| Spinocerebellar Ataxia | 145 | DisGeNet |
| Neuropsychiatric (DSM) | Number of Genes | Source |
| Autism (SFARI scores) | 906<sup>1</sup><br>(144,216,468,78) | SFARI Genes Module |
| Autistic Disorder | 1,043 | DisGeNet |
| Schizophrenia | 2,697 | Lit Review [15–20] and DisGeNet |
| Mental Depression | 1,468 | DisGeNet |
| Depression Unipolar | 641 | DisGeNet |
| Depression Bipolar | 116 | DisGeNet |

<sup>1</sup> SFARI scores for Autism and FX genes were also used to assess each scored module separately. FXTAS stands for Fragile X Tremor Ataxia Syndrome

**Table 3.** Genes associated with non-neurological diseases.

| Non-Neurological | Number of Genes | Source |
| --- | --- | --- |
| Colon Cancer | 3,669 | DisGeNet |
| Diabetes Mellitus (noninsulin dependent) | 3,134 | DisGeNet |
| Oestrogen Receptor Positive Breast Cancer | 510 | DisGeNet |
| Congenital Heart Disease | 267 | DisGeNet |
| Hematologic Neoplasm | 827 | DisGeNet |
| Systemic Lupus Erythematosus | 1,883 | DisGeNet |
| Psoriasis | 1,308 | DisGeNet |
| Irritable Bowel Syndrome | 1,483 | DisGeNet |
| Mixed | Number of Genes | Source |
| Mitochondria | 41 | Lit Review [8] |
| Mitochondrial Myopathies | 121 | DisGeNet |
| Mitochondrial Diseases | 284 | DisGeNet |
| MELAS Syndrome | 81 | DisGeNet |
| PTSD | 418 | DisGeNet |

MELAS (a form of dementia) stands for Mitochondrial Encephalopathy, Lactic Acidosis, and Stroke-like episodes.

The question that we ask is, given the known neurological phenotypes, is there convergence between the most affected tissues upon associated genes’ removal and the changes in the tissues that will be obtained by the removal of genes associated with mental illnesses? Furthermore, is there convergence with the outcome from removing the genes associated with non-neurological illnesses? Tables 2–3 show the number of genes removed in each respective case. It also shows the source of the reported genes associated with each condition/disease.

### Stochastic Analyses

Since the count values for each tissue can be conceived as a random series of numbers, we use maximum likelihood estimation (MLE) to model the numbers representing the counts, as generated by the exponential distribution using equation (2)

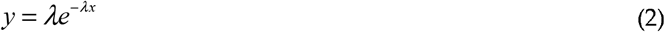

Here *x* represents the normalized count value (as per Equation (1)) and *y* the value from the exponential distribution. We seek the value of *λ* to model this counting random process representing the genes’ expression in the tissue. To that end, we estimate the likelihood *L*(*λ|x*_1_, *x*_2_,…,*x_n_*) where the series of counts *x_i_*, with *i* ranging from *1* to *n* represent the normalized counts (according to Equation 1) across all genes for one tissue. Appendix B shows the steps to find λ. And this is computed for each of the 54 tissues. We then rank the departure of λ (resulting from genes removal) from the λ obtained for the full human genome (see below). This is explained in Figure 3.

**Figure 3.**
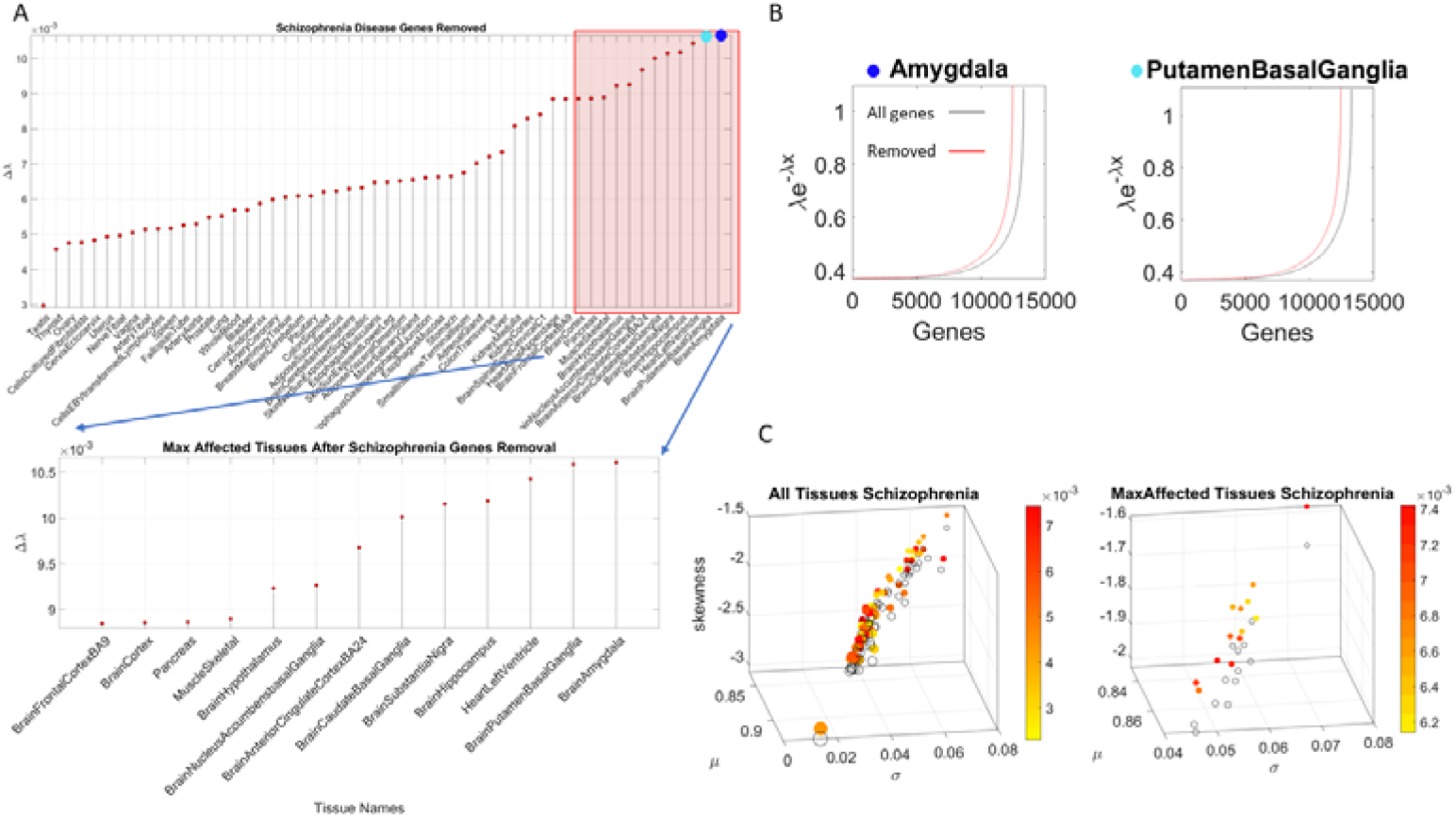
Sample metrics used for the stochastic analyses of a data sample (using 2,697schizophrenia associated genes reported in DisGenNet portal and in the literature.) (A) Effect of removing the schizophrenia genes from the GTex human genome set expressed across 54 tissues. Tissues are sorted in ascending order, by the absolute difference Δλ between genes’ expression on the 54 tissues before and after removal. Red square highlights the top 13 median ranked tissues shown in the panel below and dark and light blue circles mark the top two tissues affected (the brain amygdala and brain putamen in the basal ganglia. (B) The exponential distribution curve is fit to the sorted normalized count representing the gene’s expression in TMP on the top median-ranked affected tissues (as in Figure 2C, taking the peaks highlighted in red and fitting the exponential distribution to the frequency histogram, as in Figure 2D) before (black line) and after (red line) the removal of the genes associated with the disease. The absolute value difference between the curves is the Δλ used to rank the tissues by the effect size. (C) The fitting of the Gamma distribution yields the shape and scale parameters used to compute the Gamma moments. The axes represent the mean, the variance, and the skewness of the distribution of the normalized values and the color map represents the Earth’s Movers Distance values measuring the difference between the resulting exponential frequency histograms in (B). The size of the circle is proportional to the kurtosis and the color-filled circles represent the tissue (54 in the left panel) with the original genes’ expression from GTex (our reference template) *vs*. the open circles representing the stochastic shift, i.e. upon the removal of the genes associated with the disease in DisGenNet. Right panel contains the top ranked tissues (13) according to the median values of the Δλ.

### Stochastic Analyses – Visualization of change relative to the normative data of the full human genome

Using MLE we also obtain for each of the 54 tissues, the frequency histograms of the normalized counts across all genes and fit the continuous Gamma family of probability distributions with shape *(a)* and scale *(b)* values, to obtain the Gamma moments and plot them on a parameter space. We do this to visualize the spread of the tissues and their shift upon genes removal. To that end, we plot the mean, the variance, and the skewness across the x-, y- and z-axis respectively. We plot the size of the marker representing the tissue proportional to the kurtosis value, and we color the marker based on the change relative to the original genome count (*i.e*. containing all the genes, without removal.)

To measure the stochastic shift between the tissues from the full genome and those upon removal of the genes identified with each known neurological condition, we take the absolute difference between the MLE λ for the full GTex genome and that for the genome upon removal of the genes associated with each condition, disorder or disease, as shown in Figure 3.

We median rank the Δλ for each tissue, sorting Δλ in ascending order across the 54 tissues. Then we create 4 median ranked blocks and plot the maximally affected block of tissues (Figure 3A).

The highest ranked group is then compared across all conditions, mental illness *vs*. those neurologically defined *vs*. those non-neurological ones. We annotate the neurological functions that such tissues are known to maximally disrupt. And we ask if there is convergence between the tissue outcome in mental illnesses, upon removing the associated genes from the human genome-tissue model, and the outcome upon removing those genes tied to the other known disorders of the nervous systems. We repeat this interrogation process using the genes associated with disorders in the DisGeNet portal, including Autism, Schizophrenia, Depression, the neurological in Table 2, and the non-neurological ones in Table 3.

To assess tissue outcome upon removal of genes associated to various non-neurological diseases, we follow these procedures and compare these to the above results. These diseases include colon cancer, breast cancer, psoriasis, diabetes, congenital heart disease, hematologic neoplast and systemic lupus. Table 3 describes the number of genes associated with each of these diseases and the sources.

We also examine other mental illnesses described by the DSM. These include schizophrenia, depression, unipolar depression, and bipolar depression. We ask if there are tissues that overlap with those affected in the neurological disorders. Lastly, we examine mitochondria related disorders and PTSD using these methods. We reasoned that these may be disorders that potentially have affected tissues across a broader range of functions, including those from the brain and other bodily organs.

## 3. Results

### Autism, Ataxia and FX have convergence in maximally affected tissues by the removal of associated genes

The maximally affected tissues upon genes removal, according to the genes’ stochastic expression (count in Transcripts Per Million, TPM) are depicted in Table 4. This tissues’ gene expression was modelled by the exponential distribution *y* = *λe^−λx^*, with *x* as the genes’ combination expressed in the tissues, and λ as the exponential rate parameter. The Δ*λ* between the neurotypical template case from the GTex portal (containing all genes) and the modeled disorder case (upon removal of the SFARI genes) provides a sense for the departure from the normative case. This difference, taken for the removal of the SFARI genes, is depicted in Figure 4A, with samples of maximally affected tissues in Figure 4B that are known to be critical for motor control, regulation, adaptation/learning, and coordination.

**Figure 4.**
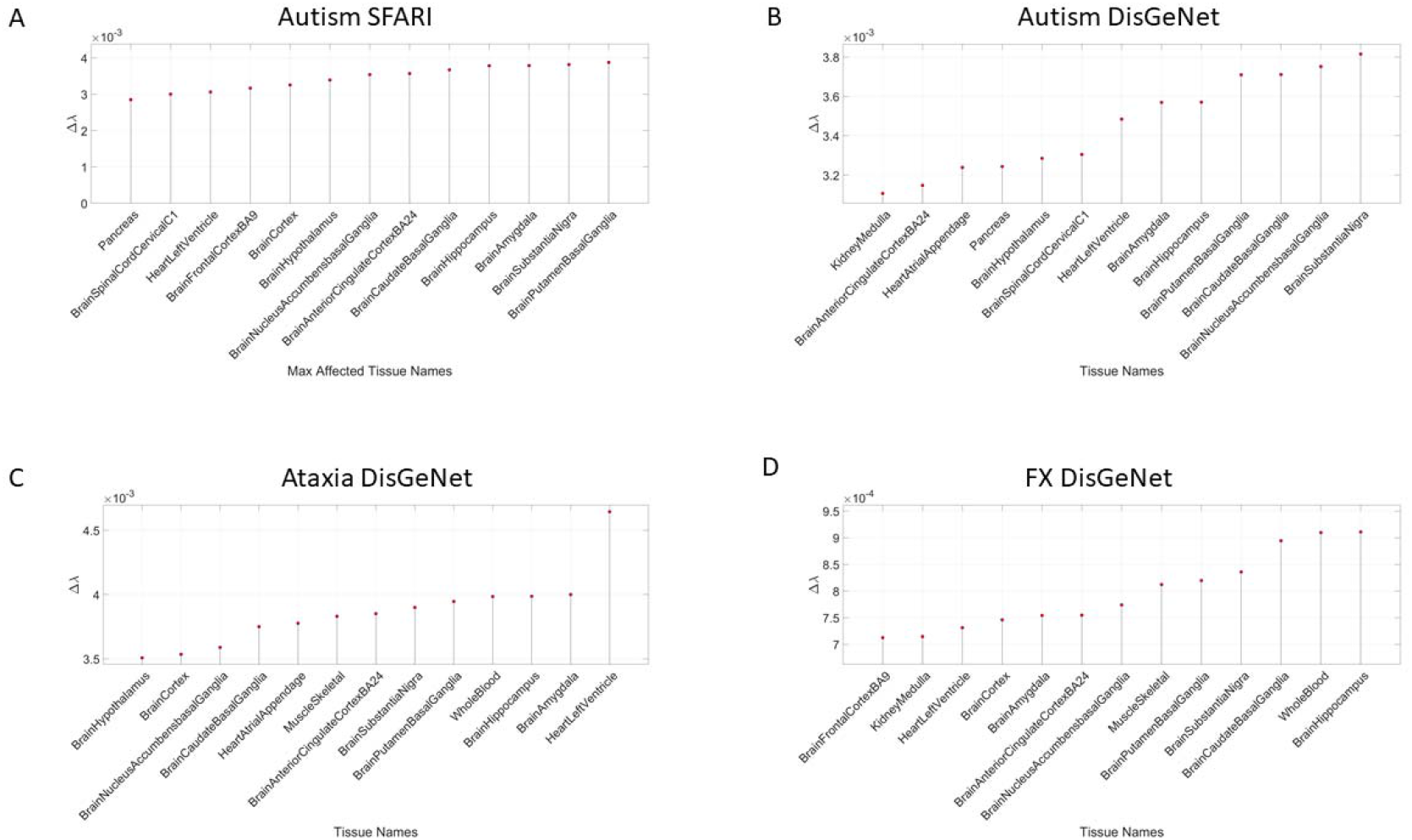
Convergence between Autism and known Neurological disorders shown by comparison of maximally affected tissues after removing genes associated with the disorder in Autism (from SFARI), Autistic Disorders (from the DisGeNet portal) *vs*. Ataxia and Fragile X (See Tables 4–5)

**Table 4.** The 13 top median ranked tissues in descending order of Δλ value, the highest affected tissues upon removal of the SFARI genes (906) linked to Autism from the human GTEx database in column 1. Ataxias, X-Chromosome, Fragile X, Parkinson’s disease and Mitochondrial disease extracted from the literature and column 3 is the same as in column 1 while removing from the SFARI Autism set 14 genes that overlap with the Ataxias and PD (see those genes listed in Supplementary Table 2). Tissues are grouped by CNS (brain and spinal cord in blue); muscle skeletal (green), heart (pink) and peripheral organs (gray). SFARI Autism (11/13 (84.6%) CNS, 1/13 (7.6%) heart and 1/13 (7.6%) peripheral organ); neurological disorders 10/13 (76.9%) CNS, 2/13 heart (15.3%) and 1/13 (7.6%) muscle skeletal); SFARI Autism without the overlapping genes from the neurological disorders 11/13 (76.9%) CNS, 1/13 (7.6%) heart and 1/13 (7.6%) peripheral organs.

| SFARI Autism | Ataxias, X, FX, PD, Mitochondria | SFARI Autism without overlapping Ataxia, PD genes |
| --- | --- | --- |
| Putamen Basal Ganglia | Hippocampus | Putamen Basal Ganglia |
| Substantia Nigra | Amygdala | Substantia Nigra |
| Amygdala | Substantia Nigra | Amygdala |
| Hippocampus | Heart Left Ventricle | Hippocampus |
| Caudate Basal Ganglia | Whole Blood | Caudate Basal Ganglia |
| Anterior Cingulate Cortex | Muscle Skeletal | Nucleus Accumbent Basal Ganglia |
| Nucleus Accumbent Basal Ganglia | Nucleus Accumbent Basal Ganglia | Anterior Cingulate Cortex |
| Hypothalamus | Putamen Basal Ganglia | Hypothalamus |
| Brain Cortex | Anterior Cingulate Cortex | Brain Cortex |
| Frontal Cortex | Brain Cortex | Frontal Cortex |
| Heart Left Ventricle | Hypothalamus | Spinal Cord |
| Spinal Cord | Heart Atrial Appendage | Heart Left Ventricle |
| Pancreas | Caudate Basal Ganglia | Kidney Cortex |

The Δλ-median ranking quantified the difference between neurotypical tissues’ gene expression *vs*. tissues’ gene expression upon removal of the genes corresponding to the disorders in question, with 4 groups ordered by the size of Δλ. This λ-quantity was first obtained relative to the neurotypical population tissues, *i.e*. including all the counts (gene expression) from all genes, *i.e*. to model an exponential process. The computation of λ using maximum likelihood estimation (MLE) is explained in the Appendix A. It does not assume any order of the counts, but rather seeks to identify the resulting λ for each tissue, treating the genes’ count (expression) as a random, memoryless stochastic process. Typically, the exponential distribution is used to model times between events, but here we used it to model the fluctuations in the values of the counts across the genes as they randomly fluctuate their expression across each of the 54 tissues reported in the GTex portal. We note this to underscore that the results spontaneously self-emerge from the random combination of the genes involved (with and without removal), rather than from the clinical criteria used to denote the genes’ relevance to Autism, or the evidence from the literature used to determine their association. There is in fact no scoring of such relevance for the genes associated to the other neurological disorders under consideration (*e.g*. the Ataxias, Parkinson’s, etc.) or for the Autistic disorders reported in the DisGenNet portal. Using those genes instead of the SFARI reveals tissues in Table 5, where we report the convergence.

**Table 5.** Highest affected tissues upon removal of the DisGeNet genes associated with Autistic Disorder (1,043) from the human GTEx database (column 1); Ataxia (813 genes) in DisGeNet (column 2) and FX (194 genes) in DisGeNet (column 3). Convergence between autism and neurological disorders is noted in the shaded tissues color coded as in Table 4, based on CNS, Heart, and peripheral organs.

| DisGeNet Autistic Disorder | DisGeNet Ataxia | FX DisGeNet |
| --- | --- | --- |
| Substantia Nigra | Heart Left Ventricle | Hippocampus |
| Nucleus Accumbent Basal Ganglia | Amygdala | Whole Blood |
| Caudate Basal Ganglia | Hippocampus | Caudate Basal Ganglia |
| Putamen Basal Ganglia | Whole Blood | Substantia Nigra |
| Hippocampus | Putamen Basal Ganglia | Putamen Basal Ganglia |
| Amygdala | Substantia Nigra | Muscle Skeletal |
| Heart Left Ventricle | Anterior Cingulate Cortex | Nucleus Accumbent Basal Ganglia |
| Spinal Cord | Muscle Skeletal | Anterior Cingulate Cortex |
| Hypothalamus | Heart Atrial Appendage | Amygdala |
| Pancreas | Caudate Basal Ganglia | Brain Cortex |
| Heart Atrial Appendage | Nucleus Accumbent Basal Ganglia | Heart Left Ventricle |
| Anterior Cingulate Cortex | Brain Cortex | Kidney Medulla |
| Kidney Medulla | Hypothalamus | Frontal Cortex |

Removal of SFARI genes, ranked by change in gene expression, reveals brain tissues linked to the CNS, in brain subcortical tissues linked to motor control (basal ganglia, striatum), memory (hypocampus), emotions (amygdala) and regulation (hypothalamus); and the spinal cord. This is also generally the case for the removal of the DisGeNet genes associated with Autistic Disorders and with FX and Ataxia. Congruent with the outcome from the removal of the SFARI genes from the GTex genome, the DisGeNet genes removal also affected the tissues associated CNS function. Important tissues for systemic organs’ functioning such as those containing smooth muscles, cardiac and skeletal muscles in the taxonomy proposed in Figure 1B were also affected (commonly) across these disorders. Figure 4 shows a summary of the results, visualizing the stochastic shifts. Tables 4–5 show the tissues ranked in descending order and color-coded according to CNS, brain, and spinal cord (blue); heart-related (pink), muscle skeletal (green) and peripheral vital organs (gray). Most tissues in Autism and the neurological disorders are from the CNS, followed by the PNS related tissues in the heart and muscle skeletal and with the vital organs towards the end of the Δλ-ranking.

Supplementary Figures 1-3 show these results separately for each neurological condition. We note in Supplementary Material that removal of the SFARI Autism syndromic genes from the GTex genome reveals maximal differences in tissues of organs with smooth and cardiac muscles, linked to involuntary and autonomic function in the proposed taxonomy of Figure 1B.

Removal of overlapping SFARI genes and Neurological disorders also reveal brain tissues linked to motor control, memory, emotions, and regulation. This is depicted in Figure 4. Given the congruence between the tissues maximally affected by removing the SFARI Autism genes from the GTEx database and those from the Neurological conditions, we next ascertain the extent to which these genes overlap with those used from the Ataxias in the literature. To that end, we divide them into the autosomal dominant, the autosomal recessive and the X-Chromosome genes. Figure 5A shows the result of this interrogation broken down by the Ataxia’s autosomal dominant, autosomal recessive and X-Chromosome genes. The genes listed in this figure overlap with those from the SFARI Autism genes. Table 4-Column 3 lists the tissues while Table 6 also has the scoring from the SFARI Autism genes. The Supplementary Material Table 2 lists the phenotypic information of the disorders associated with these genes, as described by the clinical literature. Figure 5C lists the PD gene that overlaps with the SFARI Autism genes, also depicted on Table 6 along with the score ranking from the SFARI portal.

**Figure 5.**
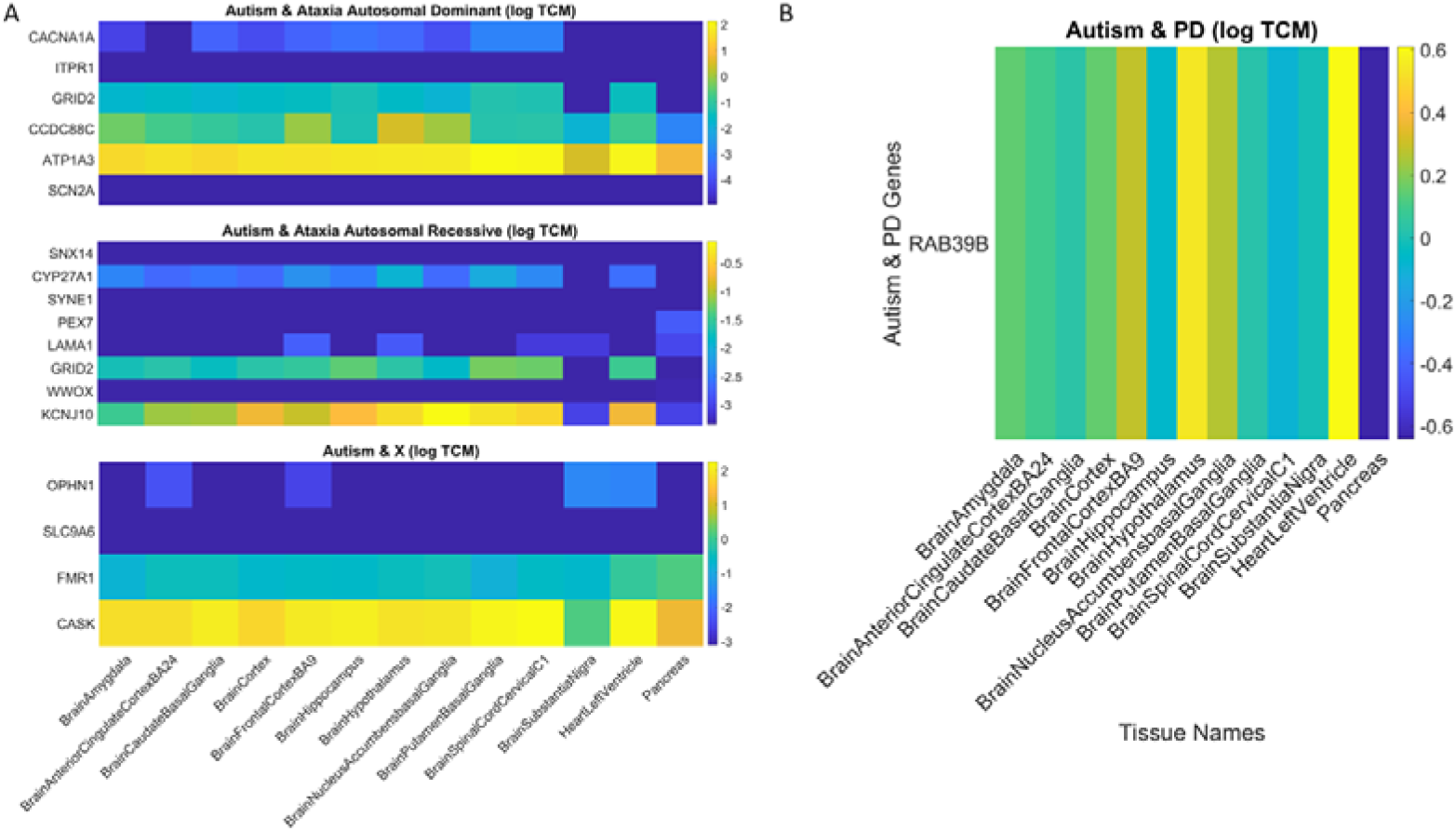
Genes’ expression on maximally affected tissues (colorbar coded in log TPM) upon removal of overlapping genes between the SFARI Autism set and the Ataxias (dominant and recessive and X-Chromosome sets) from the literature (A) and (B) from Parkinson’s disease. Horizontal axis is the tissues’ names and vertical axis the genes’ names.

**Table 6.**
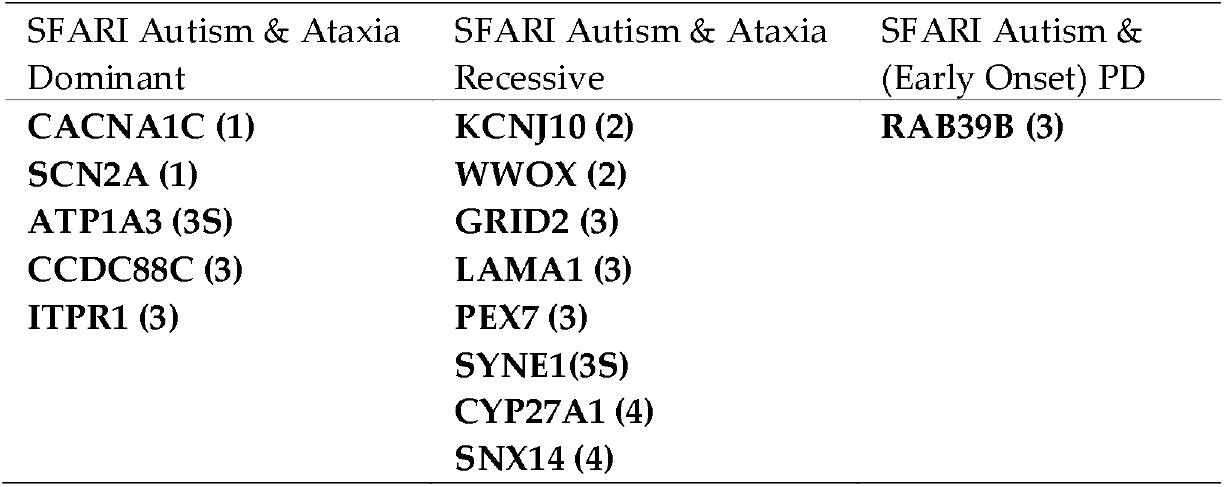
Overlapping genes between the Ataxias (dominant and recessive genes) and the Parkinson’s disease with the genes from the SFARI portal. Scores in parenthesis refer to the scoring of the gene according to the SFARI site (see Methods for explanation on each category). Syndromic is (4). Supplementary Figures 5-18 provide the GTEx violin plots of these genes’ expression in the top-ranked tissues unveiled by our analyses in Table 4. Supplementary Table 2 compiles the genes’ additional information from various sources in the clinical literature.

We note that removing this subset of 14 overlapping genes from the SFARI Autism set (Table 6), does not change the primary result whereby the most affected tissues upon removal of the SFARI Autism set from the GTEx dataset are those associated with sub-cortical brain structures critical for motor control, adaptation/learning, regulation, coordination and autonomic function, as well as memory and emotion. This is shown in Figure 5A-B and in the third column of Table 4. We also plot in Figure 4D the top ranked tissues affected by the removal from the GTEx dataset of the FX genes reported in the DisGeNet Portal. Supplementary Figures 19-22 further provide details on X-Chromosome genes in Figure 5A implicated in Autism according to the SFARI genes portal.

The results that convergence in the ranked descending order of CNS (brain and spinal cord tissues), followed by heart-related tissues, muscle skeletal tissue and lastly peripheral vital organs for systemic functioning in SFARI-Autism and well-known neurological disorders from the literature is also congruent with the results using the genes associated with these conditions in the DisGeNet portal. There we interrogated Autistic Disorders, Ataxias and Fragile X, confirming the overlapping in genes, their expression on the 54 tissues of the GTex database and the orderly levels of tissues maximally affected by the removal of the associated genes. We grouped the tissues by CNS, heart, muscle skeletal and peripheral vital organs to follow the proposed taxonomy of Figure 1B.

In the remaining of the paper, we consistently use this grouping to simplify visualization of the tables and data presentation. Figure 6 shows the results for different types of the Ataxias and FX, while Table 7 summarizes the top ranked affected tissues in different types of Ataxias, color-coded according to the grouping approximating the taxonomy proposed in Figure 1B.

**Figure 6.**
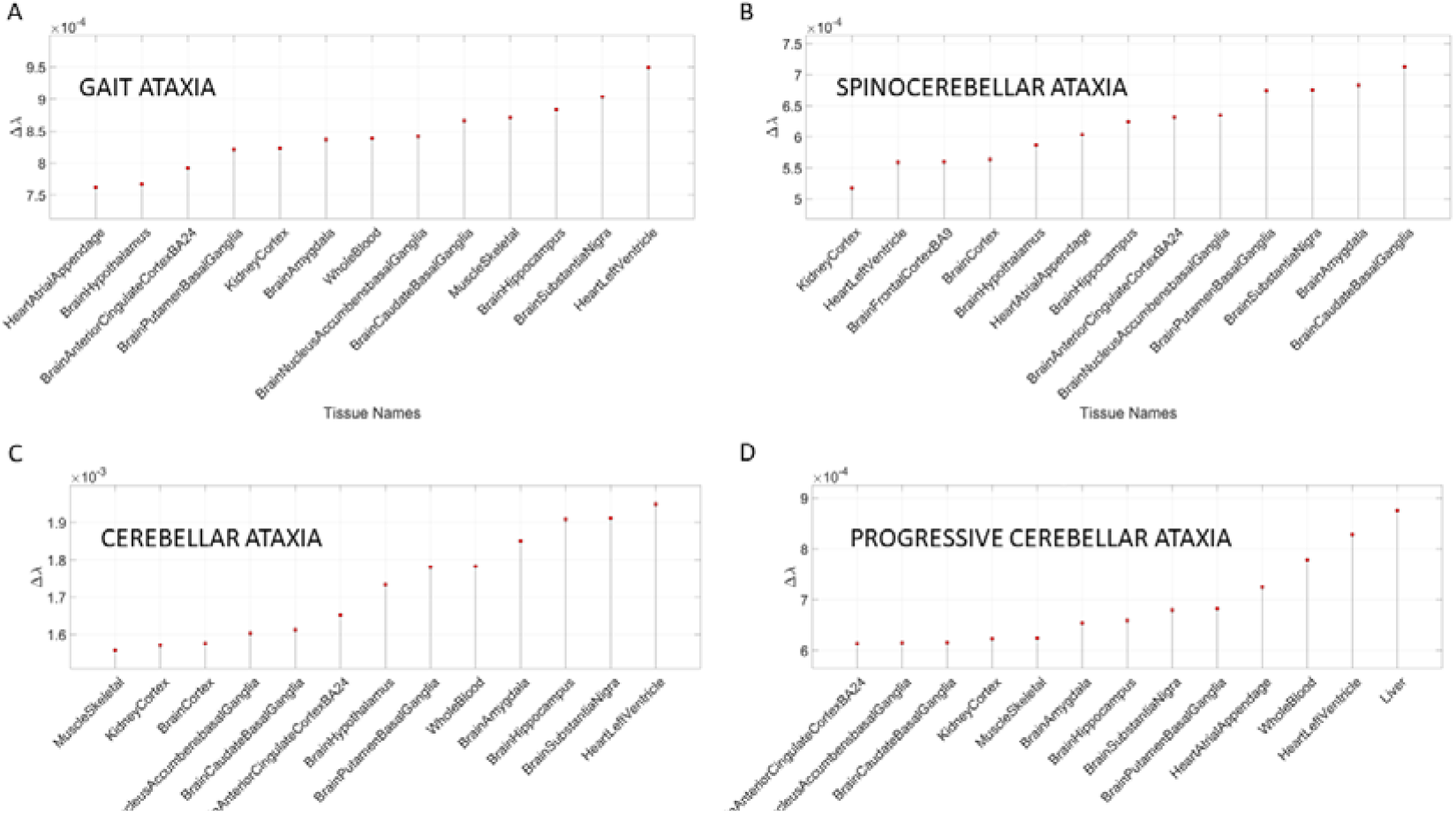
Different types of Ataxia showing the top Δλ-ranked tissues also shown in Table 7. (A) Gait Ataxia; (B) Spinocerebellar Ataxia; (C) Cerebellar Ataxia; (D) Progressive Cerebellar Ataxia.

**Table 7.** Highest affected tissues upon removal of the DisGeNet genes associated with different types of Ataxias, color-coded by CNS (brain, spinal cord), heart-related, muscle-skeletal, and peripheral vital organ for systemic functioning. Predominance of CNS is evident, followed by heart-related and muscle skeletal and peripheral organs.

| Gait Ataxia | Spinocerebellar Ataxia | Cerebellar Ataxia | Progressive-C Ataxia |
| --- | --- | --- | --- |
| Heart Left Ventricle | Caudate Basal Ganglia | Heart Left Ventricle | Liver |
| Substantia Nigra | Brain Amygdala | Substantia Nigra | Heart Left Ventricle |
| Hippocampus | Substantia Nigra | Hippocampus | Whole Blood |
| Muscle Skeletal | Putamen Basal Ganglia | Amygdala | Heart Atrial Appendage |
| Caudate Basal Ganglia | N Accumbens BG | Whole Blood | Putamen Basal Ganglia |
| N Accumbens BG | Ant Cingulate Cortex | Putamen Basal Ganglia | Substantia Nigra |
| Whole Blood | Hippocampus | Hypothalamus | Hippocampus |
| Brain Amygdala | Heart Atrial Appendage | Ant Cingulate Cortex | Brain Amygdala |
| Kidney Cortex | Hypothalamus | Caudate Basal Ganglia | Muscle Skeletal |
| Putamen Basal Ganglia | Brain Cortex | N Accumbens BG | Kidney Cortex |
| Ant Cingulate Cortex | Frontal Cortex | Brain Cortex | Caudate Basal Ganglia |
| Hypothalamus | Heart Left Ventricle | Kidney Cortex | N Accumbens BG |
| Heart Atrial Appendage | Kidney Cortex | Muscle Skeletal | Ant Cingulate Cortex |

### Removal of Genes Associated with Schizophrenia and Multiple Forms of Mental Depression Reveal Convergence with Neurological Disorders

The removal from the normative GTex genome of the DisGeNet genes associated with mental illnesses such as Schizophrenia, Depression, Bipolar Depression and Unipolar Depression resulted in convergence of maximally affected tissues involved in the CNS, especially those brain regions necessary for neuromotor control, memory, and emotion. This is depicted in Table 8 and Figure 7. Several of these tissues were also found to be affected upon removal of the SFARI genes and the genes associated in DisGeNet with Autistic Disorders. Furthermore, these are maximally affected tissues in the well-known neurological conditions depicted in Figures 4–5, Tables 4–6.

**Figure 7.**
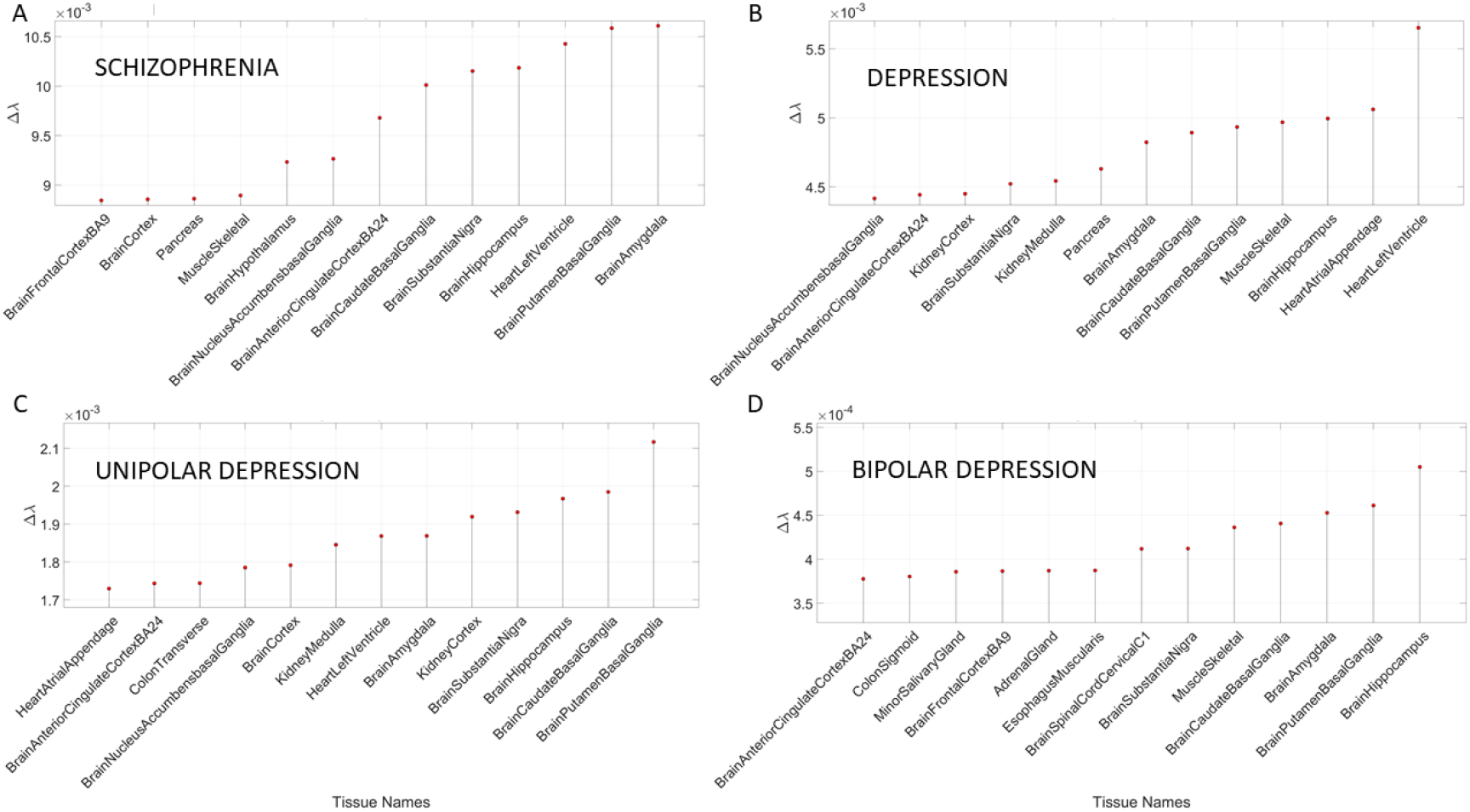
Maximally affected tissues in Schizophrenia and Depression, and in different types of Depression (Unipolar and Bipolar) shown in Table 8.

**Table 8.** Highest affected tissues upon removal of the DisGeNet genes associated with Schizophrenia (2,697) from the human GTEx database; Depression genes (1,468); Unipolar Depression genes (641) and Bipolar Depression genes (116). Convergence between Schizophrenia and Depression is high, with maximally affected CNS tissues (10/13), followed by heart-related and muscle skeletal. Unipolar and bipolar depression also show systemic effect of vital peripheral organs (N stands for Nucleus, Ant for Anterior, and BG for Basal Ganglia.)

| Schizophrenia | Depression | Unipolar Depression | Bipolar Depression |
| --- | --- | --- | --- |
| Amygdala | Heart Left Ventricle | Putamen Basal Ganglia | Hippocampus |
| Putamen Basal Ganglia | Heart Atrial Appendage | Caudate Basal Ganglia | Putamen Basal Ganglia |
| Heart Left Ventricle | Hippocampus | Hippocampus | Amygdala |
| Hippocampus | Muscle Skeletal | Substantia Nigra | Caudate Basal Ganglia |
| Substantia Nigra | Putamen Basal Ganglia | Kidney Cortex | Muscle Skeletal |
| Caudate Basal Ganglia | Caudate Basal Ganglia | Amygdala | Substantia Nigra |
| Ant Cingulate Cortex | Amygdala | Heart Left Ventricle | Spinal Cord |
| N Accumbens BG | Pancreas | Kidney Medulla | Esophagus Muscularis |
| Hypothalamus | Kidney Medulla | Brain Cortex | Adrenal Gland |
| Muscle Skeletal | Substantia Nigra | N Accumbens BG | Frontal Cortex |
| Pancreas | Kidney Cortex | Colon Transverse | Minor Salivary Gland |
| Brain Cortex | Ant Cingulate Cortex | Ant Cingulate Cortex | Colon Sigmoid |
| Frontal Cortex | N Accumbens BG | Heart Atrial Appendage | Ant Cingulate Cortex |

### Removal of Genes Associated with Non-Neurological Disorders Reveal Other non-CNS Tissues

In addition to the examination of mental illnesses and neurological disorders, we also interrogated the GTex genome upon removal of genes associated with various non-neurological disorders. These included various forms of cancers, inflammatory and auto-immune disorders and other tissues related to the heart, the circulatory and the endocrine systems. Tables 9–10 summarize the results of this interrogation and Figures 8–9 show the Δλ-ranking graphs.

**Figure 8.**
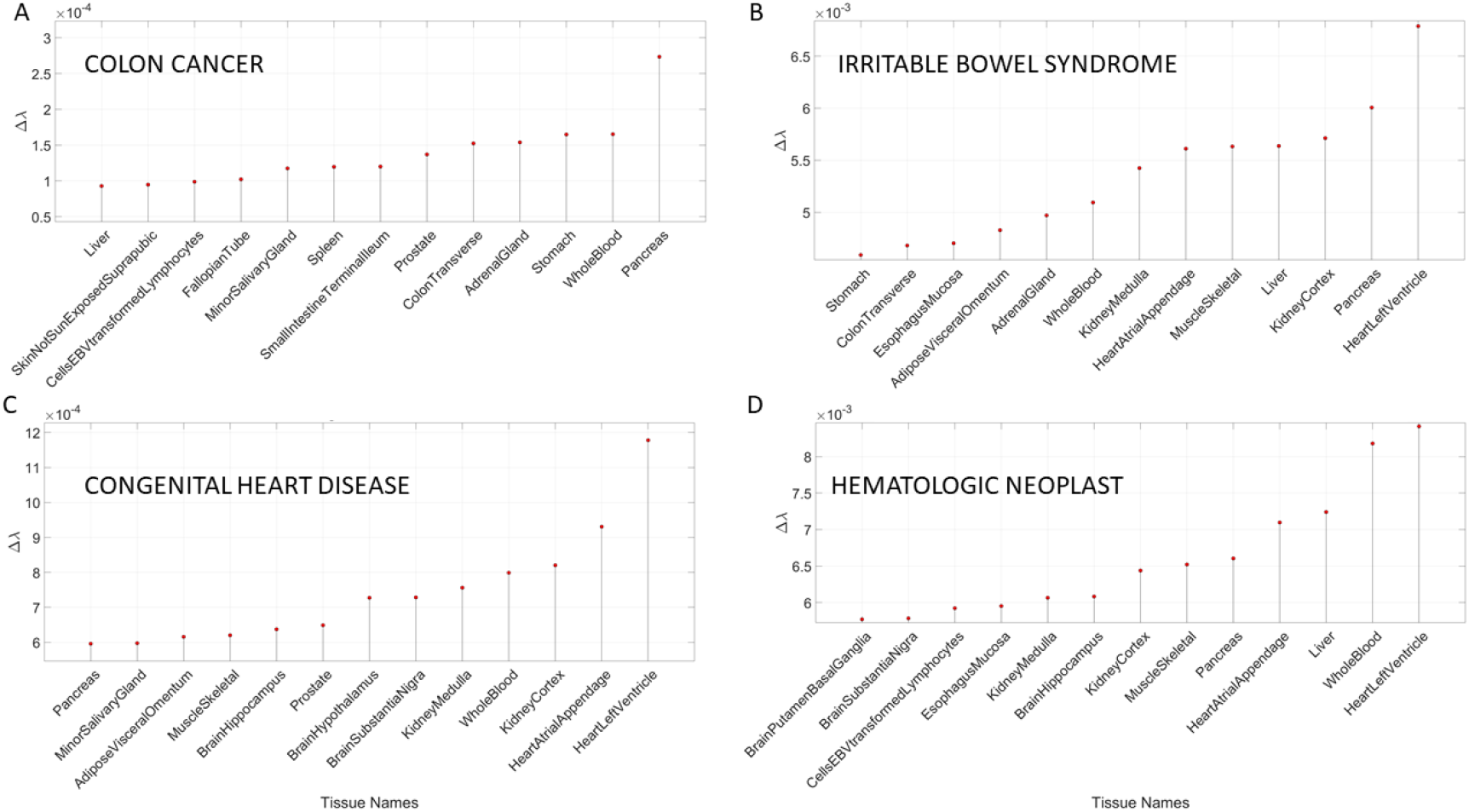
Maximally affected tissues in sample non-neurological diseases of Table 9 reveal primarily non-CNS tissues involving peripheral vital organs for systemic functioning, followed by heart-related and muscle skeletal tissues. As before, the interrogation of the GTex genome is based on the genes associated with diseases in the DisGeNet portal. (A) Colon cancer; (B) Irritable Bowel Syndrome; (C) Congenital heart disease and (D) Hematologic Neoplast. Color code as in previous Tables.

**Figure 9.**
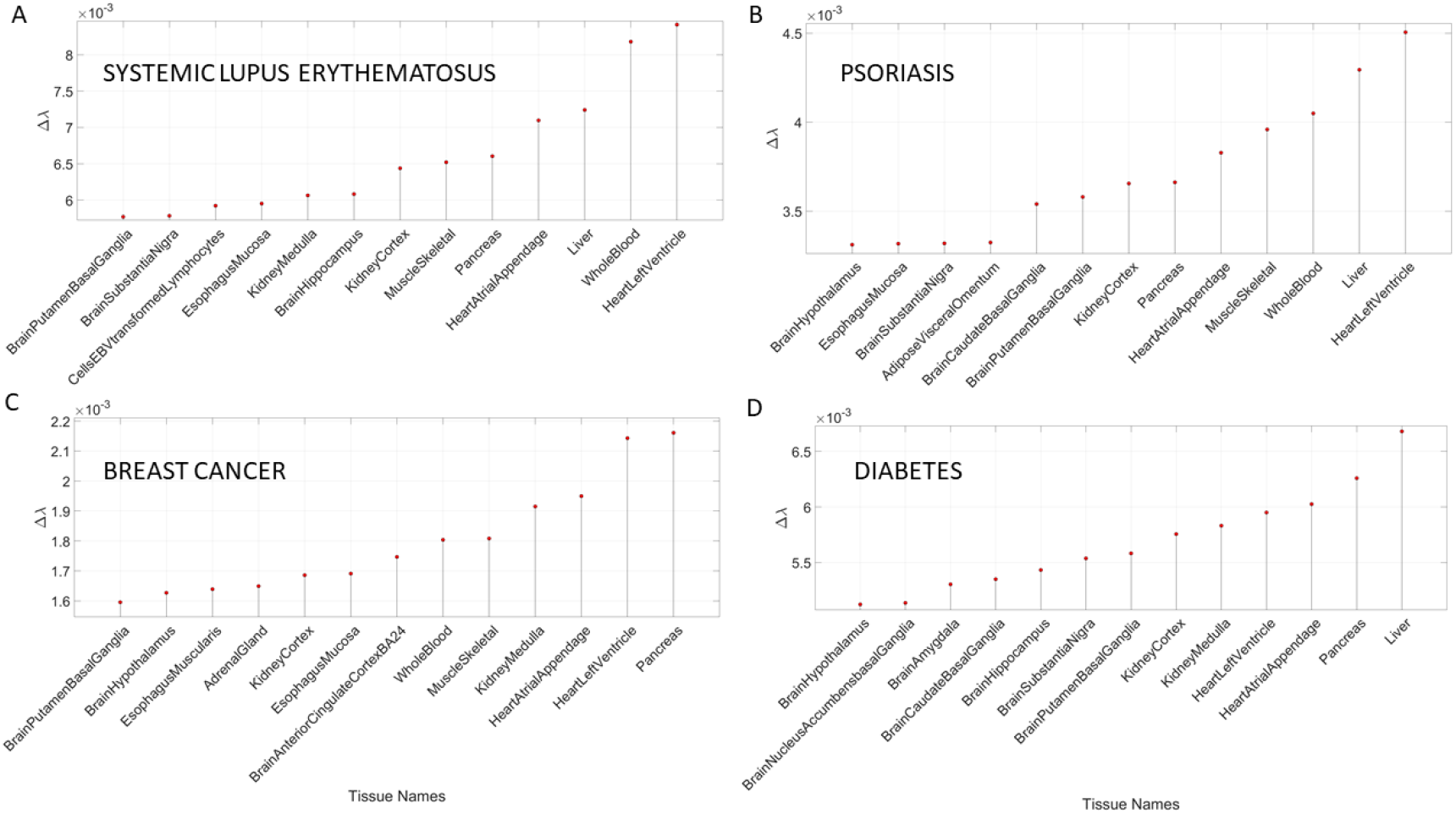
Maximally affected tissues in non-neurological diseases depicted in Table 10 median-ranked according to the Δλ-values obtained from the absolute difference between the tissues according to the full genome of the GTex database and the GTex genome without the genes associated to each disease, according to the queries to the DisGeNet portal.

**Table 9.** Highest affected tissues upon removal of the DisGeNet genes associated with Colon Cancer (3,669) from the human GTEx database; Irritable Bowel Syndrome genes (1,483); Congenital Heart Disease genes (267) and Hematologic Neoplast genes (827). Color code as in previous Tables.

| Colon Cancer | Irritable Bowel Syndrome | Congenital Heart Disease | Hematologic Neoplast |
| --- | --- | --- | --- |
| Pancreas | Heart Left Ventricle | Heart Left Ventricle | Heart Left Ventricle |
| Whole Blood | Pancreas | Heart Atrial Appendage | Whole Blood |
| Stomach | Kidney Cortex | Kidney Cortex | Liver |
| Adrenal Gland | Liver | Whole Blood | Heart Atrial Appendage |
| Colon Transverse | Muscle Skeletal | Kidney Medulla | Pancreas |
| Prostate | Heart Atrial Appendage | Substantia Nigra | Muscle Skeletal |
| Small Intestine T | Kidney Medulla | Hypothalamus | Kidney Cortex |
| Spleen | Whole Blood | Prostate | Hippocampus |
| Minor Salivary Gland | Adrenal Gland | Hippocampus | Kidney Medulla |
| Fallopian Tube | Adipose Visceral Oment | Muscle Skeletal | Esophagus Mucosa |
| EBT Lymphocytes | Esophagus Mucosa | Adipose Visceral O | EBT Lymphocytes |
| SkinNoSunExposed S | Colon Transverse | Minor Salivary Gland | Substantia Nigra |
| Liver | Stomach | Pancreas | Putamen Basal Ganglia |

**Table 10.** Highest affected tissues upon removal of the DisGeNet genes associated with Systemic Lupus Erythematosus (1,883) from the human GTEx database; Psoriasis genes (1,308); Breast Cancer (510) and Diabetes genes (3,134). The top half of the highest ranked tissues show no convergence with CNS-related tissues found in the mental illnesses and neurological disorders interrogated in this work. Instead, heart-related tissue, muscle skeletal tissue and tissues related to peripheral vital organs for systemic functioning are found. The bottom half of the top Δλ-ranked tissues are a mixture of tissues in peripheral bodily organs and brain-related tissues. The latter are from motor-control, coordination, and adaptation subcortical areas and from emotion, memory, and regulatory areas. Color code as in previous Tables.

| Systemic Lupus Erythematosus | Psoriasis | Breast Cancer | Diabetes |
| --- | --- | --- | --- |
| Heart Left Ventricle | Heart Left Ventricle | Pancreas | Liver |
| Whole Blood | Liver | Heart Left Ventricle | Pancreas |
| Liver | Whole Blood | Heart Atrial Appendage | Heart Atrial Appendage |
| Heart Atrial Appendage | Muscle Skeletal | Kidney Medulla | Heart Left Ventricle |
| Pancreas | Heart Atrial Appendage | Muscle Skeletal | Kidney Medulla |
| Muscle Skeletal | Pancreas | Whole Blood | Kidney Cortex |
| Kidney Cortex | Kidney Cortex | Ant Cingulate Cortex | Putamen Basal Ganglia |
| Hippocampus | Putamen Basal Ganglia | Esophagus Mucosa | Substantia Nigra |
| Kidney Medulla | Caudate Basal Ganglia | Kidney Cortex | Hippocampus |
| Esophagus Mucosa | Adipose Visceral O | Adrenal Gland | Caudate Basal Ganglia |
| EBT Lymphocytes | Substantia Nigra | Esophagus Muscularis | Amygdala |
| Substantia Nigra | Esophagus Mucosa | Hypothalamus | N Accumbens BG |
| Putamen Basal Ganglia | Hypothalamus | Putamen Basal Ganglia | Hypothalamus |

The results of the maximally affected tissues upon the removal of the genes associated with these non-neurological disorders revealed a very different picture than those upon removal of the genes associated with the mental illnesses (Autism, Schizophrenia and the Depressions) and those associated with the known neurological conditions (the various forms of Ataxia, FX and Parkinson’s disease.) Namely, the CNS-related tissues were less affected in these non-Neurological diseases than those related to the PNS (muscle skeletal and ANS heart) and those linked to peripheral bodily organs were the most visibly affected. The exception was Diabetes, maximally affected tissues in peripheral organs, but also CNS and PNS tissues in the tail of the top Δλ-ranked tissues. We next interrogate the genome in relation to mitochondrial disorders of several kinds and acquired PTSD.

### Removal of Genes Associated with Mitochondrial Diseases Reveal Heart-related Tissues are Maximally Affected but PTSD is mixed

**Table 11.** Highest affected tissues upon removal of the DisGeNet genes associated with Mitochondrial disease (284) from the human GTEx database; Mitochondrial Myopathies genes (121); Mitochondrial Encephalopathy, Lactic Acidosis, and Stroke-like episodes, MELAS (81) and Post-Traumatic Stress Disorder genes (418).

| Mitochondrial Disease | Mitochondrial Myopathies | MELAS | PTSD |
| --- | --- | --- | --- |
| Heart Left Ventricle | Liver | Heart Left Ventricle | Caudate Basal Ganglia |
| Muscle Skeletal | Heart Left Ventricle | Liver | Kidney Medulla |
| Pancreas | Muscle Skeletal | Kidney Cortex | Kidney Cortex |
| Heart Atrial Appendage | Hippocampus | Pancreas | Pancreas |
| Kidney Cortex | Whole Blood | Heart Atrial Appendage | Putamen Basal Ganglia |
| Whole Blood | Ant Cingulate Cortex | Kidney Medulla | Heart Left Ventricle |
| Kidney Medulla | Brain Amygdala | Putamen Basal Ganglia | Hypothalamus |
| Ant Cingulate Cortex | Substantia Nigra | Esophagus Mucosa | Substantia Nigra |
| Putamen Basal Ganglia | Heart Atrial Appendage | Brain Amygdala | Heart Atrial Appendage |
| N Accumbens BG | Putamen Basal Ganglia | Adipose Visceral Omen | N Accumbens BG |
| Adrenal Gland | Spinal Cord | Artery Coronary | Hippocampus |
| Hippocampus | Pancreas | Stomach | Colon Sigmoid |
| Caudate Basal Ganglia | Hypothalamus | Colon Traverse | Amygdala |

Removal of genes associated with Mitochondrial Disorders of various types from the GTex genome, according to the genes in the DisGeNet portal, reveal a mixture of tissues associated with peripheral vital organs for systemic functions, heart-related and muscle skeletal tissues and CNS-related tissues. The top half of the highest ranked tissues in Mitochondrial Disease shows affected tissues related to the heart, muscle skeletal and peripheral organs, while the bottom half shows more involvement of brain-related tissues in sub-cortical regions of motor control. In contrast, Mitochondrial Myopathies show a predominance of CNS-related tissues, including the brain and spinal cord, with top Δλ-ranked tissues related to the heart and muscle skeletal tissues. MELAS shows a predominance of tissues associated with peripheral vital organs for systemic function and heart-related tissues. Only two brain regions for motor control and emotion are present in the bottom ranked tissues of the topmost affected tissues.

The case of acquired PTSD also reveals a mixture of tissues from brain, heart, and peripheral organs. There we see maximally affected tissues linked to sub-cortical regions of the brain involved in motor control, adaptation, learning, and coordination intermixed with tissues linked to peripheral bodily organs (like the kidneys) and the autonomic systems’ heart. Furthermore, we see also tissues linked to the hypothalamus, a regulatory brain structure. Figure 10 shows the graphs of the Δλ-difference median ranked as in the previous cases, for these disorders.

**Figure 10.**
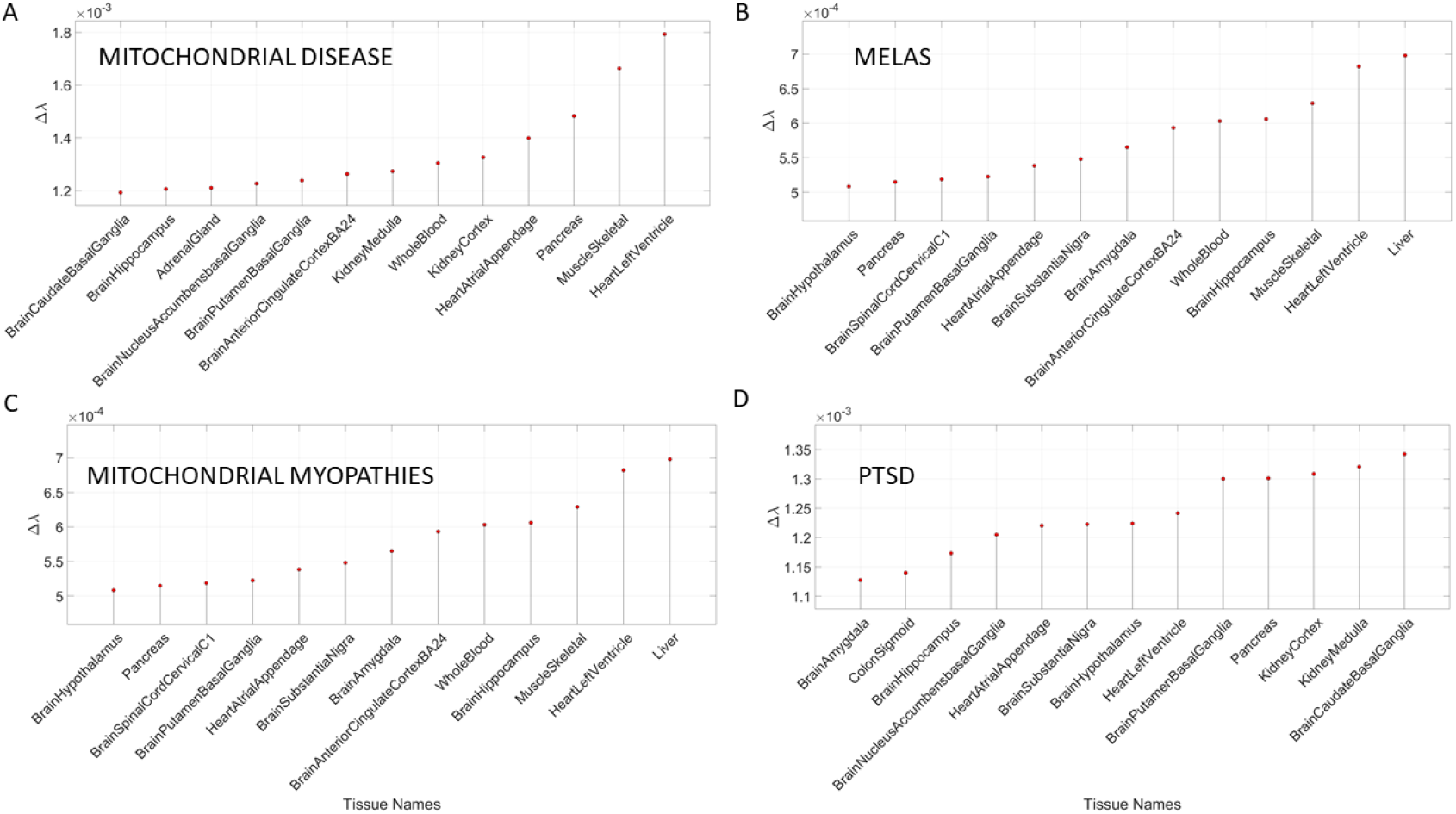
Maximally affected tissues in Mitochondrial Diseases and in PTSD of Table 11, shown in graphical form according to the Δλ values.

We summarize the results across all 54 tissues (in alphabetical order from left to right) in Figure 11. Here a color map depicts the values of Δλ normalized for each disease (along the rows) across the tissues (columns) by dividing by the maximum Δλ value of each row. The pattens reveal that neurological disorders and mental illnesses shared the tissues related to the brain as the maximally affected. They also reveal that the whole blood tissue is not as affected in the mental illnesses as in the neurological disorders (marking a point of divergence that warrants further investigation). Heart-related tissues and muscle skeletal are also shared between these mental illnesses and neurological disorders when the genes specific to each disorder are removed from the GTex genome. Interestingly, the pancreas shows as a tissue from a peripheral bodily organ commonly affected across most disorders and diseases interrogated here (with lower value in the neurological ones.)

**Figure 11.**
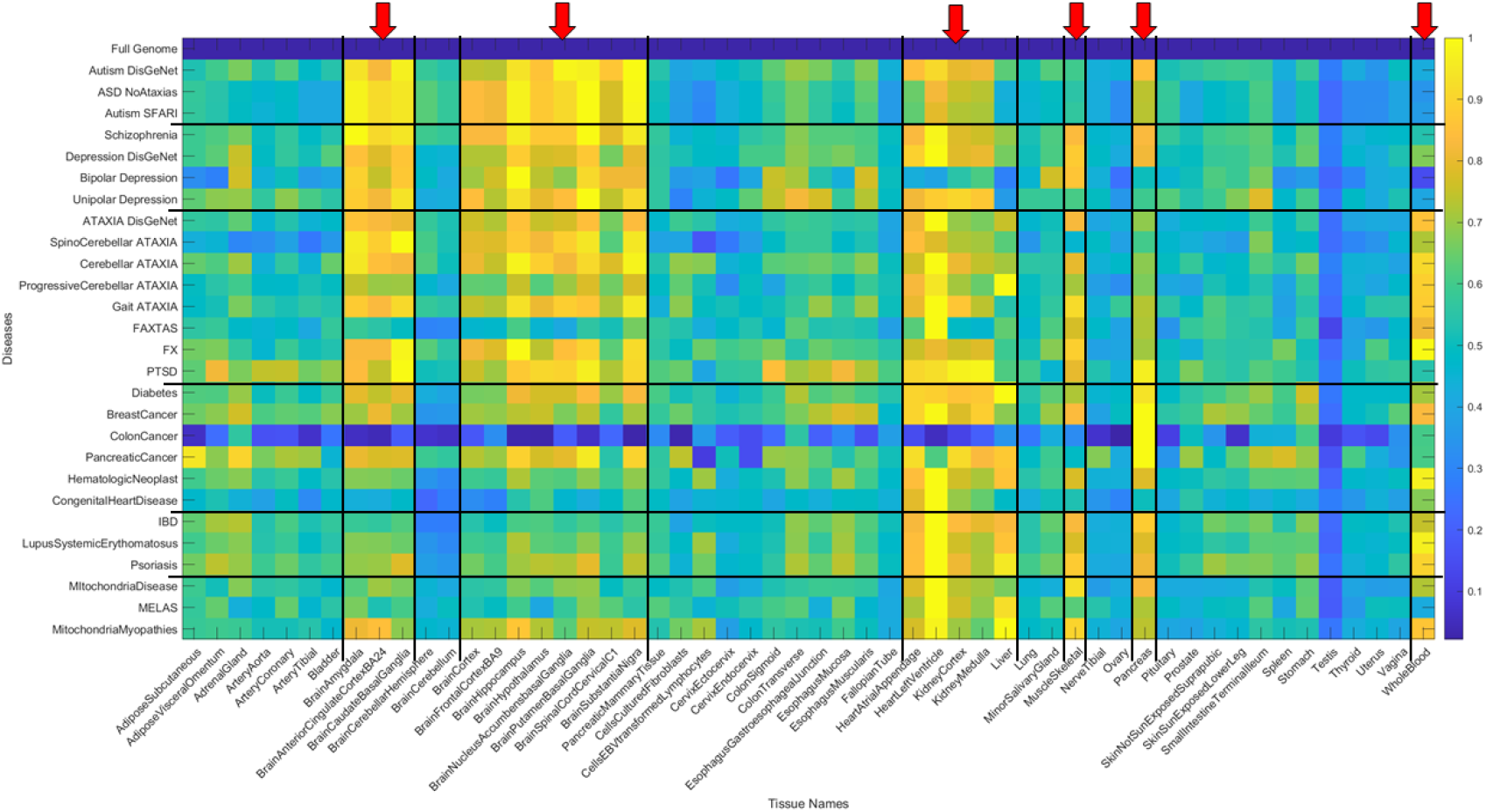
Summary of disorders/diseases (27 rows) x tissues (54 columns) in alphabetical order. Entries are Δλ (difference with respect to the genes’ expression values from the full Gtex genome), values normalized by the maximum across the tissues for each row (disorder/disease.) First row is the 0-Δλ difference reference from full genome. Red arrows mark the maximally affected tissues across all diseases, showing mental illnesses on the top, followed by neurological disorders, including several types of cancer, auto-immune disorders, diabetes, etc. Black lines delineate blocks of diseases (rows) and blocks of genes’ expression on tissues (columns.)

The non-neurological diseases reveal less involvement of the CNS-related tissues but highly overlapping with the heart and muscle skeletal tissues. Tissues linked to the kidneys, liver and pancreas are also maximally affected in these diseases, with colon cancer showing an interesting pattern whereby the pancreas reveals top normalized Δλ value. Figure 12 summarizes the patterns in binary by turning ON (yellow) values above.8 considered high and OFF those below (blue). This cutoff is chosen to further highlight overlapping and differences across diseases based on high Δλ.

**Figure 12.**
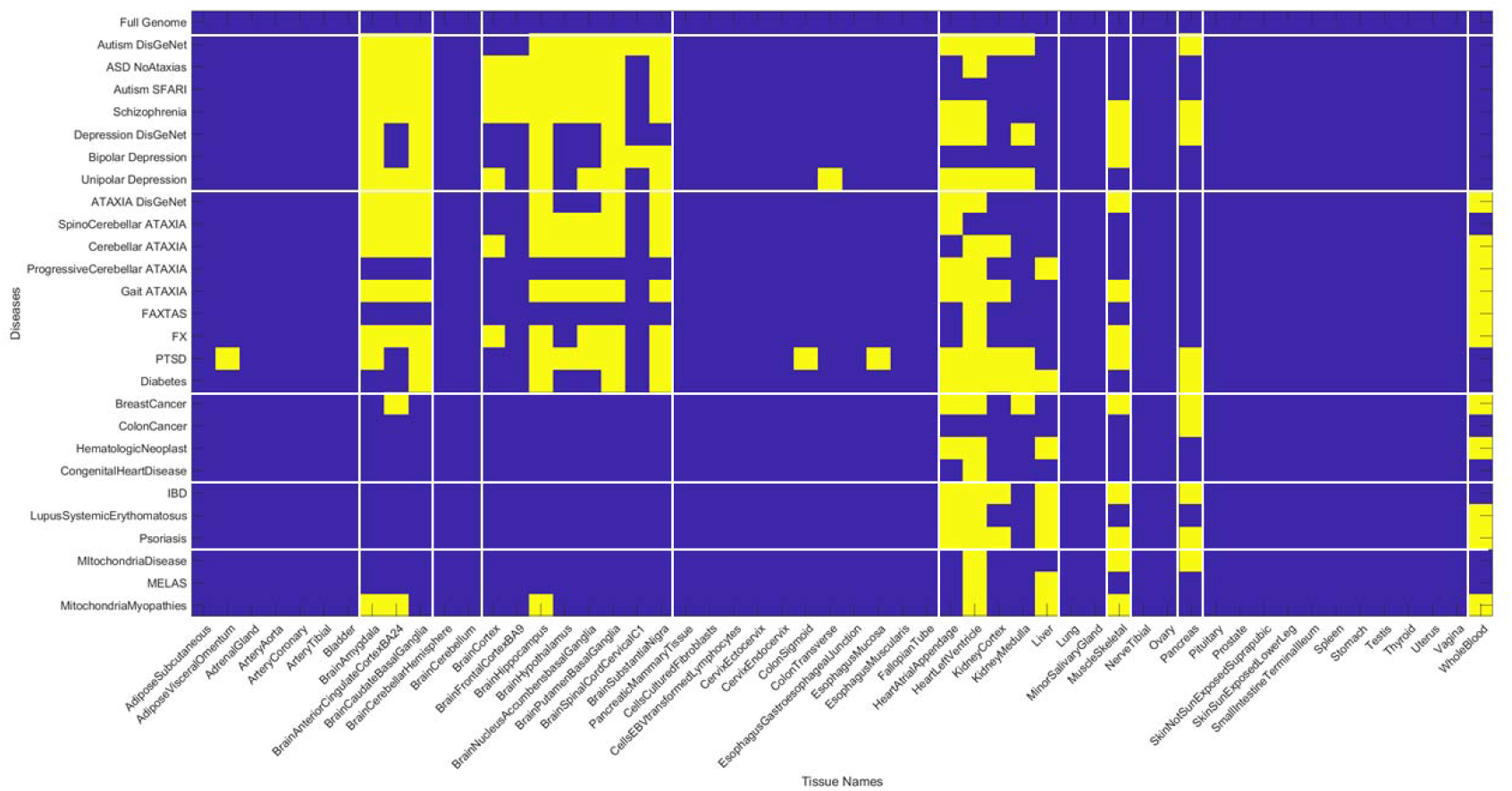
Binary version of matrix in Figure 11 upon thresholding by high normalized Δλ value of .8 shows the overlap across mental illnesses, neurological disorders and non-neurological diseases are primarily in the heart tissues, the muscle skeletal and organs like the pancreas, liver, and kidney. Whole blood tissue is shared between neurological and non-neurological disorders but not present in the mental illnesses. Brain tissues are shared between mental illnesses and neurological disorders.

The patterns revealed by the normalized Δλ high values show convergence of maximally affected brain tissues in mental illnesses with the neurological disorders but not with the non-neurological ones (except for diabetes which does affect some brain regions). The Mitochondrial diseases do not show the same intensity of the CNS-related Δλ values as the mental illnesses and the neurological disorders do, but they do share the heart and muscle skeletal patterns with all the examined diseases and disorders. This is interesting, given that some of the children with various forms of mitochondrial disorders may receive diagnoses of autism. In summary, there is clear overlap between mental illness and neurological disorders suggesting involvement of the central nervous systems in both, but also contributions from the peripheral nervous systems, particularly the heart, the muscle skeletal tissues and to a lesser degree, tissues of peripheral organs. The latter are most affected in the non-neurological diseases. Figure 13 shows the most affected tissue in each disease/disorder (also depicted on Appendix B Table.)

**Figure 13.**
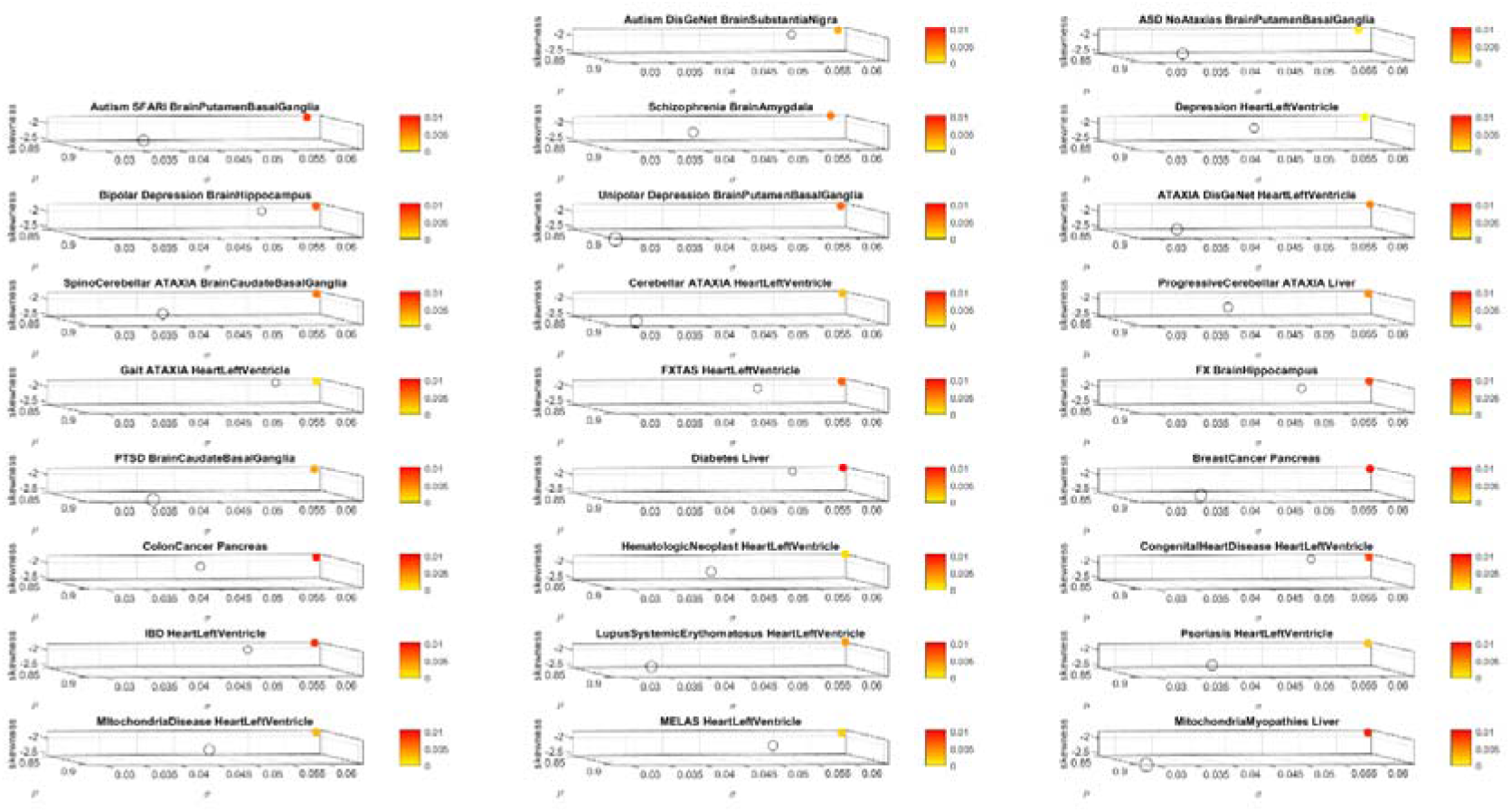
Stochastic shift of genes expression on tissue, after genes removal for each disease/disorder under consideration (colored circle) relative to the genes expression on that tissue using the full genome in GTex (open circles). The top affected tissue in each disorder/disease under consideration was selected according to the maximum Δλ value, the absolute difference between the empirically estimated λ using MLE of the exponential distribution rate parameter for the full GTex genome and for the genome minus the genes associated to each disease/disorder. Since the exponential distribution is a particular case of the continuous Gamma family of probability distributions (when the Gamma shape parameter is 1), we also used MLE to estimate the shape and scale Gamma parameters and the 4 Gamma moments, plotted here in a 5 dimensional parameter space. Along the x-axis we plot the empirically estimated Gamma mean; along the y-axis we plot the Gamma variance; along the z-axis we plot the Gamma skewness and the kurtosis is used to represent the size of the marker (more kurtotic distributions have higher value, i.e. larger circle.) The fifth dimension is the color representing the Δλ (see also Methods Figure 3C visualizing one single disease and all 54 tissues or summarizing the top 13 median-ranked tissues as those most affected).

## 4. Discussion

This work interrogated the human genome by removing genes associated with various diseases and comparing the outcome from the remaining genes’ expression on 54 tissues commonly examined in the GTex portal. These tissues involve parts of the central nervous systems (the brain and the spinal cord), parts of the peripheral nervous systems (the muscle skeletal tissue and the heart tissues as part of the autonomic nervous system) which we grouped as in the Figure 1B. Other tissues are from peripheral vital organs for systemic bodily functions (whole blood, pancreas, liver, kidneys, lungs, etc.)

We compared mental illnesses such as Autism, Schizophrenia, and various types of Mental Depression (including Unipolar and Bipolar), with well-known neurological disorders such as different types of Ataxias, Fragile X and Parkinson’s Disease. We found convergence between the tissues maximally affected by the removal of disease-associated genes across these mental illnesses and neurological disorders. CNS-related tissues in subcortical regions of the brain related to motor control, motor learning, motor coordination, and motor adaptation, as well as memory and emotion, were predominantly maximally affected by the corresponding genes’ removal across both the mental illnesses and the neurological disorders. This convergence demonstrates overlap between Psychiatric and Neurological conditions with specific involvement of motor, memory, and emotional axes. In Autism and FX, we obtained congruent results on maximally affected tissues both using for removal the genes from the SFARI-Autism database, and the genes upon querying the DisGeNet portal. In addition to the genes reported by querying DisGeNet, we also used the genes reported in the literature for Schizophrenia and Depression, and for Parkinson’s disease. We found congruence in all cases.

To further test our hypothesis that these mental illnesses are *disorders of the nervous systems* and that removing the gene pool associated with them gives rise to overlapping tissues related to the CNS functioning, we also queried DisGeNet about other non-neurological diseases. We found that in such cases, the predominance of maximally affected tissues was on tissues associated with peripheral vital bodily organs related to the disease, as pancreas, kidney, liver, and colon transverse in colon cancer. Furthermore, several of these diseases had maximally affected heart-related tissues and whole blood. Other cases also revealed a predominance of peripheral organs. We lastly, interrogated the genome in relation to Mitochondrial diseases and to acquired PTSD. In these cases, we hypothesized and confirmed a mixture of tissues related to peripheral organs (for mitochondrial diseases) and to the CNS (for PTSD.)

In the case of Mitochondrial diseases, the heart-related tissues were revealed as the most affected along with muscle-skeletal tissue. Furthermore CNS-related tissues were more affected by the genes’ removal in Mitochondrial Myopathies (i.e. the amygdala and the anterior cingulate cortex), as compared to MELAS or to Mitochondrial disease. The common thread across all three types of Mitochondrial related disorders was the heart-related tissues. The case of acquired PTSD showed a mixture of CNS-related tissues, tissues related to bodily peripheral organs, and heart-related tissues. The kidney and pancreas were also affected in PTSD. When we examined the maximum Δλ for each disorder/disease under examination, we found that the basal ganglia was maximally affected in Autism, Unipolar Depression, Spinocerebellar Ataxia, and PTSD), while the heart left ventricle tissue was maximally affected in Depression, Ataxia, Cerebellar Ataxia, Gait Ataxia and Fragile X Tremor Ataxia Syndrome (FXTAS). This result indicates overlap between the Psychiatric mental illnesses and the Neurological disorders. It also shows the importance of examining mental illnesses in a more systemic way that includes the Autonomic Nervous Systems of the PNS.

This test on non-neurological illnesses served as a control to show that removal of the genes associated with each disorder did have specificity with the disorder, and yet, very different outcome when comparing the mental illnesses to the non-neurological disorders. Among the top tissues affected across non-neurological diseases, the pancreas was maximally affected by the removal of disease-associated genes in Breast and Colon Cancer, while the liver was maximally affected in Diabetes. The heart left ventricle was maximally affected across autoimmune disorders like psoriasis, Lupus Systemic Erythematosus, and Irritable Bowel Disease (IBD). The heart was also maximally affected in Hematologic Neoplast and Congenital Heart Disease, Mitochondrial Disease and MELAS, in contrast to the Mitochondrial Myopathies which showed the liver as the maximally affected tissue.

This exercise demonstrated that despite the stochastic nature of genes’ expression, upon their removal and random recombination, there is convergence across Psychiatric and Neurological disorders, thus potentially rendering both as *disorders of the nervous systems*. We found in both cases a strong prevalence of the CNS, but also found important differences in tissues from the PNS, including the heart and the muscle skeletal tissues involved in both mental illnesses and neurological disorders. Because of these convergences, and the fact that there are treatments and accommodations to help persons with neurological disorders, it may be possible to leverage some of those types of bodily-based supports to help persons with mental illnesses. Behaviors that are described by observation to define mental illnesses can now be connected with underlying tissues involved in voluntary, involuntary, and autonomic function across the CNS and the PNS and mapped to the genome, thus closing the present gap between behaviors and genomics in the Precision Medicine knowledge network. In this sense, the present methods offer a new way to interrogate the genome and link tissues with behavioral phenotyping.

A surprising finding here is the potential contributions of peripheral structures and organs to mental illness. Tissues of the autonomic nervous systems were maximally impacted by the removal of the genes associated with these mental illnesses, as was the muscle skeletal tissue among the top ranked ones. Tissues associated to sub-cortical brain regions necessary for motor control, learning, adaptation, and coordination (basal ganglia and striatum) were highly impacted by the removal of the genes in both mental illnesses and neurological disorders, along with those tissues important for memory (hippocampus), emotion (amygdala) and regulation (hypothalamus). Surprisingly, we did not see cerebellum-related tissues among the top affected by the removal of the genes (even in the ataxias) where we do know that the cerebellum plays a large role [21–24]. This was also the case in autism, where we know the cerebellum has been implicated [25,26].

Lastly, the autoimmune disorders that we examined had very different brain-tissue patterns from the mental illnesses and neurological conditions but shared the heart-related tissues and the muscle skeletal tissue. In this sense, the contributions from the peripheral systems to mental illnesses and to auto-immune disorders seems important. While blood tissue marked a departure of neurological disorders from mental illnesses, as it was maximally affected in neurological disorders but not in the mental illnesses. All and all these genes’ removal revealed surprising results that invite rethinking how we may want to describe, diagnose, and treat mental illnesses in general.

### Caveats and Future Directions

Although we found evidence that the mental illnesses and neurological disorders have remarkable overlap in the types of brain tissues that are maximally affected by the removal of their corresponding associated genes, we recognize that genes’ removal is a crude way to interrogate the human genome and its expression of the 54 tissues of the GTex database. Future work will aim at developing more sophisticated methods to explore genes’ over expression too, and to build simulations of the use of these methods in *e.g*. combination with transcriptome evolution during neuronal differentiation in the development of cell lines from induced pluripotent stem cells. This will be important to assess asynchronous genes’ expression behaviors over time when cell lines differentiate into neuronal types. Full transcriptome dynamic interrogation longitudinally, over time is now possible using these stochastic analyses in combination with the various data repositories featuring disease-associated genes.

The present work merely scratches the surface on possible new ways to interrogate the human genome in relation to diseases of all types (not just mental or neurological), to possibly build comparative models of outcomes *in tissues* that can be related to behavioral phenotypic manifestations of the clinical disorder. In this sense, the work presented here can help bridge the gap between behavioral description of a mental illness, or a neurological disorder and its genetic underpinnings via the affected tissues. Combining this approach with the new wave of digital biomarkers that describe human behavior digitally at a microscopic level [4, 27–30], using objective means and a finer level of granularity beyond naked eye detection, could help us redefine many Psychiatric disorders and medical conditions under the Precision Medicine paradigm.

## 5. Conclusions

We here offer a new roadmap to reframe Psychiatry using the Precision Medicine paradigm. The new stochastic approach can initiate the steps to connect behavioral phenotypic description from clinical observation, with the underlying neurobiology of mental illnesses. Borrowing knowledge from Neurology and Brain Science, it will be possible to shift Psychiatry from an art to a quantitative objective science under the tenets of Precision Medicine, by integrating all layers of the knowledge network. This would help design personalized targeted treatments utilizing the person’s genome, localizing the most affected tissues defining central nervous systems functions and distinguishing those from tissues related to vital organs for systemic functions. This new approach could potentially mark the beginning of a transformative era in Mental Health.

## 6. Patents

EBT holds the US Patent “Methods and Systems for the Diagnoses and Treatments of Nervous Systems Disorders” combined in the paper as micro-movement spikes, MMS data type and continuous Gamma probability distributions family empirical estimation.

## Supporting information

Supplementary Material - Reframing Psychiatry for PM

## Supplementary Materials

The following are available online at www.mdpi.com/xxx/s1, Supplementary Materials File.

## Author Contributions

Conceptualization, methodology, software, validation, formal analysis, investigation, resources, data curation, writing EBT.

## Funding

This research was funded by the New Jersey Governor’s Council for the Medical Research and Treatments of Autism NJDOH - CAUT17BSP024 and by the generosity of the Nancy Lurie Marks Family Foundation, Career Development Award to EBT.

## Acknowledgments

I thank the SFARI researchers for the curation and maintenance of their genes module and the compilation of literature database supporting the repository.

## Conflicts of Interest

The authors declare no conflict of interest. The funders had no role in the design of the study; in the collection, analyses, or interpretation of data; in the writing of the manuscript, or in the decision to publish the results.

## Appendix A

We estimate the likelihood *L*(*λ* | *x*_1_, *x*_2_,…,*x_n_*) where *x_i_* is the series of counts representing the gene expression on each given tissue and *i* ranges from *1* to *n*, the number of genes.

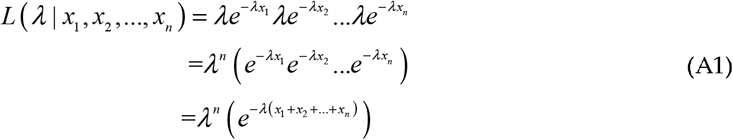

To obtain the maximum likelihood, we take the derivative of the likelihood in equation (A1) and set it to 0 (since the derivative is 0 at the maximum likelihood value).

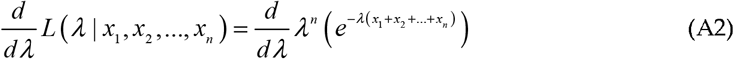

We take the *log* here because the derivative of the function and the derivative of the *log* of the function equals *0* at the same point, so for the purposes of finding where the derivative is 0, the original function in equation (A2) and the *log* of it are interchangeable.

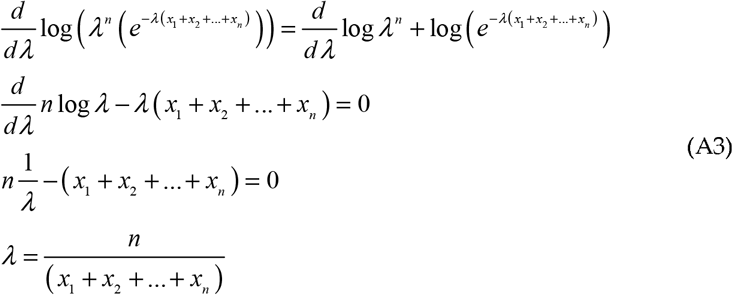

And with this result in equation (A3), we can obtain the maximum likelihood estimate of each λj, given all the 56,146 genes expressed with some random value for each of the *j* = 1:54 tissues.

## Appendix B

The data file name from the GTEx Portal https://www.gtexportal.org/home/datasets used in this paper is GTEx_Analysis_2017-06-05_v8_RNASeQCv1.1.9_gene_median_tpm.gct.csv

The data file name from SFARI Gene is located at https://gene.sfari.org/database/human-gene/ and named SFARI-Gene_genes_03-04-2020release_03-05-2020export.csv

**Appendix B Table:**
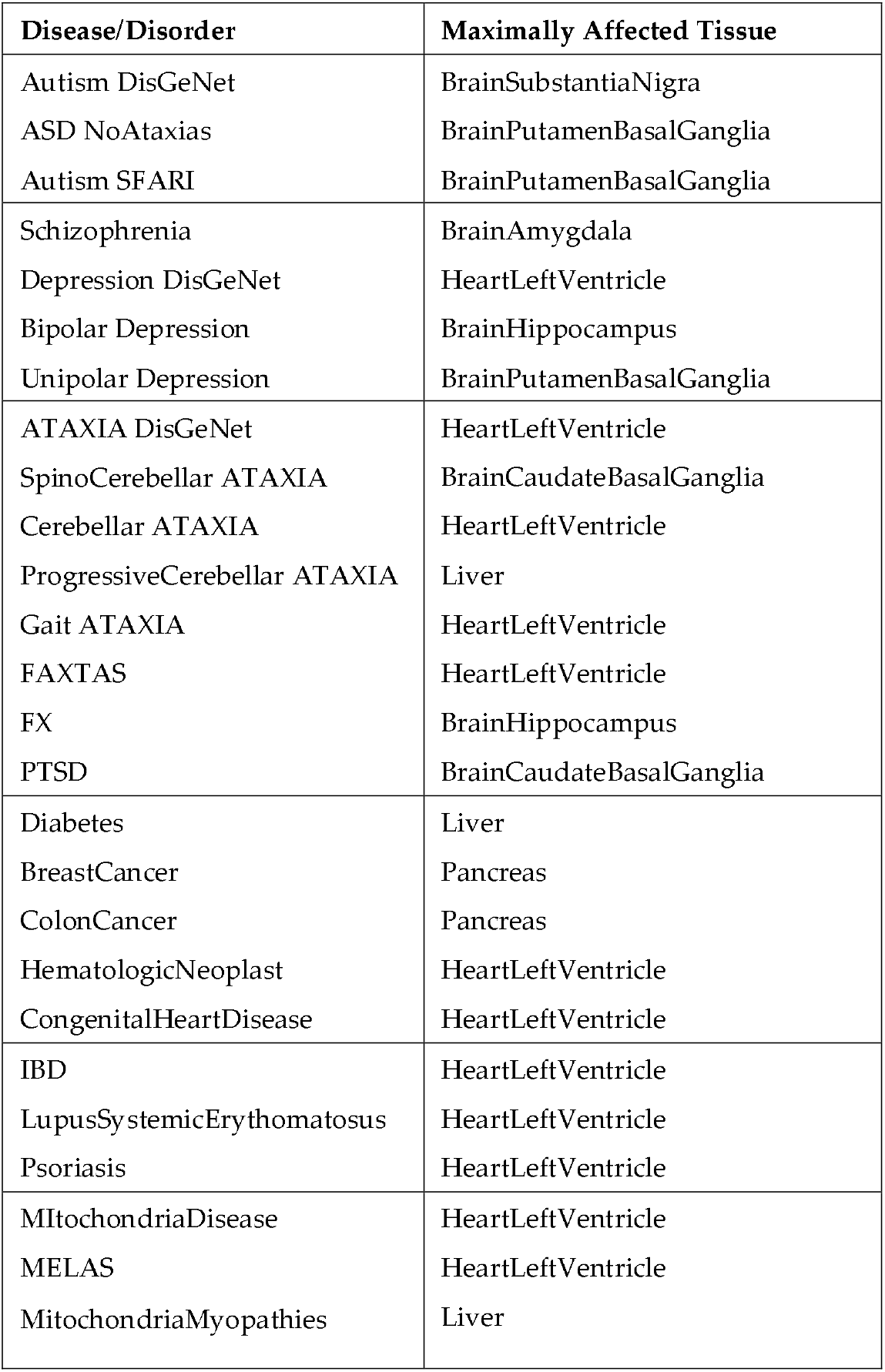
Top affected tissue in each disease/disorder according to the maximum Δλ value (See Figure 13 in the main text.)

## Footnotes

1 Taken from the site: “Transcripts Per Million (TPM) is a normalization method for RNA-seq, should be read as *for every 1,000,000 RNA molecules in the RNA-seq sample, x came from this gene/transcript*. For each transcript in the gene model, the number (raw count) of reads mapped is divided by the transcript’s length, giving a normalized transcript-level expression. The distribution of ambiguous reads (between transcripts of the same gene, or between different genes) is handled by OmicSoft’s RSEM implementation. The sum of ALL normalized transcript expression values is divided by 1,000,000, to create a scaling factor. Each transcript’s normalized expression is divided by the scaling factor, which results in the TPM value. Gene-level TPM’s are calculated by summing up the transcript-level TPM for each gene. In this scaling, the sum of all TPMs (transcript-level or gene-level) should always equal 1,000,000. For cells that have approximately the same number of transcripts-per-cell, the TPM expression values can be compared between these cells to estimate relative abundance. For a given sample, TPM values will linearly scale with FPKM values for genes or transcripts, but FPKM will not add up to 1,000,000, so TPM can also be thought as FPKM, scaled to sum to 1,000,000.”

## Notes

https://zenodo.org/record/3951607#.XxSGkShKiUk

### Competing Interest Statement

The authors have declared no competing interest.

