## Supplementary Material - Reframing Psychiatry for PM for "Reframing Psychiatry for Precision Medicine"

**Elizabeth B Torres** [1,2,3*]

[1] Rutgers University Department of Psychology;
[2] Rutgers University Center for Cognitive Science (RUCCS)
[3] Rutgers University Computer Science, Center for Biomedicine Imaging and Modelling (CBIM)

### Supplementary Material

Sample Psychological criteria that sidelines sensory motor issues in autism: The ADOS-2 manual [1, 2], under the "Guidelines for Selecting a Module" section states (emphasis added):

> "Note that the ADOS-2 was developed for and standardized using populations of children and adults *without significant sensory and motor impairments*. Standardized use of any ADOS-2 module presumes that the individual can walk independently and is free of visual or hearing impairments that could potentially interfere with use of the materials or participation in specific tasks."

Sample Psychiatric criteria from the DSM-5 [3] that does not include sensory-motor issues:

**A. Persistent deficits in social communication and social interaction across multiple contexts, as manifested by the following, currently or by history (examples are illustrative, not exhaustive, see text):**

1. Deficits in social-emotional reciprocity, ranging, for example, from abnormal social approach and failure of normal back-and-forth conversation; to reduced sharing of interests, emotions, or affect; to failure to initiate or respond to social interactions.
2. Deficits in nonverbal communicative behaviors used for social interaction, ranging, for example, from poorly integrated verbal and nonverbal communication; to abnormalities in eye contact and body language or deficits in understanding and use of gestures; to a total lack of facial expressions and nonverbal communication.
3. Deficits in developing, maintaining, and understanding relationships, ranging, for example, from difficulties adjusting behavior to suit various social contexts; to difficulties in sharing imaginative play or in making friends; to absence of interest in peers.

*Specify* current severity: Severity is based on social communication impairments and restricted repetitive patterns of behavior. (See table below.)

**B. Restricted, repetitive patterns of behavior, interests, or activities, as manifested by at least two of the following, currently or by history (examples are illustrative, not exhaustive; see text):**

1. Stereotyped or repetitive motor movements, use of objects, or speech (e.g., simple motor stereotypies, lining up toys or flipping objects, echolalia, idiosyncratic phrases).

2. Insistence on sameness, inflexible adherence to routines, or ritualized patterns or verbal nonverbal behavior (e.g., extreme distress at small changes, difficulties with transitions, rigid thinking patterns, greeting rituals, need to take same route or eat food every day).
3. Highly restricted, fixated interests that are abnormal in intensity or focus (e.g, strong attachment to or preoccupation with unusual objects, excessively circumscribed or perseverative interest).
4. Hyper- or hyporeactivity to sensory input or unusual interests in sensory aspects of the environment (e.g., apparent indifference to pain/temperature, adverse response to specific sounds or textures, excessive smelling or touching of objects, visual fascination with lights or movement).

**Supplementary Material Table 1**. Comparison of top-ranked tissues most affected by genes' removal according to the $\Delta\lambda$ values measuring the departure of the MLE $\lambda$ to model the genes' expression for each tissue as an Exponential memoryless random process. First column is from removal of the SFARI Autism genes. Second column is from removal of the Ataxia and X-Chromosome genes. Third column tissues result from the Fragile-X and Parkinson's disease genes and Fourth column tissues result from the removal of the genes associated with Mitochondrial disease. Sources for the genes are from the literature reported in Table 2 of the main text – Methods section. Acronyms are BG for Basal Ganglia; Anterior Cing Cortex for Anterior Cingulate Cortex; Nucleus Accum for Nucleus Accumbens; CellCultureFB for Cell Culture Fibroblast

| SFARI Autism | ATAXIAS & X | FX-PD | MITOCHONDRIA |
|---|---|---|---|
| Putamen BG | Whole Blood | Amygdala | Heart Left Ventricle |
| Substantia Nigra | Putamen BG | Substantia Nigra | Pancreas |
| Amygdala | Muscle Skeletal | Hippocampus | Muscle Skeletal |
| Hippocampus | Hippocampus | Kidney Cortex | Whole Blood |
| Caudate BG | Heart Left Ventricle | Brain Cortex | Hippocampus |
| Anterior Cing Cortex | Nucleus Accum BG | Nucleus Accum BG | Heart Atrial Appendage |
| Nucleus Accum BG | Caudate BG | Heart Left Ventricle | Substantia Nigra |
| Hypothalamus | Substantia Nigra | Hypothalamus | Amygdala |
| Brain Cortex | Amygdala | Putamen BG | Anterior Cing Cortex |
| Frontal Cortex | Anterior Cing Cortex | Heart Atrial Appendage | Brain Cortex |
| Heart Left Ventricle | Kidney Medulla | Anterior Cing Cortex | Esophagus Mucosa |
| Spinal Cord | Hypothalamus | Frontal Cortex | Liver |
| Pancreas | Heart Atrial Appendage | CellCultureFB | Kidney Medulla |

Acronyms are BG for Basal Ganglia; Cing for Cingulate; Accum for Accumbens

### Supplementary Figure 1

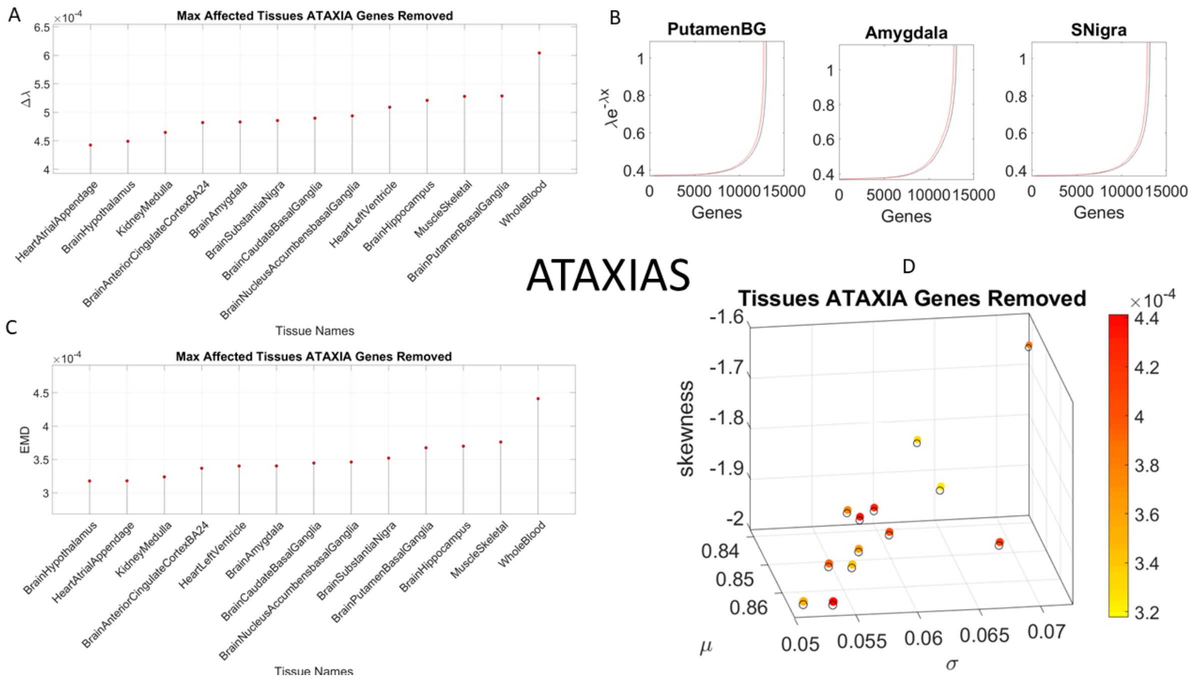


**Supplementary Figure 1. Outcome of genes' expression on most affected tissues upon removing the Ataxia genes from the normative data**. (A) Highest median-ranked set of most affected tissues according to $\Delta\lambda$ quantifying the difference in exponential distributions describing the stochastic signatures of each tissue based on the genes' count (TCM). (B) Shift in the exponential description of the counts across genes for selected converging tissues common to removing the SFARI Autism genes and the Ataxia genes (other tissues shown in Supplementary Table 1). (C) Consistent results are obtained using the continuous Gamma family of probability distributions empirically estimated using MLE for each tissue and EMD as a similarity metric to quantify departure from normative signatures upon Ataxia genes removal (inclusive of dominant, recessive and X-Chromosome genes involved in the Ataxias.) (D) Visualization of the stochastic shift on the Gamma parameter space are depicted using a color gradient to represent the intensity of the shift and circles to represent the position (mean, variance, skewness and kurtosis proportional to the marker size, larger markers are peakier distributions), with open circles as the normative data and colored filled ones as those resulting from removing the Ataxia genes.

### Supplementary Figure 2

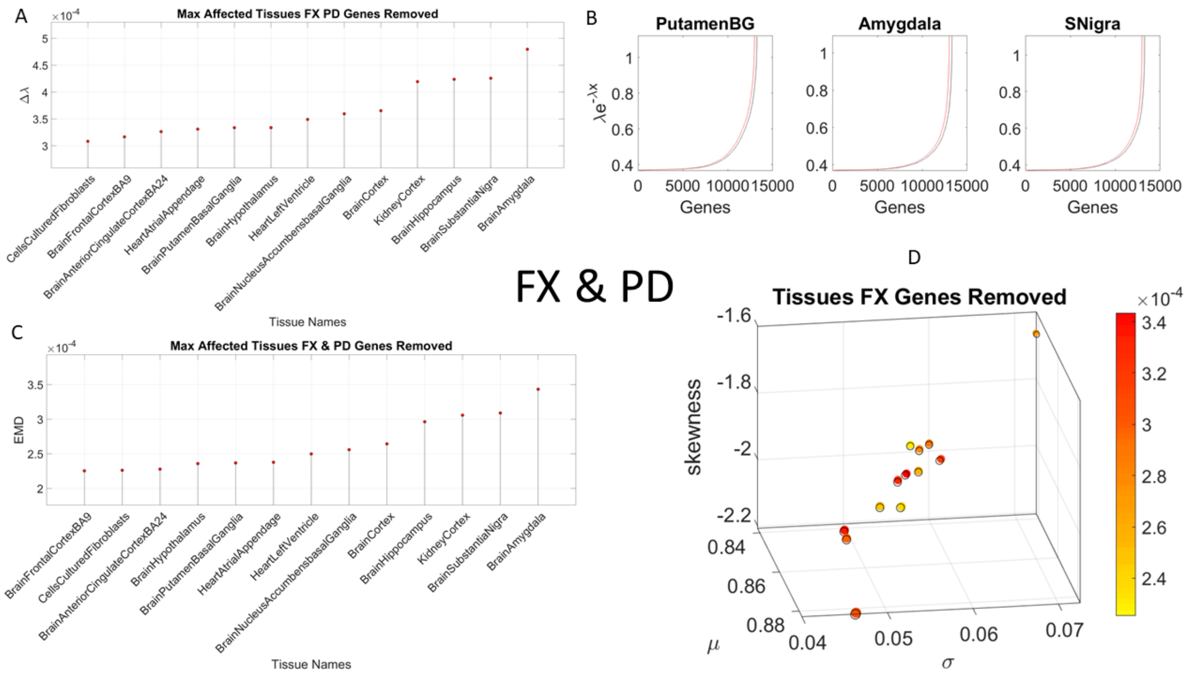


**Supplementary Figure 2. Outcome of genes' expression on most affected tissues upon removing the FX and PD genes from the normative data**. (A) Highest median-ranked set of most affected tissues according to $\Delta\lambda$ quantifying the difference in exponential distributions describing the stochastic signatures of each tissue based on the genes' count (TCM). (B) Shift in the exponential description of the counts across genes for selected converging tissues common to removing the SFARI Autism genes and the FX and PD genes (other tissues shown in Supplementary Table 1). (C) Consistent results are obtained using the continuous Gamma family of probability distributions empirically estimated using MLE for each tissue and EMD as a similarity metric to quantify departure from normative signatures upon FX and PD genes removal. (D) Visualization of the stochastic shift on the Gamma parameter space are depicted using a color gradient to represent the intensity of the shift and circles to represent the position (mean, variance, skewness and kurtosis proportional to the marker size, larger are peakier distributions), with open circles as the normative data and colored filled ones as those resulting from removing the FX and PD genes.

#### Supplementary Figure 3

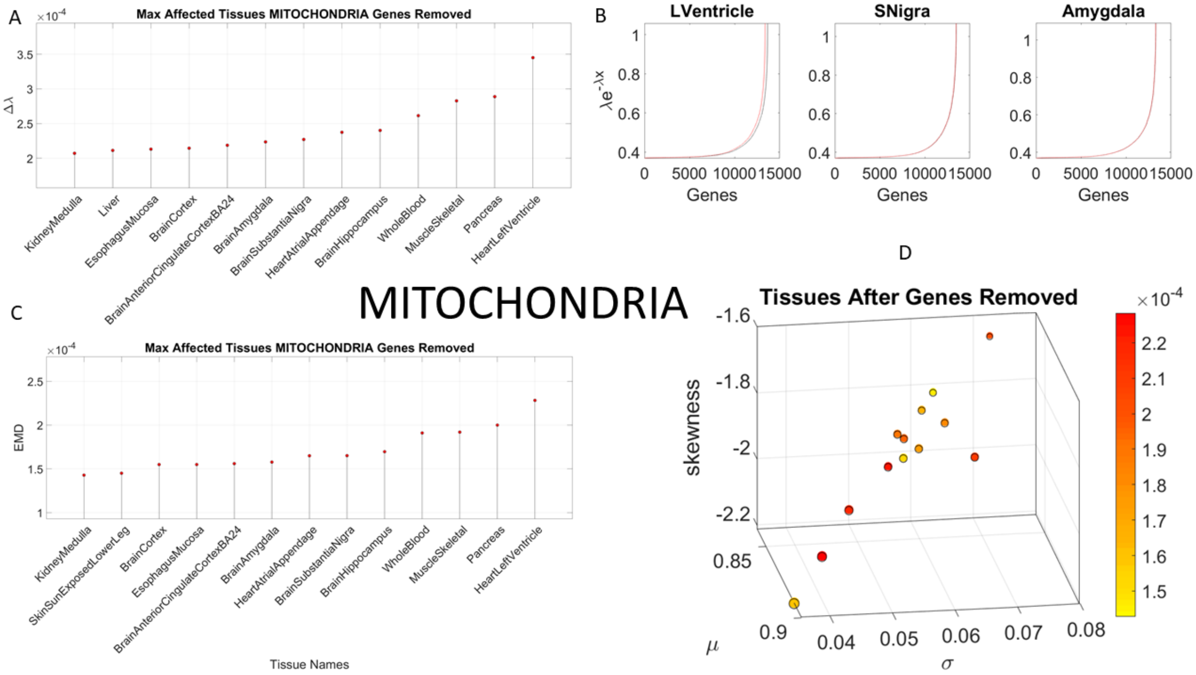


**Supplementary Figure 3. Outcome of genes' expression on most affected tissues upon removing the Mitochondrial disease genes from the normative data**. (A) Highest median-ranked set of most affected tissues according to $\Delta\lambda$ quantifying the difference in exponential distributions describing the stochastic signatures of each tissue based on the genes' count (TCM). (B) Shift in the exponential description of the counts across genes for selected converging tissues common to removing these and the SFARI Autism genes (other tissues shown in Supplementary Table 1). (C) Consistent results are obtained using the continuous Gamma family of probability distributions empirically estimated using MLE for each tissue and EMD as a similarity metric to quantify departure from normative signatures upon FX and PD genes removal. (D) Visualization of the stochastic shift on the Gamma parameter space are depicted using a color gradient to represent the intensity of the shift and circles to represent the position (mean, variance, skewness and kurtosis proportional to the marker size, larger are peakier distributions), with open circles as the normative data and colored filled ones as those resulting from removing the FX and PD genes.

### Supplementary Figure 4

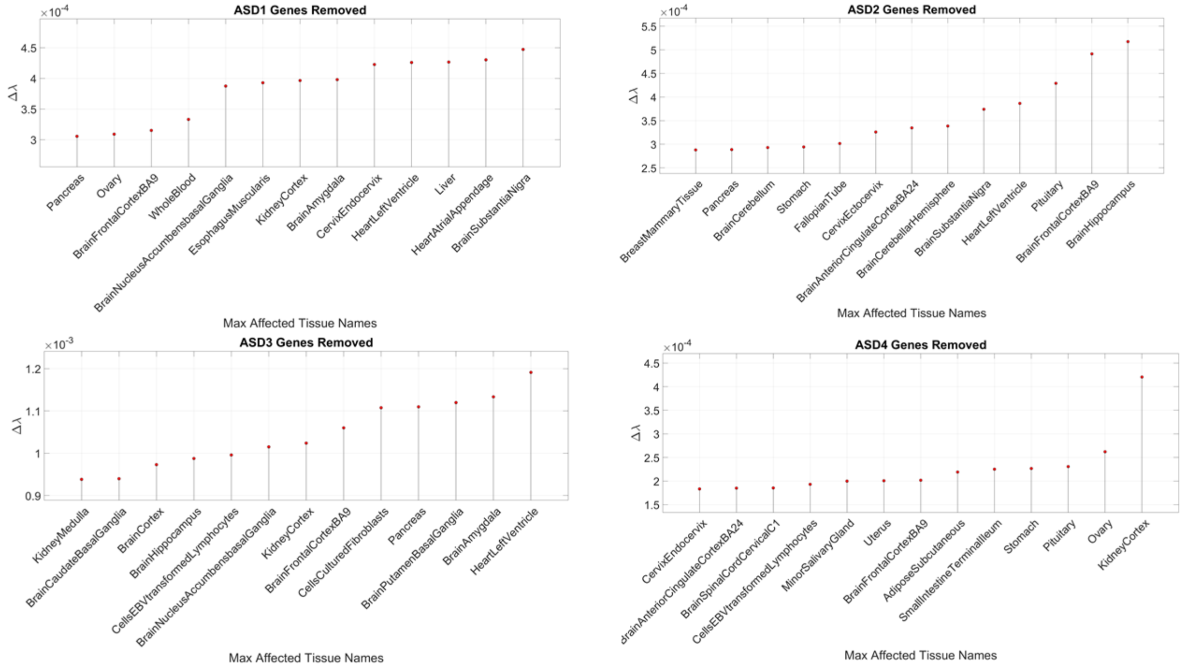


**Supplementary Figure 4. Outcome of genes' expression on most affected tissues upon removing genes based on SFARI Autism scores**. Maximally affected tissues in 1-3 scored SFARI Autism genes include the Substantia Nigra, Basal Ganglia and cerebellar tissues involved in motor control, adaptation/learning, regulation and coordination. They also include brain tissues involved in emotions (amygdala) and memory (hippocampus). All three top confidence genes include autonomic structures as well (heart atrial appendage and heart left ventricle). The syndromic genes (scored 4) include tissues vital to survival, with smooth muscles involved in involuntary functions such as secretion aiding swallowing peristalsis, digestion, and waste excretion. They also impact the brain frontal cortex (Brodmann area 9.)

Supplementary Table 2

| SFARI Autism (score) & Ataxia Dominant | Gene Name and Characteristics | Chromosome | GeneCategory rare/common variants | Aut/Tot Reports |
|---|---|---|---|---|
| CACNA1C (1)<br><br>Timothy Syndrome long QT syndrome | Calcium channel, voltage-dependent, L type, alpha 1C subunit involved in cardiac muscle functioning. By PCR of human heart, followed by sequence analysis, 3 variants of CACNA1C were identified [4], coined CACH2. (OMIM 114205) | 12p13.33 | Rare Single Gene Mutation, Syndromic, Genetic Association, Functional (31 / 9) | 12/45 |
| SCN2A (1)<br><br>Dravet Syndrome | Sodium channel, voltage-gated, type II, alpha subunit, voltage-gated ion channel essential for the generation and propagation of action potentials, **chiefly in nerve and muscle**. (OMIM 182390) | 2q24.3 | Rare Single Gene Mutation, Syndromic (213/0) | 33/62 |
| ATP1A3 (3S)<br>CAPOS Syndrome<br><br>Cerebellar ataxia, areflexia, pes cavus, optic atrophy, and sensorineural hearing loss<br><br>Dystonia-12 | ATPase Na+/K+ transporting subunit alpha 3. The protein encoded by this gene belongs to the family of P-type cation transport ATPases, and to the subfamily of Na+/K+ -ATPases. Na+/K+ -ATPase is an integral membrane protein responsible for establishing and maintaining the electrochemical gradients of Na and K ions across the plasma membrane. This gene encodes an alpha 3 catalytic subunit. Heterozygous variants in ATP1A3 are responsible for alternating hemiplegia of childhood 2 (AHC2; OMIM 614820) and (OMIM 601338) and see OMIM 182350 entry: The ATP1A3 isoform is exclusively expressed in neurons of various brain regions, including the basal ganglia, hippocampus, and cerebellum | 19q13.2 | Rare Single Gene Mutation, Syndromic, Functional (14/0) | 3/13 |
| CCDC88C (3)<br><br>congenital hydrocephalus<br>Spino Cerebellar Ataxia 40 | Coiled-coil domain containing 88C Encodes coiled-coil domain-containing protein that interacts with the dishevelled protein and is a negative regulator of the canonical Wnt signaling pathway, acting downstream of DVL to inhibit CTNNB1/Beta-catenin stabilization. (OMIM 616053) | 14q32.11-q32.12 | Rare Single Gene Mutation (8/0) | 6/6 |
| ITPR1 (3) | This gene encodes an intracellular receptor inositol 1,4,5-trisphosphate | | | 5/9 |

| SFARI Autism (score) & Ataxia Recessive | Gene Name and Characteristics | Chromosome | GeneCategory rare/common variants | Aut/Tot Reports |
|---|---|---|---|---|
| spinocerebellar ataxia SCA type 15 | receptor type 1. Upon stimulation by inositol 1,4,5-trisphosphate, this receptor mediates calcium release from the endoplasmic reticulum. | 3p26.1 | Rare Single Gene Mutation (11/0) | |
| KCNJ10 (2)<br><br>SeSAME syndrome | Potassium voltage-gated channel subfamily J member 10. This gene encodes a member of the inward rectifier-type potassium channel family, characterized by having a greater tendency to allow potassium to flow into, rather than out of, a cell. The encoded protein may form a heterodimer with another potassium channel protein and may be responsible for the potassium buffering action of glial cells in the brain. Expressed in renal epithelial cells, inner ear cells, and glial cells in the central nervous system (OMIM 602208) | 1q23.2 | Rare Single Gene Mutation, Syndromic, Genetic Association (11/2) | 4/11 |
| WWOX (2)<br><br>autosomal recessive spinocerebellar ataxia-12 (SCAR12)<br><br>Biallelic mutation can also cause early infantile epileptic encephalopathy-28 (a more severe, overlapping disorder) | WW domain containing oxidoreductase. This gene encodes a member of the short-chain dehydrogenases/reductases (SDR) protein family. This gene spans the FRA16D common chromosomal fragile site and appears to function as a tumor suppressor gene. Biallelic variants in the WWOX gene are responsible for early infantile epileptic encephalopathy-28 (EIEE28; OMIM 616211) | 16q23.1 | Rare Single Gene Mutation, Syndromic (25/9) | 3/9 |
| GRID2 (3)<br><br>SCA18 | Glutamate receptor, ionotropic, delta 2. Human glutamate receptor delta-2 (GRID2) is a relatively new member of the family of ionotropic glutamate receptors which are the predominant excitatory neurotransmitter receptors in the mammalian brain. GRID2 is a predicted 1,007 amino acid protein that shares 97% identity with the mouse homolog which is expressed selectively in cerebellar Purkinje cells. GRID2 is strongly suggested to have a role in neuronal apoptotic death. | 4q22.1-q22.2 | Rare Single Gene Mutation, Syndromic, Genetic Association (14/2) | 6/9 |
| LAMA1 (3) | Laminin, alpha 1. Binding to cells via a high affinity receptor, laminin is | | | 4/6 |

| | | | | |
|---|---|---|---|---|
| Tourette syndrome<br><br>Poretti-Boltshauser syndrome | thought to mediate the attachment, migration and organization of cells into tissues during embryonic development by interacting with other extracellular matrix components. | 18p11.31 | Rare Single Gene Mutation, Genetic Association (4/1) | |
| PEX7 (3)<br><br>Zellweger syndrome | Peroxisomal biogenesis factor 7. This gene encodes the cytosolic receptor for the set of peroxisomal matrix enzymes targeted to the organelle by the peroxisome targeting signal 2 (PTS2). Defects in this gene cause peroxisome biogenesis disorders (PBDs), which are characterized by multiple defects in peroxisome function. | 6q23.3 | Rare Single Gene Mutation, Genetic Association (2/2) | 2/3 |
| SYNE1(3S)<br><br>SCAR8 | Spectrin repeat containing, nuclear envelope 1. Multi-isomeric modular protein which forms a linking network between organelles and the actin cytoskeleton to maintain the subcellular spatial organization. | 6q25.2 | Rare Single Gene Mutation, Genetic Association (18/2) | 9/16 |
| CYP27A1 (4)<br>Cerebrotendinous xanthomatosis | Cytochrome P450 family 27 subfamily A member 1 | 2q35 | Rare Single Gene Mutation, Syndromic | 3 |
| SNX14 (4)<br>SCAR20 | Sorting nexin 14 | 6 q14.3 | Rare Single Gene Mutation, Syndromic | 6 |

| **SFARI Autism (score) & X** | **Gene Name and Characteristics** | **Chromosome** | **GeneCategory rare/common variants** | **Aut/Tot Reports** |
|---|---|---|---|---|
| CASK (1)<br>FG Syndrome 4<br>MICPCH Syndrome | This gene encodes a calcium/calmodulin-dependent serine protein kinase. The encoded protein is a MAGUK (membrane-associated guanylate kinase) protein family member. These proteins are scaffold proteins and the encoded protein is located at synapses in the brain. Mutations in this gene are associated with FG syndrome 4 (OMIM 300422) and mental retardation and microcephaly with pontine and cerebellar hypoplasia (OMIM 300749). | Xp11.4 | Rare Sinvle Gene Mutation, Syndromic 27/0 | 4/15 |
| FMR1 (1)<br>Fragile X syndrome<br>FX-Associated Tremor Ataxia Syndrome | The protein encoded by this gene binds RNA and is associated with polysomes. The encoded protein may be involved in mRNA trafficking from the nucleus to the cytoplasm. | Xq27.3 | Rare Single Gene Mutation, Syndromic, Genetic Association, Functional (31/0) | 7/12 |

| | | | | |
|---|---|---|---|---|
| SLC9A6 (1) | This gene encodes a sodium-hydrogen exchanger that is a member of the solute carrier family 9. The encoded protein may be involved in regulating endosomal pH and volume. | Xq26.3 | Rare Single Gene Mutation, Syndromic, Functional (24/0) | 2/15 |
| OPHN1 (2) | A Rho-GTPase-activating protein RhoGAP involved in cell migration and outgrowth of axons and dendrites. Loss of function of the protein is associated with X-linked mental retardation. | Xq12 | Rare Single Gene Mutation, Syndromic (17/0) | 4/15 |

| **SFARI Autism (score) & PD** | **Gene Name and Characteristics** | **Chromosome** | **GeneCategory rare/common variants** | **Aut/Tot Reports** |
|---|---|---|---|---|
| RAB39B (3) **Waisman Syndrome** **Early onset Parkinson's disease** | RAB39B, member RAS oncogene family. This gene encodes a member of the Rab family of proteins. Rab proteins are small GTPases that are involved in vesicular trafficking. | Xq28 | Rare Single Gene Mutation (12/0) | 6/12 |

Supplementary Table 2. Genes common to the SFARI Autism set and the Ataxias and PD-early onset literature. Information extracted from the SFARI file SFARI-Gene_genes_03-04-2020release_03-05-2020export.xlsx and from OMIM, the Online Mendelian Inheritance in Man database.

### Supplementary Figures of Genes in Supplementary Table 2 – **Autosomal Dominant Ataxia** genes overlapping with SFARI Autism genes

The following figures and text are compiled from various sources including SFARI Genes Module, GeneCard, OMMI, PubMed, and Wikipedia. Phenotypic descriptions include in mild cases, functional issues interfering with a broad range of activities of daily living. In severe cases, they have life threatening issues owing to autonomic dysfunction, malformations of tissues critical for vital functions (breathing, digestion and excretion mediated by smooth muscles.) The figures summarize the genes' expression in the tissues identified by the SFARI Autism genes that overlap with the Autosomal Dominant and Autosomal Recessive Ataxias and with the early-onset Parkinson's disease that we identified here as overlapping with the SFARI Autism gene. Violin plots reflecting log TMP are from GeneCard (in alphabetical order) and male and female genes' expression in brain tissues (Amygdala, Anterior Cingulate Cortex, Caudate Basal Ganglia, Cerebellar Hemisphere, Cerebellum, Cortex, Frontal Cortex, Hippocampus, Nucleus Accumbens Basal Ganglia, Putamen Basal Ganglia, Spinal Cord, Substantia Nigra), and in the Heart Left Ventricle and Pancreas identified by the top ranked tissues affected by removing from the GTEx human sample, the SFARI Autism genes.

#### Supplementary Figure 5 (CACNA1C Timothy Syndrome, Long QT Syndrome, Brugada Syndrome)

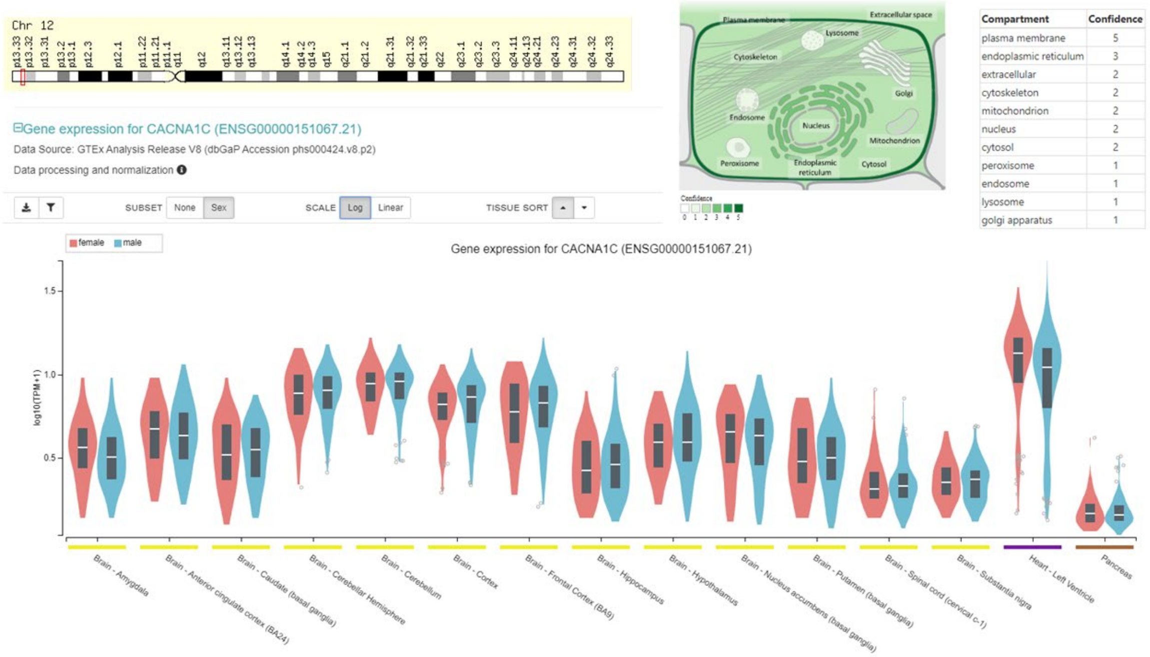


| Compartment | Confidence |
|---|---|
| plasma membrane | 5 |
| endoplasmic reticulum | 3 |
| extracellular | 2 |
| cytoskeleton | 2 |
| mitochondrion | 2 |
| nucleus | 2 |
| cytosol | 2 |
| peroxisome | 1 |
| endosome | 1 |
| lysosome | 1 |
| golgi apparatus | 1 |

**Supplementary Figure 5. Information on gene CACNA1C common to the SFARI Autism set and the Ataxia Autosomal Dominant genes reported in the literature**. This gene encodes an alpha-1 subunit of a voltage-dependent calcium channel. Calcium channels mediate the influx of calcium ions into the cell upon membrane polarization. Relevance to autism quoted from SFARI "This gene has been associated with syndromic autism, where a subpopulation of individuals with a given syndrome develop autism. In particular, mutation of the CACNA1C gene has been found to be associated with Timothy syndrome, patients which all also fall under the category of ASD. In addition, several studies have shown a genetic association between the CACNA1C gene and schizophrenia as well as bipolar disorder." Quoted from Wikipedia "The most striking sign of Timothy syndrome is the co-occurrence of both syndactyly (about 0.03% of births) and long QT syndrome (1% per year) in a single patient. Other common symptoms include cardiac arrhythmia (94%), heart malformations (59%), and autism or an autism spectrum disorder (80% who survive long enough for evaluation). Facial dysmorphologies such as flattened noses also occur in about half of patients. Children with this disorder have small teeth, which due to poor enamel coating, are prone to dental cavities and often require removal. **The average age of death due to complications of these symptoms is 2.5** years [5, 6]. Atypical Timothy syndrome has largely the same symptoms as the classical form. Differences in the atypical form are the lack of syndactyly, the presence of musculoskeletal problems (particularly hyperflexible joints), and atrial fibrillation. Patients with atypical Timothy syndrome also have more facial deformities, including protruding foreheads and tongues. Finally, one patient with atypical Timothy syndrome had a body development discrepancy wherein her upper body was normally developed (that of a 6-year-old) while her lower half resembled a 2- or 3-year-old [7, 8]. Children with Timothy syndrome tend to be born via caesarean section due to fetal distress. Diagnosis: Syndactyly and other deformities are typically observed and diagnosed at birth. Long QT syndrome sometimes presents itself as a complication due to surgery to correct syndactyly. Other times, children collapse spontaneously while playing. In all cases, Long QT syndrome is confirmed with ECG measurements. Sequencing of the *CACNA1C* gene further confirms the diagnosis. The disorder was thence named "Timothy syndrome" in honor of Katherine W. Timothy, who was among the first to identify a case and performed much of the phenotypic analysis that revealed other abnormalities [9]" https://en.wikipedia.org/wiki/Timothy_syndrome. Gene expression from GTex records shown for male and female genotypes in log-scale of TPM. Location in Chromosome and subcellular compartment information taken from: https://www.genecards.org/cgi-bin/carddisp.pl?gene=CACNA1C&search=CACNA1C. Long QT syndrome (LQTS) is a condition which affects repolarization of the heart after a heartbeat. It results in an increased risk of an irregular heartbeat which leads to fainting, drowning, or sudden death. These episodes can be triggered by exercise or stress (https://en.wikipedia.org/wiki/Long_QT_syndrome). See also Brugada syndrome (BrS) [10, 11] also cataloged as a genetic disorder in which the electrical activity within the heart is abnormal [12]. This disorder affects cardiac muscle function and increases the risk of abnormal heart rhythms and sudden cardiac death. Those affected may have episodes of passing out. The abnormal heart rhythms seen in those with BrS often occur at rest. They may be triggered by a fever. Physicians note that these children should avoid extraneous exercises (OMIM records).

### Supplementary Figure 6 (SCN2A Dravet Syndrome)

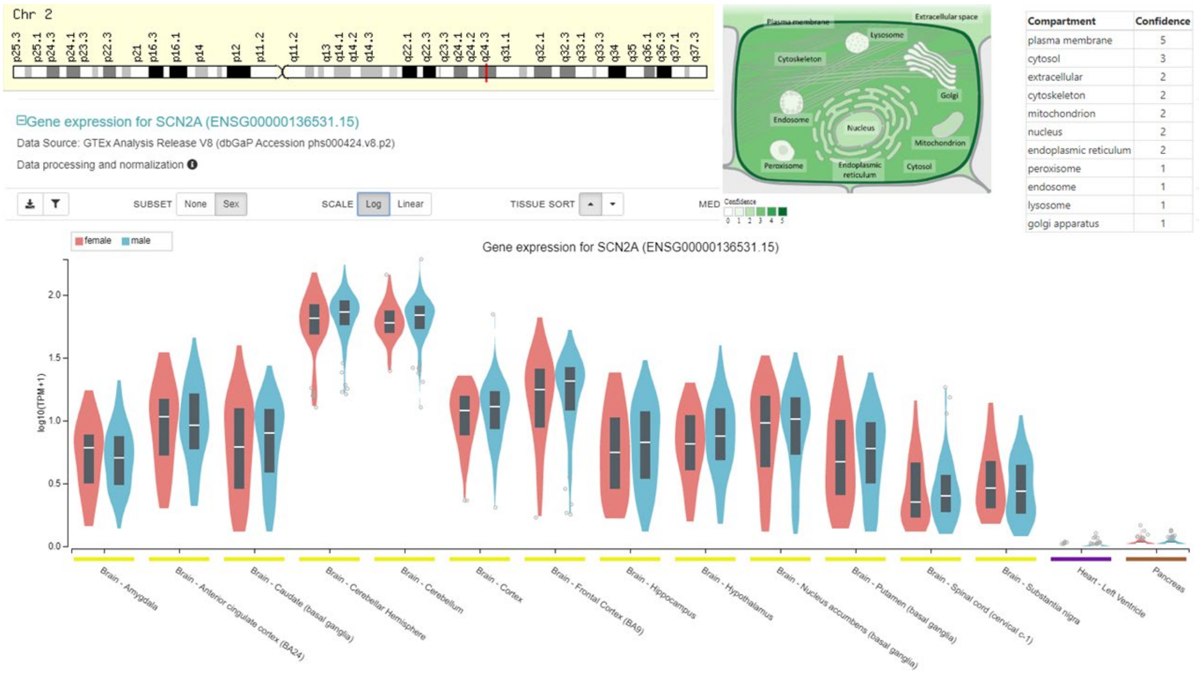


**Supplementary Figure 6. Information on gene SCN2A common to the SFARI Autism set and the Ataxia Autosomal Dominant genes reported in the literature**. Quoted from SFARI Genes Module: "Recurrent mutations in the SCN2A gene have been identified in multiple individuals with ASD as described below. Rare ASD-associated variants in the SCN2A gene were initially identified in a study by Weiss and colleagues in 2003 [13] based on exon screening in a region of linkage with autism. Sanders et al., 2012 subsequently reported 2 de novo loss-of-function (LoF) variants in SCN2A among 200 ASD families from the Simons Simplex Collection [14]. A third de novo LoF variant in the SCN2A gene was identified in a simplex ASD case in Tavassoli et al., 2014; this variant was not observed in dbSNP or other genomic databases [15]. A fourth de novo LoF variant in SCN2A was identified in a female ASD proband with intellectual disability in Jiang et al., 2013 [16]; this variant was not present in a female sibling with ASD but normal IQ. Analysis of rare coding variation in 3,871 ASD cases and 9,937 ancestry-matched or paternal controls from the Autism Sequencing Consortium (ASC) in De Rubeis et al., 2014 [17] identified SCN2A as a gene meeting high statistical significance with a FDR 0.01, meaning that this gene had a 99% chance of being a true autism gene. This gene was identified in Iossifov et al. 2015 [18] as a strong candidate to be an ASD risk gene based on a combination of de novo mutational evidence and the absence or very low frequency of mutations in controls. Functional analysis of ASD-associated de novo missense and likely gene disruptive SCN2A variants identified in probands from the Simons Simplex Collection and the Autism Sequencing Consortium using whole-cell voltage-clamp electrophysiology in Ben-Shalom et al., 2017 [19] found that these variants dampened or eliminated channel function, consistent with a LoF effect. Wolff et al., 2017 reported the phenotypes of 71 previously unpublished patients with SCN2A mutations; ASD was reported as a phenotype in 23 of these patients [20]." Besides ASD/Autism, this gene is associated with neurodevelopmental and intellectual disorders and attention deficit hyperactivity disorders (NDD, ID, ADHD respectively). Dravet syndrome, severe myoclonic epilepsy of infancy (SMEI), a catastrophic type of epilepsy with prolonged seizures that are often triggered by hot temperatures or fever is also associated with disruption in this gene. It is very difficult to treat with anticonvulsant medications and often begins before 1 year of age [21]. Diagnosis: Onset of seizures in the first year of life in an otherwise healthy infant; Initial seizures are typically prolonged and are generalized or unilateral. Presence of other seizure types (i.e. myoclonic seizures) and seizures associated with fever due to illness or vaccinations are also present. Seizures induced by prolonged exposure to warm temperatures and seizures in response to strong lighting or certain visual patterns have been also reported. Initially normal EEGs and later EEGs with slowing and severe generalized polyspikes are observed, along with normal initial development followed by slow development during the first few years of life. In the motor domain, some degree of **hypotonia** and u**nstable gait and balance issues** are reported with notable ankle pronation and flat feet and/or **development of a crouched gait with age.** Dravet syndrome is a severe form of epilepsy, responsible for roughly 10% of cases in children [22]. It is a rare genetic disorder that affects an estimated 1 in every 20,000–40,000 births [23, 24]. Charlotte Dravet first described severe myoclonic epilepsy of infancy in Centre Saint Paul, Marseille France in 1978 and the name was later changed to Dravet syndrome in 1989 [25, 26].

#### Supplementary Figure 7 (ATP1A3 CAPOS Syndrome)

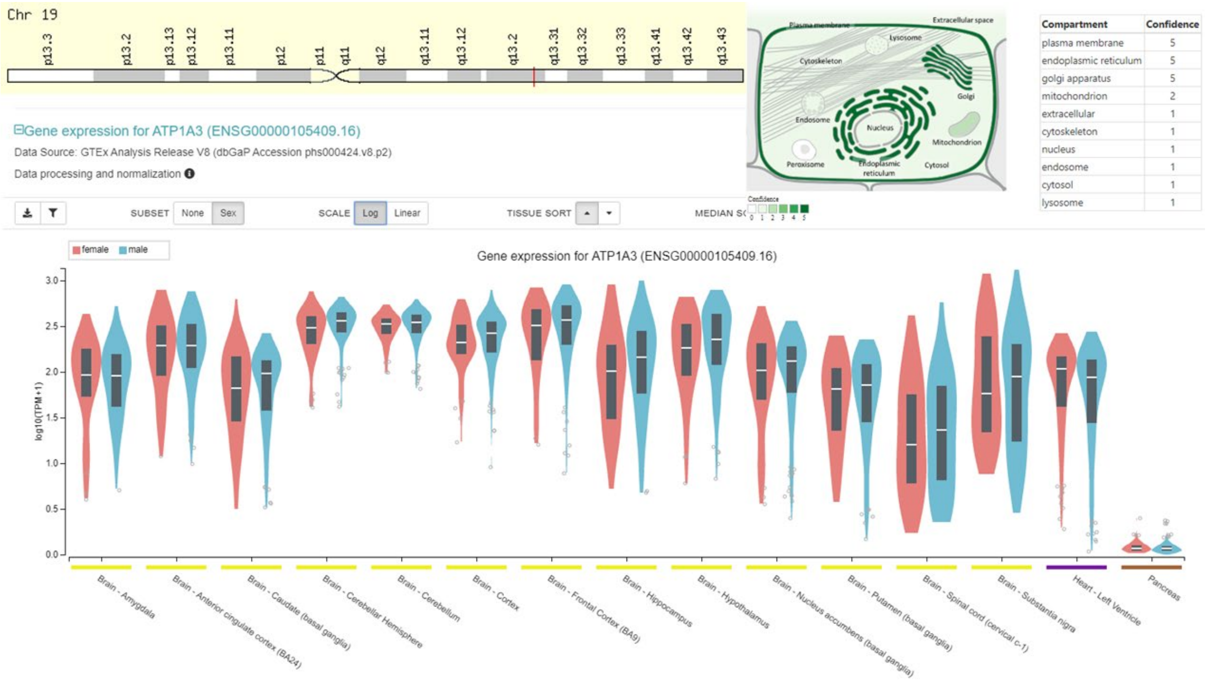


**Supplementary Figure 7. Information on gene ATP1A3 common to the SFARI Autism set and the Ataxia Autosomal Dominant genes reported in the literature**. Quoted from SFARI Genes Module: "Two siblings from a British family with CAPOS syndrome and a p.Glu818Lys missense variant in ATP1A3 were diagnosed with autism spectrum disorder; this variant was also observed in two additional families with CAPOS syndrome in the absence of ASD [27]. A probably damaging missense variant in ATP1A3 was identified in an ASD proband from the Autism Sequencing Consortium (De Rubeis et al., 2014). A second de novo missense variant that was predicted to be damaging was observed in an ASD proband from a Japanese trio in Takata et al., 2018. TADA analysis using a combined dataset of previously published cohorts from the Simons Simplex Collection and the Autism Sequencing Consortium, as well as the Japanese ASD cohort from Takata et al., 2018, identified ATP1A3 as a gene significantly enriched in damaging de novo mutations in ASD cases (pBH < 0.05)." From the orphaned site (orpha.net) (CAPOS syndrome) is a rare autosomal dominant neurological disorder characterized by early onset cerebellar ataxia, associated with areflexia, progressive optic atrophy, sensorineural deafness, a pes cavus deformity, and abnormal eye movements. It has a prevalence <1/1million with Autosomal dominant or mitochondrial inheritance. In 10 patients from 3 unrelated families with cerebellar ataxia, areflexia, pes cavus, optic atrophy, and sensorineural hearing loss [28-35] (CAPOS; OMIM 601338), including the original family reported by [27, 36] identified a heterozygous c.2452G-A transition in the ATP1A3 gene, resulting in a glu818-to-lys substitution at a highly conserved residue. The mutation, which was found by whole-exome sequencing of 2 of the families and confirmed by Sanger sequencing, was filtered against the dbSNP (builds 129 and 130) and 1000 Genomes Project databases and was not found in 1,834 controls. The mutation occurred de novo in the oldest affected generation of 1 family, but haplotype analysis could not rule out the possibility of a remote relationship between the other 2 families. All families were of Caucasian European descent. From OMIM 182350

Dystonia 12: The patient had a somewhat unusual phenotype, presenting at age 19 with rapidly progressive ataxia and dysarthria and tremor, resulting in loss of independent ambulation, and minimal dystonia. Exome sequencing showed that the patient also carried a de novo heterozygous missense E482K variant in the UBQLN4 gene, which may have played a role in the prominent cerebellar ataxia and cerebellar atrophy observed in this patient.

### Supplementary Figure 8 (CCDC88C) Congenital hydrocephalus and Spino Cerebellar Ataxia 40, SCA40

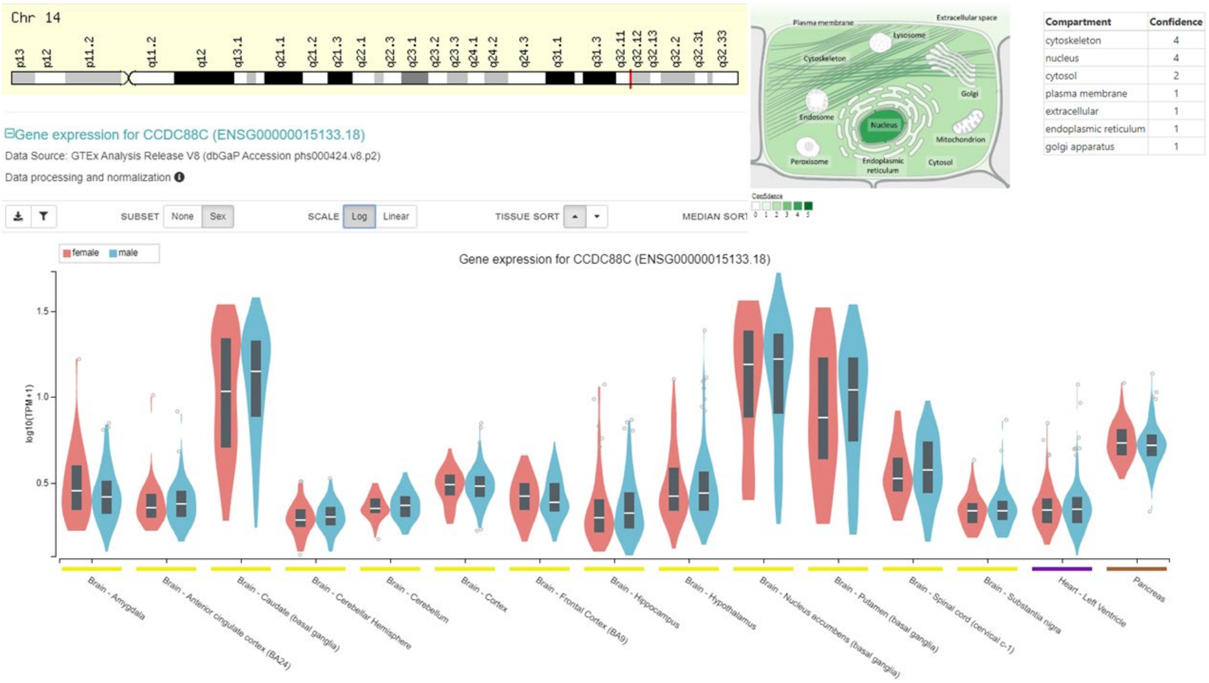


**Supplementary Figure 8. Information on gene CCDC88C common to the SFARI Autism set and the Ataxia Autosomal Dominant genes reported in the literature**. Quoted from SFARI Genes Module: "Two de novo missense variants in the CCDC88C gene were identified in simplex ASD probands, with no de novo events in this gene observed in 1,786 unaffected siblings from the Simons Simplex Collection (P=0.07) [37, 38]." From various sources: Presents congenital hydrocephalus, the child is born with an excessive accumulation of cerebrospinal fluid (CSF) in the brain. This typically causes increased pressure inside the skull. In babies, there is a rapid increase in head size. Other symptoms may include vomiting, sleepiness, seizures, and downward pointing of the eyes [39]. Older people may have headaches, double vision, poor balance, urinary incontinence, personality changes, or mental impairment. This condition may also be acquired as a consequence of CNS infections, meningitis, brain tumors, head trauma, toxoplasmosis, or intracranial hemorrhage (subarachnoid or intraparenchymal), and is usually painful (www.ucsfbenioffchildrens.org). Some phenotypic features are eyes that appear to gaze downward; irritability; seizures; separated sutures; sleepiness; vomiting. Symptoms that may occur in older children can include: brief, shrill, high-pitched cry; changes in personality, memory, or the ability to reason or think; changes in facial appearance and eye spacing (craniofacial disproportion); crossed eyes or uncontrolled eye; movements; difficulty feeding; excessive sleepiness; headaches; irritability, poor temper control; loss of bladder control (urinary incontinence); loss of coordination and trouble walking; muscle spasticity (spasm); slow growth (child 0–5 years); delayed milestones; failure to thrive; slow or restricted movement; vomiting [39] From OMIM 616053 "Tsoi et al. (2014) [40] reported a family from Hong Kong, China, in which 5 individuals had adult-onset spinocerebellar ataxia. Two affected individuals were described in detail. The 65-year-old proband presented with unsteady gait and dysarthria at age 43. Features included gait ataxia, wide-based gait, ocular dysmetria, intention tremor, scanning speech, dysdiadochokinesis, and hyperreflexia. She became a 'wheelchair user' 18 years after disease onset. Her younger brother presented with ataxic gait and dysarthria at age 42 years. He also had ocular dysmetria, impaired vertical gaze, scanning speech, and spastic paraparesis, and became a 'wheelchair user' 17 years after disease onset. Brain MRI of both patients showed pontocerebellar atrophy.”

#### Supplementary Figure 9 (ITPR1 SCA 15,16,29)

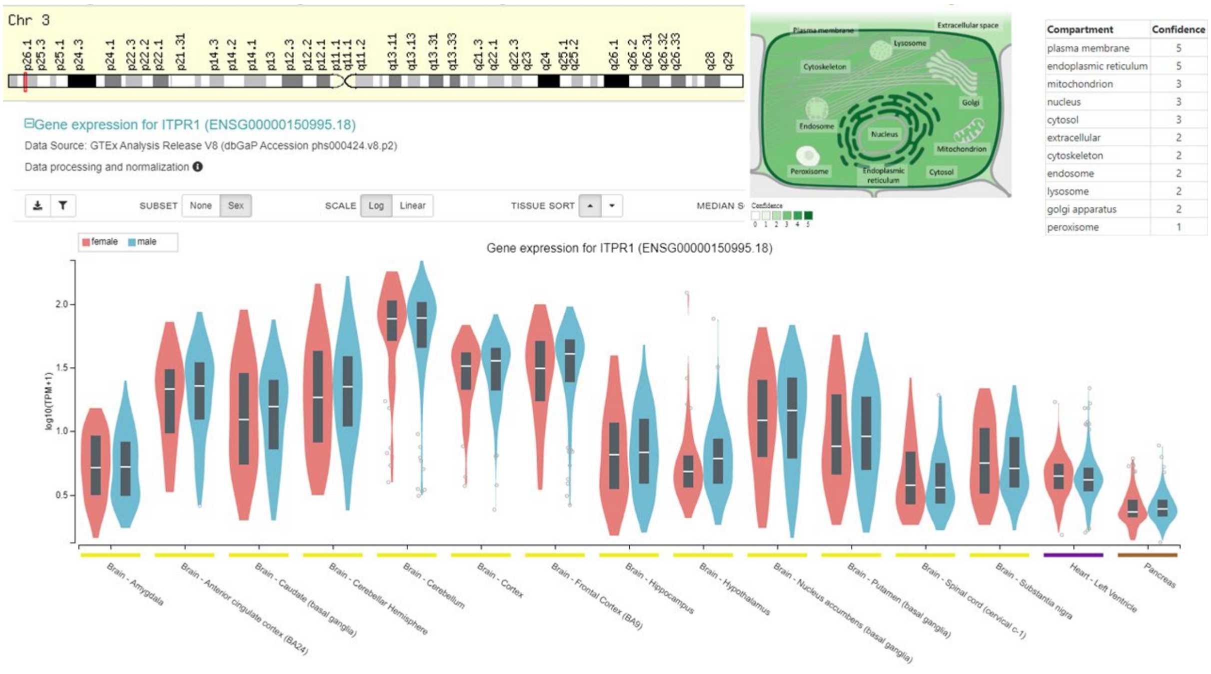


**Supplementary Figure 9. Information on gene ITPR1 common to the SFARI Autism set and the Ataxia Autosomal Dominant genes reported in the literature**. Quoted from SFARI Genes Module: “A likely damaging missense variant in the ITPR1 gene was identified in an ASD proband from the Autism Clinical and Genetic Resources in China (ACGC) cohort (Wang et al., 2016). Two additional possibly damaging de novo missense variants in ITPR1 had previously been identified in ASD probands from the Autism Sequencing Consortium [17] and the Simons Simplex Collection [37].” Compiled from various sources: is a progressive, degenerative (Dorland's Medical Dictionary) genetic disease with multiple types, each of which could be considered a neurological condition in its own right. An estimated 150,000 people in the United States have a diagnosis of spinocerebellar ataxia at any given time. SCA is hereditary, progressive, degenerative, and often fatal. There is no known effective treatment or cure. SCA can affect anyone of any age. In many cases people are not aware that they carry a relevant gene until they have children who begin to show signs of having the disorder (http://www.ninds.nih.gov/disorders/ataxia/ataxia.htm) Dominant hereditary spinocerebellar ataxia is included in a group of hereditary ataxia diseases and 40 different SCA forms have been described. It presents slowly progressive incoordination of gait and is often associated with poor coordination of hands, speech, and eye movements. A review of different clinical features among SCA subtypes was recently published describing the frequency of non-cerebellar features, like parkinsonism, chorea, pyramidalism, cognitive impairment, peripheral neuropathy, seizures, among others [41, 42]. As with other forms of ataxia, SCA frequently results in atrophy of the cerebellum, loss of fine coordination of muscle movements leading to unsteady and clumsy motion, and other symptoms. The most common cause of *acquired* cerebellar ataxia are stroke, multiple sclerosis (MS) and alcoholism. (https://en.wikipedia.org/wiki/Spinocerebellar_ataxia)

### Supplementary Figures of Genes in Supplementary Table 2 – **Autosomal Recessive Ataxia** genes overlapping with SFARI Autism genes

#### Supplementary Figure 10 (KCNJ10 SeSAME Syndrome)

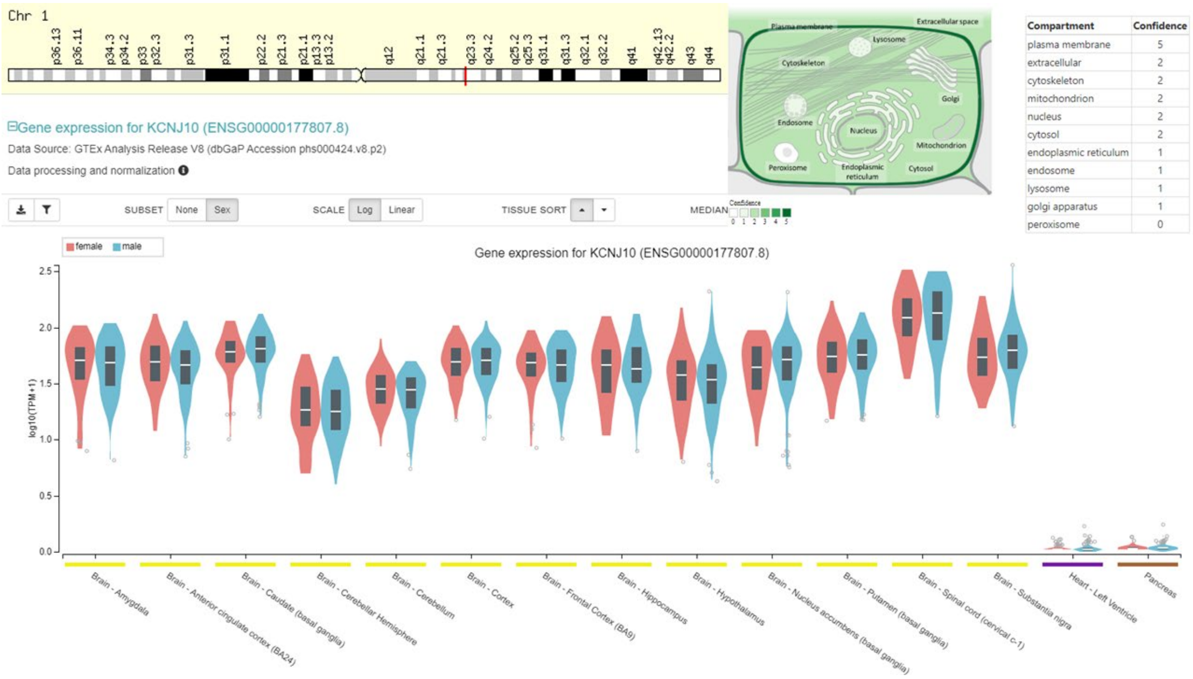


**Supplementary Figure 10. Information on gene KCNJ10 common to the SFARI Autism set and the Ataxia Autosomal Recessive genes reported in the literature**. Quoted from SFARI "Rare mutations in the KCNJ10 gene have been identified with autism (Sicca et al., 2011). In particular, that study found two non-synonymous SNPs (P18Q and V84M) in unrelated individuals with a seizure disorder who also have ASD and ID. Both of these mutations were shown to be functional in heterologous systems. In addition, genetic association has been found between KCNJ10 and seizure susceptibility in patients with epilepsy (Buono et al., 2004). Mutations in this gene have been associated with seizure susceptibility of common idiopathic generalized epilepsy syndromes." SeSAME syndrome is characterized by Seizures, Sensorineural deafness, Ataxia (lack of muscle coordination), intellectual (Mental) disability, and Electrolyte imbalance (low levels of potassium and magnesium in the blood, hypokalemia and hypomagnesemia, and metabolic alkalosis). It may also be known as EAST syndrome Epilepsy, Ataxia, Sensorineural deafness, and Tubulopathy (kidney problems [43] found KCNJ10 expression mainly distal to the macula densa in the early and late distal convoluted tubules, connecting tubules, and early cortical collecting ducts, but also in the thick ascending loop.) Seizures tend to start in early childhood. The seizures are typically of the generalized tonic-clonic seizure type (also known as grand mal seizures), but they usually respond well to medication. Non-progressive, cerebellar ataxia and hearing loss start later. Treatment includes antiepileptic medication, physical, educational and speech therapy, hearing aid and management of the kidney and electrolytes problems (OMIM 602208). Autosomal Recessive Deafness 4 With Enlarged Vestibular Aqueduct [44] identified mutations in the KCNJ10 gene associated with deafness with enlarged vestibular aqueduct (DFNB4; OMIM 600791) in probands from 2 families who also carried heterozygous mutations in the SLC26A4 gene (605646). The SLC26A4 mutations had been previously implicated in EVA-related hearing loss, and the KCNJ10 mutations reduced potassium conductance activity, which is critical for generating and maintaining the endocochlear potential. Yang and colleagues [44] also showed that haploinsufficiency of Slc26a4 in mouse results in reduced protein expression of KCNJ10 in the stria vascularis of the inner ear.

### Supplementary Figure 11 (WWOX SCA12)

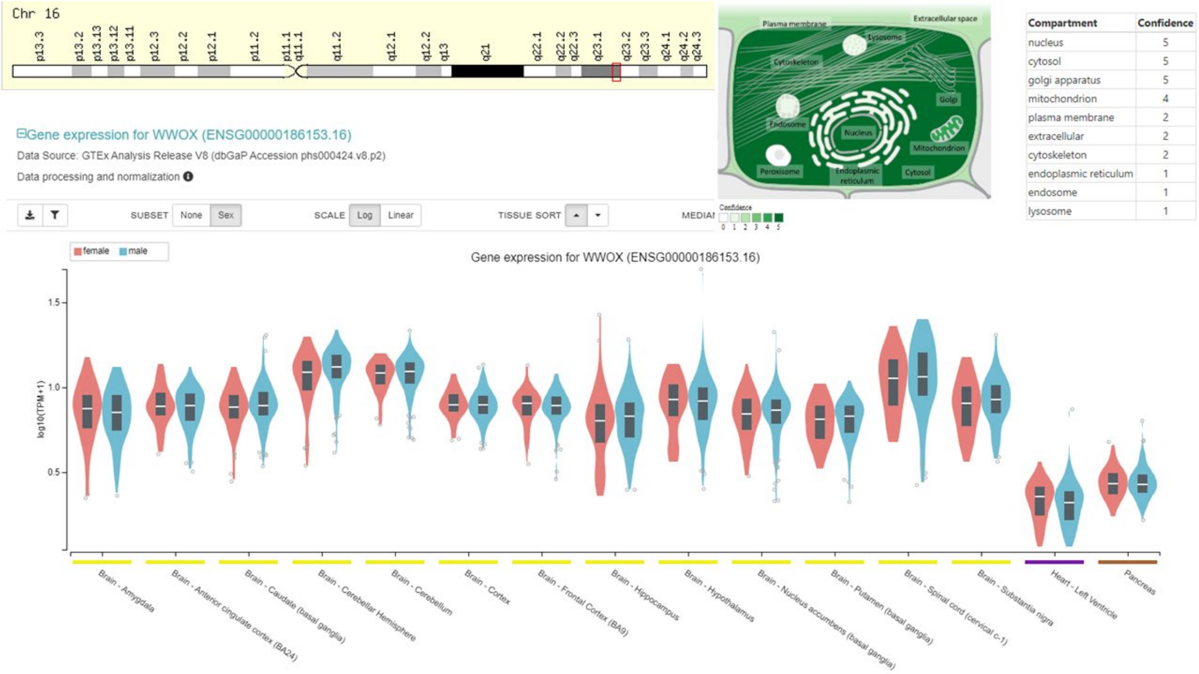


**Supplementary Figure 11. Information on gene WWOX common to the SFARI Autism set and the Ataxia Autosomal Recessive genes reported in the literature.** Autosomal recessive spinocerebellar ataxia-12 is a neurologic disorder characterized by onset of generalized seizures in infancy, delayed psychomotor development with mental retardation, and cerebellar ataxia. Some patients may also show spasticity (summary by [45]). The transmission pattern of spinocerebellar ataxia in the family reported by [46] was consistent with autosomal recessive inheritance.

### Supplementary Figure 12 (GRID2 Spinocerebellar ataxia, autosomal recessive 18)

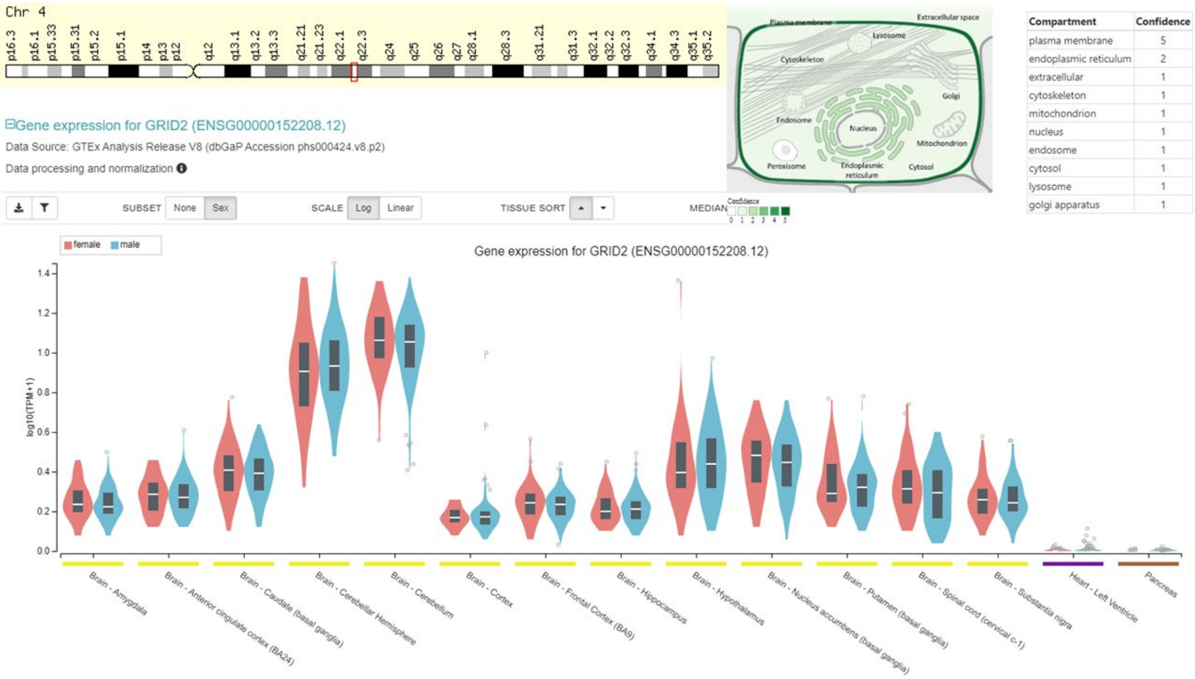


**Supplementary Figure 12. Information on gene GRID2 common to the SFARI Autism set and the Ataxia Autosomal Recessive genes reported in the literature**. Quoted from SFARI "Rare mutations in the GRID2 gene have been identified with ASD [47]. In particular, that study found two non-synonymous SNPs in GRID2 in 3 of 339 individuals with ASD." Spinocerebellar ataxia, autosomal recessive, 18 (SCAR18): A form of spinocerebellar ataxia, a clinically and genetically heterogeneous group of cerebellar disorders. Patients show progressive incoordination of gait and often poor coordination of hands, speech and eye movements, due to degeneration of the cerebellum with variable involvement of the brainstem and spinal cord. SCAR18 features include progressive cerebellar atrophy, delayed psychomotor development, severely impaired gait, ocular movement abnormalities, and intellectual disability. (OMIM 602368) The transmission pattern of SCAR18 in the families reported by Hills and colleagues [48] was consistent with autosomal recessive inheritance. They reported 3 children from a consanguineous Turkish family with early-onset cerebellar ataxia associated with eye movement abnormalities and intellectual disability. All patients had delayed psychomotor development with hypotonia, truncal and appendicular ataxia, and difficulty walking; 1 patient was wheelchair-bound at age 14 years. All patients also showed occasional or persistent tonic upgaze and nystagmus. Speech was severely limited. An unrelated child of Mexican descent, born of unrelated parents, had a similar phenotype. Brain imaging of 1 patient from the Turkish family and the Mexican patient showed progressive and severe cerebellar atrophy mainly affecting the flocculus. None of the patients had seizures or dysmorphic features. Utine and colleagues [49] reported 3 boys from a consanguineous Turkish kindred with cerebellar ataxia (at age 4 months boy presented nystagmus, hypotonia, and delayed psychomotor development, and later showed unsteady gait, incoordination of gross motor movements, and dysarthria.) Physical examination showed poor overall growth, rotatory nystagmus, truncal ataxia, oculomotor apraxia, pale optic disc, dysmetria, and dysdiadochokinesis. There was also evidence of pyramidal tract involvement, with hyperreflexia and extensor plantar responses. Serial brain imaging showed progressive cerebral and cerebellar atrophy. Two male cousins were similarly affected.

### Supplementary Figure 13 (LAMA1 Tourette's, Poretti-Boltshauser Syndrome)

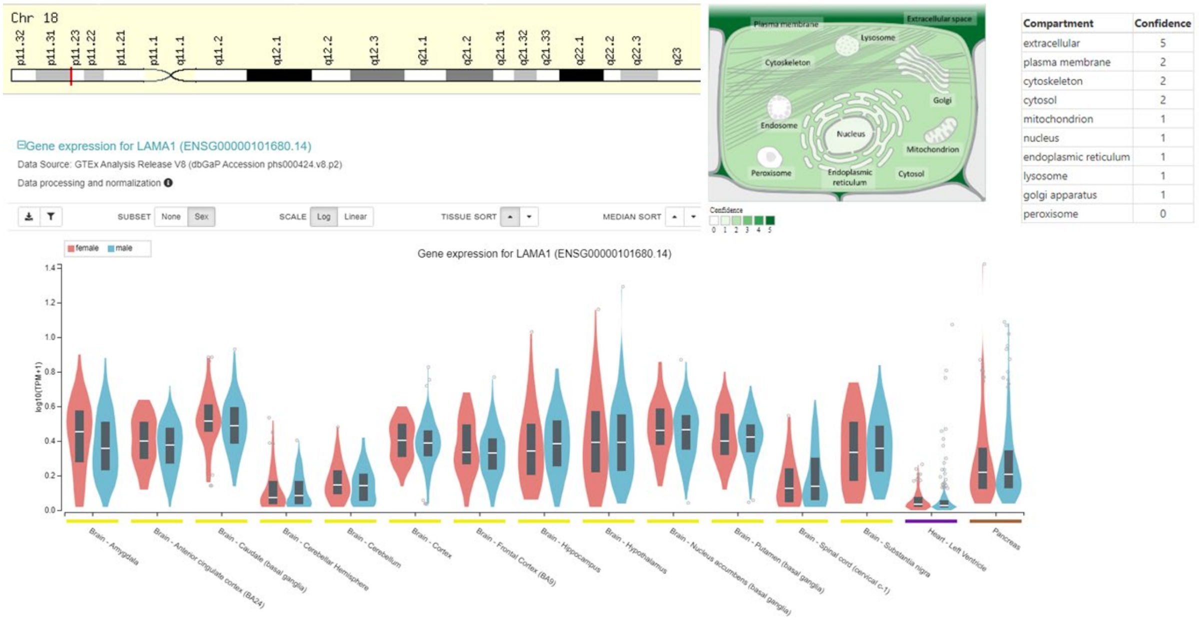


| Compartment | Confidence |
|---|---|
| extracellular | 5 |
| plasma membrane | 2 |
| cytoskeleton | 2 |
| cytosol | 2 |
| mitochondrion | 1 |
| nucleus | 1 |
| endoplasmic reticulum | 1 |
| lysosome | 1 |
| golgi apparatus | 1 |
| peroxisome | 0 |

**Supplementary Figure 13. Information on gene LAMA1 common to the SFARI Autism set and the Ataxia Autosomal Recessive genes reported in the literature**. Poretti-Boltshauser syndrome (OMIM 615960) is an autosomal recessive disorder characterized by cerebellar dysplasia, cerebellar vermis hypoplasia, cerebellar cysts in most patients, high myopia, variable retinal dystrophy, and eye movement abnormalities. Affected individuals have delayed motor development and often have speech delay. Cognitive function can range from normal to intellectually disabled [50]. Furthermore, cystic cerebellar dysplasia and biallelic *LAMA1* mutations leads to lamininopathy associated with tics, vocal tics, obsessive compulsive traits, and myopia due to cell adhesion and migration defects. Mildly wide-based gait; mild postural tremor and dysmetria of the upper extremities, oculomotor apraxia and severe anxiety [51].

#### Supplementary Figure 14 (PEX7 Zellweger syndrome)

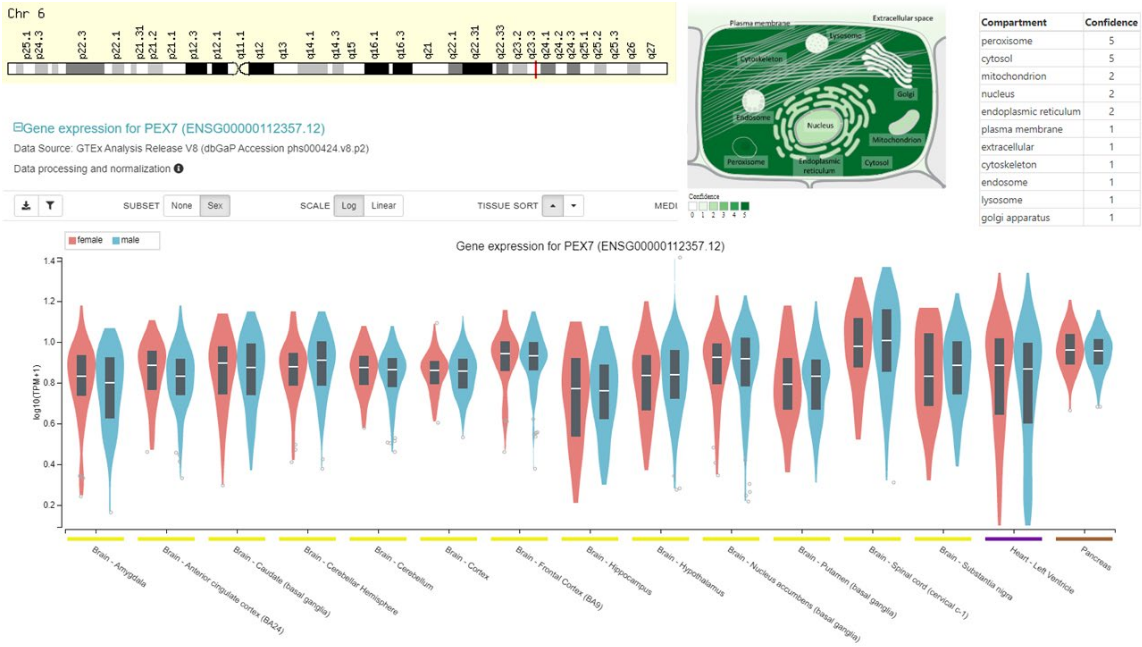


**Supplementary Figure 14. Information on gene PEX7 common to the SFARI Autism set and the Ataxia Autosomal Recessive genes reported in the literature**. Quoted from SFARI "Genetic association has been found between the PEX7 gene and ASD in a Korean population cohort (Ro et al., 2012). A homozygous loss-of-function missense variant in the PEX7 gene was identified in three affected children born to consanguineous parents from a multiplex ASD pedigree; this variant was heterozygous in both parents and in three unaffected children (Yu et al., 2013). The peroxisome biogenesis disorders (PBDs) are a group of genetically heterogeneous autosomal recessive, lethal diseases characterized by multiple defects in peroxisome function. Braverman et al. (1997) stated that at least 11 complementation groups (CGs) had been defined by somatic cell hybridization studies in patients with PBD phenotypes. Complementation groups 1 through 10 are not predictive of phenotype and contain patients with overlapping clinical features, including Zellweger syndrome (ZS; see 214100) as the most severe phenotype, neonatal adrenoleukodystrophy (NALD; see 601539), and infantile Refsum disease (IRD), an autosomal recessive disorder characterized by progressive adult retinitis pigmentosa, peripheral neuropathy, anosmia, and cerebellar ataxia, among other features. The onset of clinical symptoms in adolescence is due to gradual accumulation of phytanic acid. The peroxisomal enzyme phytanoyl-CoA hydroxylase (PHYH) catalyzes the first step of alpha-oxidation of phytanic acid. Biochemical analyses in Refsum patients reported by Van den Brink and colleagues [52] showed defects not only in phytanic acid alpha-oxidation, but also in plasmalogen synthesis and peroxisomal thiolase. These findings indicated that mutations in PEX7 may result in a broad clinical spectrum ranging from severe rhizomelic chondrodysplasia punctata to relatively mild Refsum disease, and that clinical diagnosis of conditions involving retinitis pigmentosa, ataxia, and polyneuropathy may require a full screen of peroxisomal functions.. Zellweger syndrome patients have developmental abnormalities as well as progressive dysfunction of the liver and central nervous system, leading to death before the end of the first year. Patients with NALD and IRD have similar but milder features. The entire collection of phenotypes was referred to by Braverman et al. (1997) as the 'Zellweger syndrome spectrum.' Metabolic abnormalities characteristic of these patients include deficiency of plasmalogens and accumulation of phytanic acid and very long chain fatty acid (VLCFA). At the cellular level, these patients exhibit deficiency of multiple peroxisomal enzymes. Zellweger syndrome is associated with impaired neuronal migration, neuronal positioning, and brain development. In addition, individuals with Zellweger syndrome can show a reduction in central nervous system (CNS) myelin (particularly cerebral), which is referred to as **hypomyelination**. Myelin is critical for normal CNS functions, and in this regard, serves to insulate nerve fibers in the brain. Patients can also show postdevelopmental sensorineuronal degeneration that leads to a progressive loss of hearing and vision [53, 54]. Zellweger syndrome can also affect the function of many other organ systems. Patients can show craniofacial abnormalities (such as a high forehead, hypoplastic supraorbital ridges, epicanthal folds, midface hypoplasia, and a large fontanel), hepatomegaly (enlarged liver), chondrodysplasia punctata (punctate calcification of the cartilage in specific regions of the body), eye abnormalities, and renal cysts. Newborns may present with profound hypotonia (low muscle tone), seizures, apnea, and an inability to eat [55].

#### Supplementary Figure 15 (SYNE1)

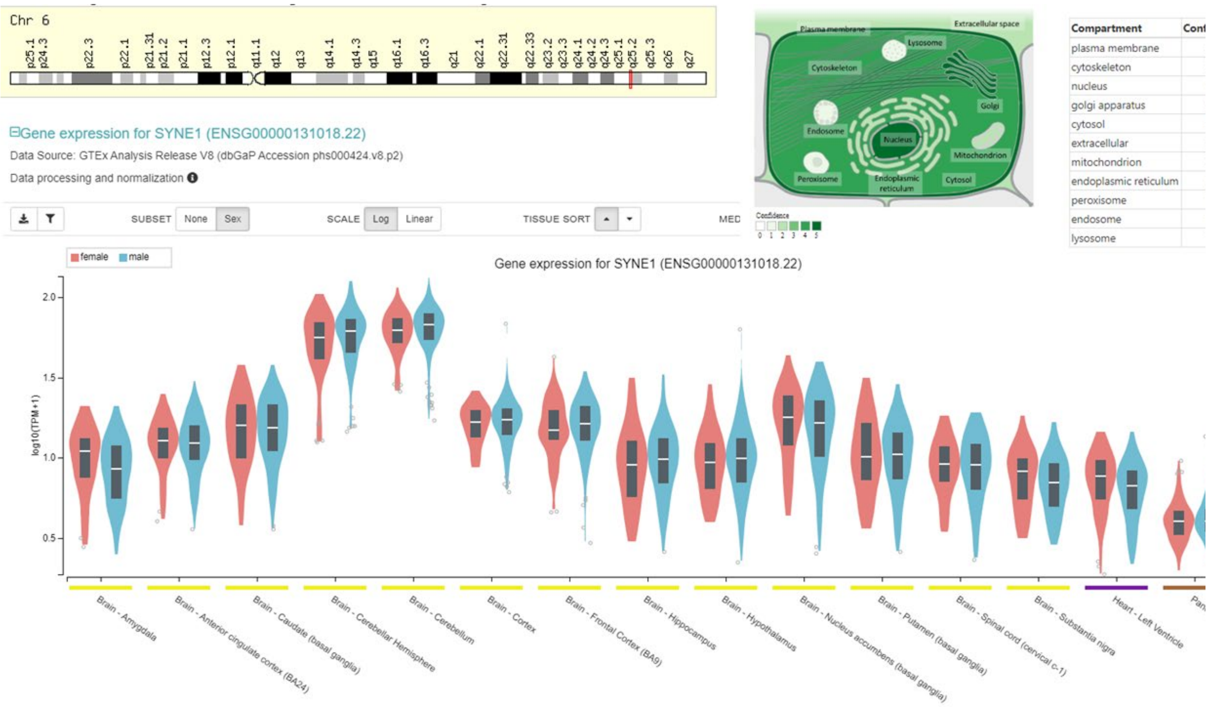


**Supplementary Figure 15. Information on gene SYNE1 common to the SFARI Autism set and the Ataxia Autosomal Recessive genes reported in the literature**. (OMIM 610743) Autosomal recessive spinocerebellar ataxia-8 (SCAR8) is a slowly progressive neurodegenerative disorder characterized by gait ataxia and other cerebellar signs, such as nystagmus and dysarthria. The age at onset is highly variable, and but most often is in the second or third decades. The disorder was initially identified in patients of French-Canadian descent, most of whom have a relatively 'pure' form of the disorder. However, subsequent studies have shown that SCAR8 occurs worldwide and most commonly manifests with additional features, including spasticity, secondary musculoskeletal abnormalities, and ocular movement anomalies, consistent with a 'complicated' phenotype. Brain imaging typically shows cerebellar atrophy, sometimes with pontine involvement. Rare patients may have an early-onset multisystemic disorder with impaired intellectual development and respiratory dysfunction [56].

Supplementary Figure 16 (CYP27A1 Cerebrotendinous Xanthomatosis (CTX))

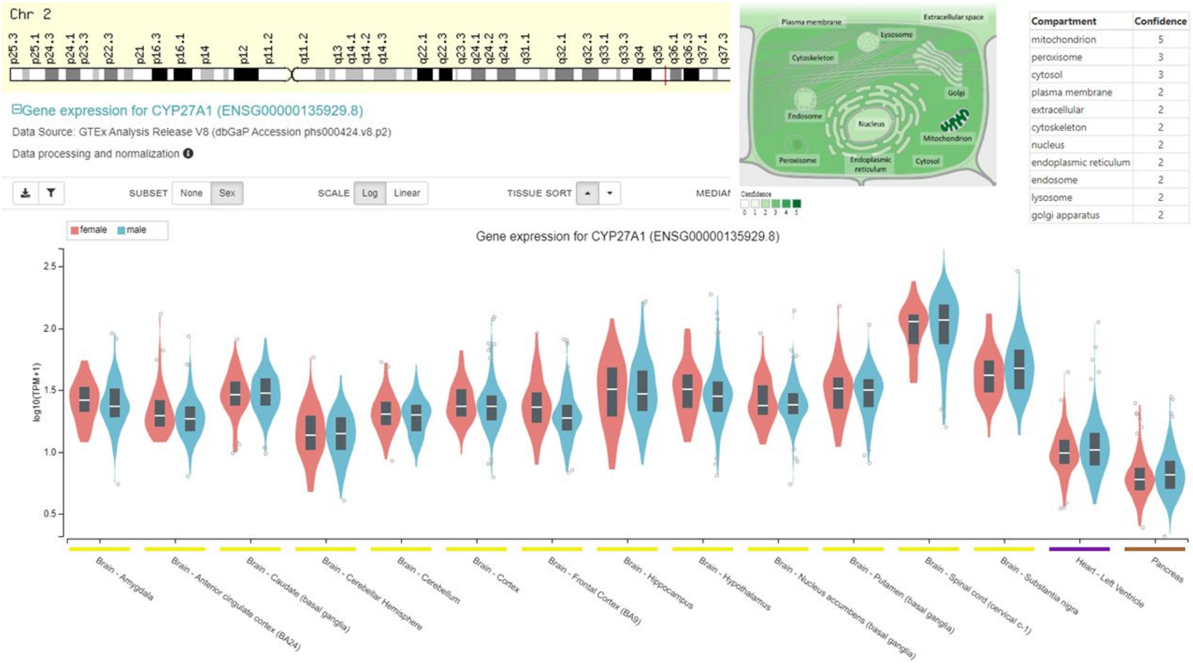


**Supplementary Figure 16. Information on gene CYP27A1 common to the SFARI Autism set and the Ataxia Autosomal Recessive genes reported in the literature**. (OMIM 213700) Cerebrotendinous xanthomatosis is a rare inherited lipid-storage disease characterized clinically by progressive neurologic dysfunction (cerebellar ataxia beginning after puberty, systemic spinal cord involvement and a pseudobulbar phase leading to death), premature atherosclerosis, and cataracts. Large deposits of cholesterol and cholestanol are found in virtually every tissue, particularly the Achilles tendons, brain, and lungs. Cholestanol, the 5-alpha-dihydro derivative of cholesterol, is enriched relative to cholesterol in all tissues. The diagnosis can be made by demonstrating cholestanol in abnormal amounts in the serum and tendon of persons suspected of being affected. Plasma cholesterol concentrations are low normal in CTX patients. Dotti and colleagues [57] examined the ophthalmologic findings of 13 CTX patients. In addition to cataracts, which were found in all cases, optic disc pallor was identified in 6 of the patients. Premature retinal senescence was also observed. Epidemiology Onset: Post-pubertal Ataxia; High incidence in Moroccan Jews; Neuropathy (60%); Clinical presentation shows sensory loss: Distal; Symmetric Weakness: Distal legs; Symmetric Tendon reflexes: Reduced Nerve conduction testing: Axonal loss Nerve Pathology; Axon loss: Reduced numbers of Large myelinated axons; Demyelination: Some patients; Thin myelin; Small onion bulbs; CNS Cerebellar signs (60%); Upper motor neuron disorders (90%); Pseudobulbar Dysfunction; Progressive Myoclonus Dementia; Systemic; Premature atherosclerosis; Xanthelasma: Tendons thickened; Cataracts: Premature, Bilateral presentation.

### Supplementary Figure 17 (SNX14 SCAR20)

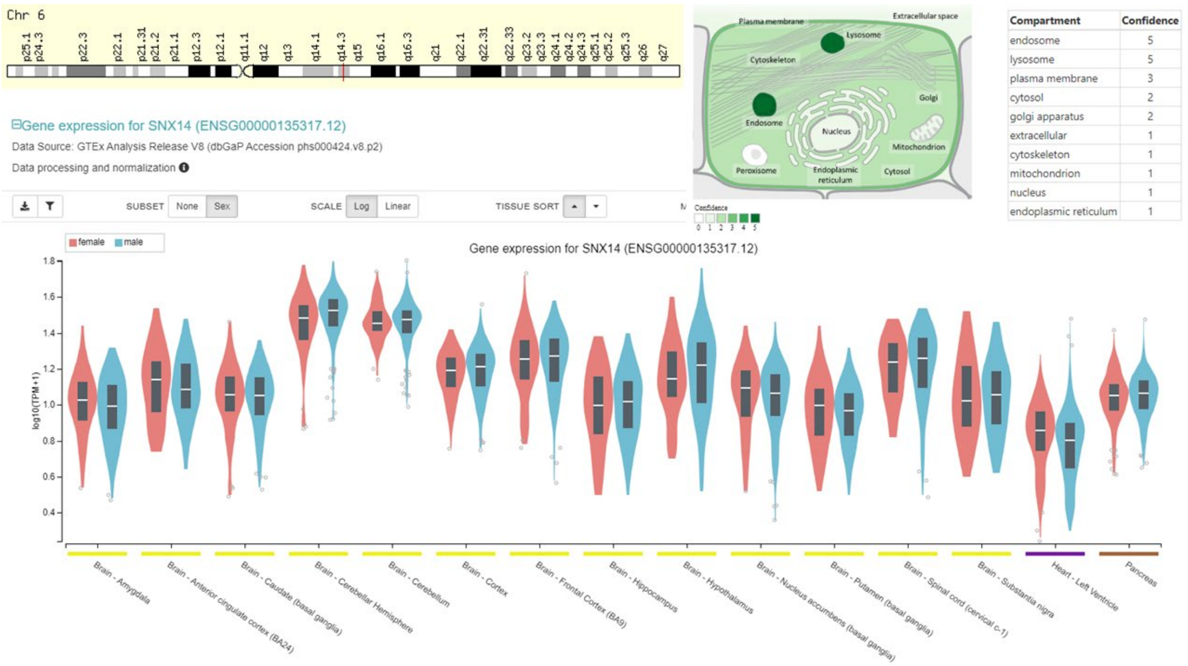


**Supplementary Figure 17. Information on gene SNX14 common to the SFARI Autism set and the Ataxia Autosomal Recessive genes reported in the literature**. Autosomal recessive spinocerebellar ataxia-20 is a neurodevelopmental disorder characterized by severely delayed psychomotor development with poor or absent speech, wide-based or absent gait, coarse facies, and cerebellar atrophy (summary by Thomas and colleagues [58] reported 5 children from 2 unrelated consanguineous families who had developmental delay, severe intellectual disability, and cerebellar ataxia. The patients also had progressive coarsening of facial features, relative macrocephaly, and progressive cerebellar atrophy on brain imaging. Some had sensorineural hearing loss; seizures were not present. Apraxia, a defect in the understanding of complex motor commands and in the execution of certain learned movements is commonly present, i.e., deficits in the cognitive components of learned movements. Balance issues (cerebellar ataxia) is also common from cerebellar degeneration. This causes a variety of elementary neurological deficits including asynergy (lack of coordination between muscles, limbs and joints), dysmetria (lack of ability to judge distances that can lead to under-overshoot in grasping movements), and dysdiadochokinesia (inability to perform rapid movements requiring antagonizing muscle groups to be switched on and off repeatedly). Absent speech or lack of development of speech and spoken language is also common. The transmission pattern of SCAR20 in the families reported by [58-60] was consistent with autosomal recessive inheritance.

### Supplementary Figure 18 (RAB39B Waisman syndrome or Early onset Parkinson's Disease)

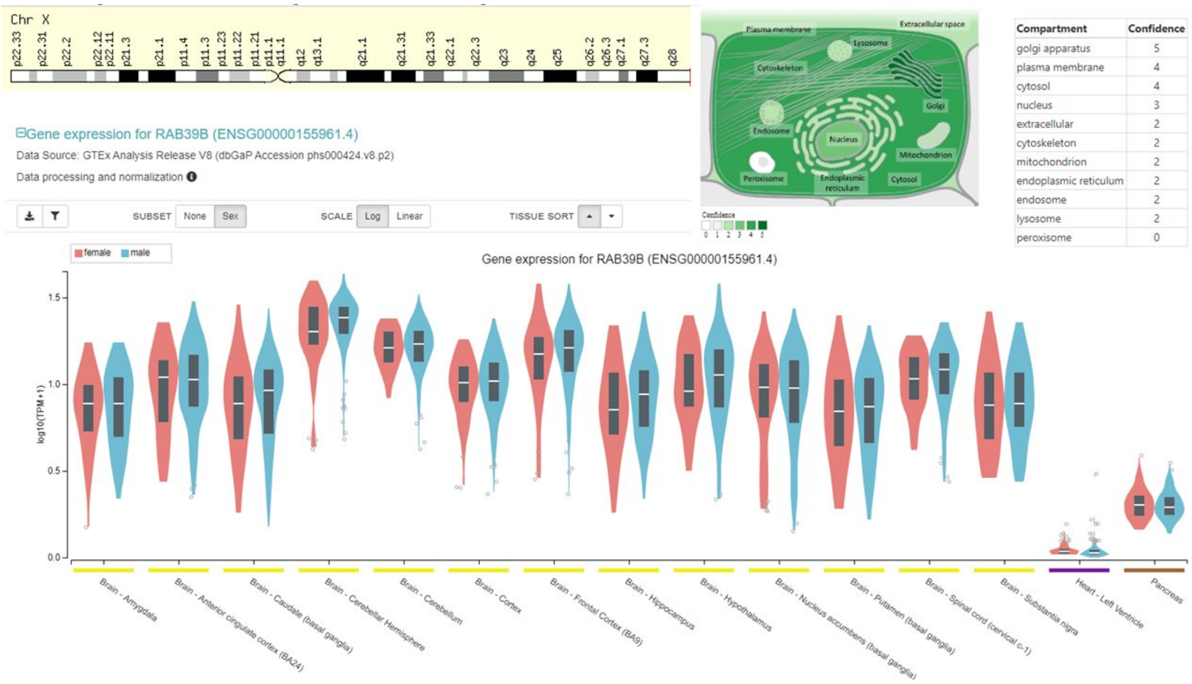


**Supplementary Figure 18. Information on gene RAB39B common to the SFARI Autism set and the Ataxia Autosomal Recessive genes reported in the literature.** Quoted from SFARI "Rare mutations in the RAB39B gene have been identified with autism (Giannandrea et al., 2010)." (OMIM 300774) The disorder is characterized by delayed psychomotor development, intellectual disability, and early-onset Parkinson disease. In vitro cellular studies showed that loss of RAB39B was associated with reduced steady-state levels of alpha-synuclein. Wilson and colleagues [61-63] concluded that dysregulation of SNCA homeostasis and defects in vesicular trafficking resulted in the manifestations of this neurologic disorder. Mutations in the RAB39B gene were not found in 187 individuals with early-onset Parkinson disease or in 48 males with neurodegeneration with brain iron accumulation. Lochte and colleagues [64] sequenced the RAB39B gene, including all introns and intron-exon boundaries, in 552 Parkinson disease patients including 330 males. Patients were mostly of German origin but also included 8 Filipinos. Mean age of onset was 50.9 +/- 15.3 years, and family history was positive in 22.2%. None of the pedigrees suggested X-linked inheritance. Lochte and colleagues also sequenced 91 Filipinos with X-linked dystonia-parkinsonism (XDP in OMIM 314250 in Xq13.1), 8 Filipinos with XDP phenotype who did not carry the XDP haplotype, and 186 German controls for variants in RAB39B. They identified 3 variants (1 synonymous, 2 intronic), for which evidence for pathogenicity was inconclusive. Noting that their findings combined with those of Yang and colleagues [65, 66]showed no convincing mutation in RAB39B in more than 1,000 Parkinson disease patients, Lochte and colleagues concluded that RAB38B mutations in classic Parkinson disease patients *without intellectual disability* and *without clear X-linked inheritance* are rare and do not need to be included in genetic testing.

### Supplementary Figures of Genes in Supplementary Table 2 – **X-Chromosome** genes with the SFARI Autism genes

#### Supplementary Figure 19 (CASK MICPCH syndrome, FG syndrome)

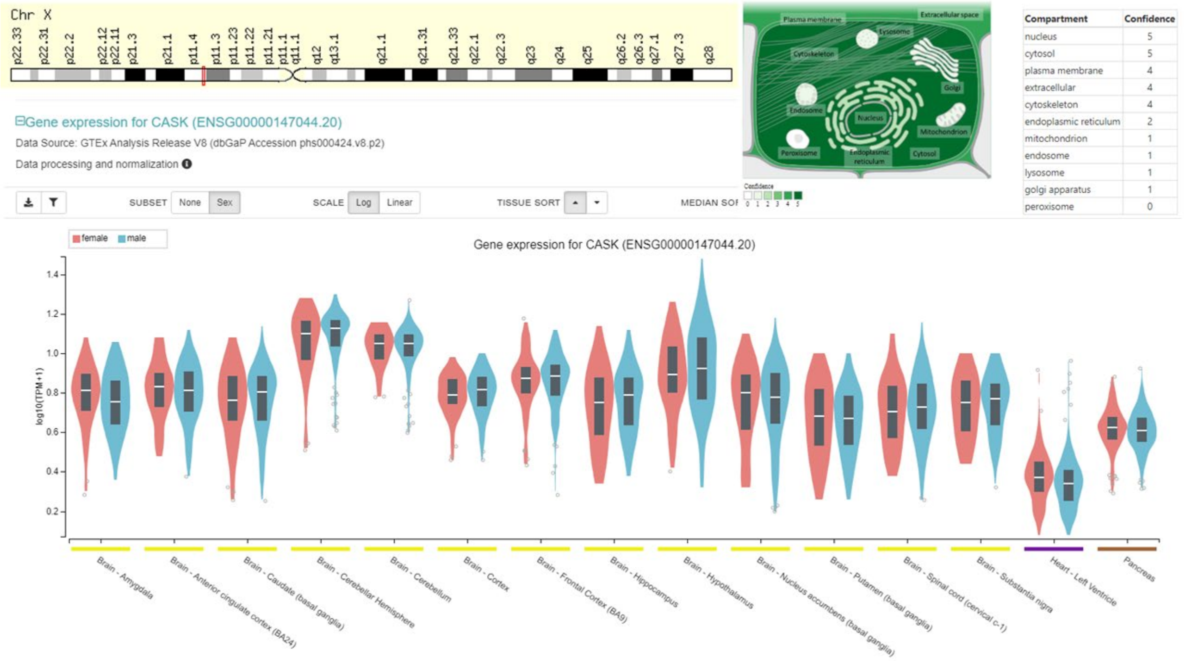


**Supplementary Figure 19. Information on gene CASK common to the SFARI Autism set and the X-Chromosome genes reported in the literature.** (OMIM 300172) The CASK gene encodes a calcium/calmodulin-dependent serine protein kinase that is a member of the membrane-associated guanylate kinase (MAGUK) protein family. MAGUKs are scaffolding proteins associated with intercellular junctions [67]. FG syndrome-4 is an X-linked recessive mental retardation syndrome characterized by congenital hypotonia, constipation, behavioral disturbances, and dysmorphic features [68, 69]. FG syndrome affects intelligence and behavior. Affected individuals tend to be friendly, inquisitive, and hyperactive, with a short attention span, have strong socialization and daily living skills. Verbal communication and language skills tend to be less developed. The physical features of FG syndrome include weak muscle tone (hypotonia), broad thumbs, and wide first (big) toes. Abnormalities of the corpus callosum are also common. Most affected individuals have constipation, and many have abnormalities of the anus such as an obstruction of the anal opening (imperforate anus). People with FG syndrome also have a distinctive facial appearance including small, underdeveloped ears; a tall, prominent forehead; and outside corners of the eyes that point downward (down-slanting palpebral fissures). (OMIM 300749) Mental retardation and microcephaly with pontine and cerebellar hypoplasia (MICPCH) is an X-linked disorder affecting ***females*** and characterized by severe intellectual disability, microcephaly, and variable degrees of pontocerebellar hypoplasia. Affected individuals have very poor psychomotor development, often without independent ambulation or speech, and axial hypotonia with or without hypertonia. Some may have sensorineural hearing loss or eye anomalies. Dysmorphic features include overall poor growth, severe microcephaly (-3.5 to -10 SD), broad nasal bridge and tip, large ears, long philtrum, micrognathia, and hypertelorism [70, 71].

Supplementary Figure 20 (FMR1 Fragile X Syndrom, FAXTAS)

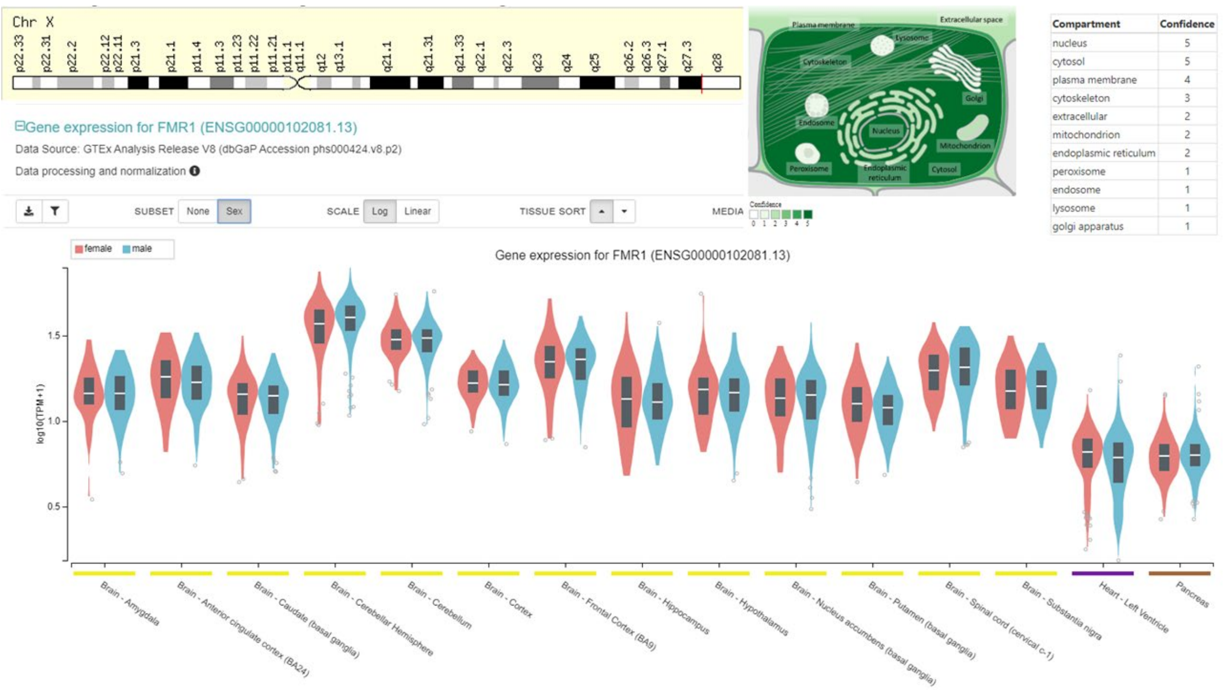


**Supplementary Figure 20. Information on gene FMR1 common to the SFARI Autism set and the X-Chromosome genes reported in the literature.** (OMIM 300624) Fragile X syndrome is characterized by moderate to severe mental retardation, macroorchidism (abnormally large testes), and distinct facial features, including long face, large ears, and prominent jaw. In most cases, the disorder is caused by the unstable expansion of a CGG repeat in the FMR1 gene and abnormal methylation, which results in suppression of FMR1 transcription and decreased protein levels in the brain [72, 73]. Jacquemont and colleagues [74, 75] provided a review of fragile X syndrome, which they characterized as a neurodevelopmental disorder, and FXTAS, which they characterized as a neurodegenerative disorder. Affected male individuals show characteristic facies, including long face, high forehead, midface hypoplasia, large mouth with long upper middle incisors, thick lips, high-arched palate, large jaw with prominent chin, and large, poorly formed ears, and macroorchidism. Mental retardation is variable, but language development is usually very delayed. Motor development is often delayed with unusual behavior, alternating anxiety and hilarity, disordered hyperactivity, and aggressiveness. Some boys with expanded FMR1 (CGG)n repeats that range in size from 55 to 200 repeats, referred to as 'premutations,' may exhibit similar, but possibly milder, clinical features to those with full expansions.

#### Supplementary Figure 21 (SLC9A6 Christianson syndrome)

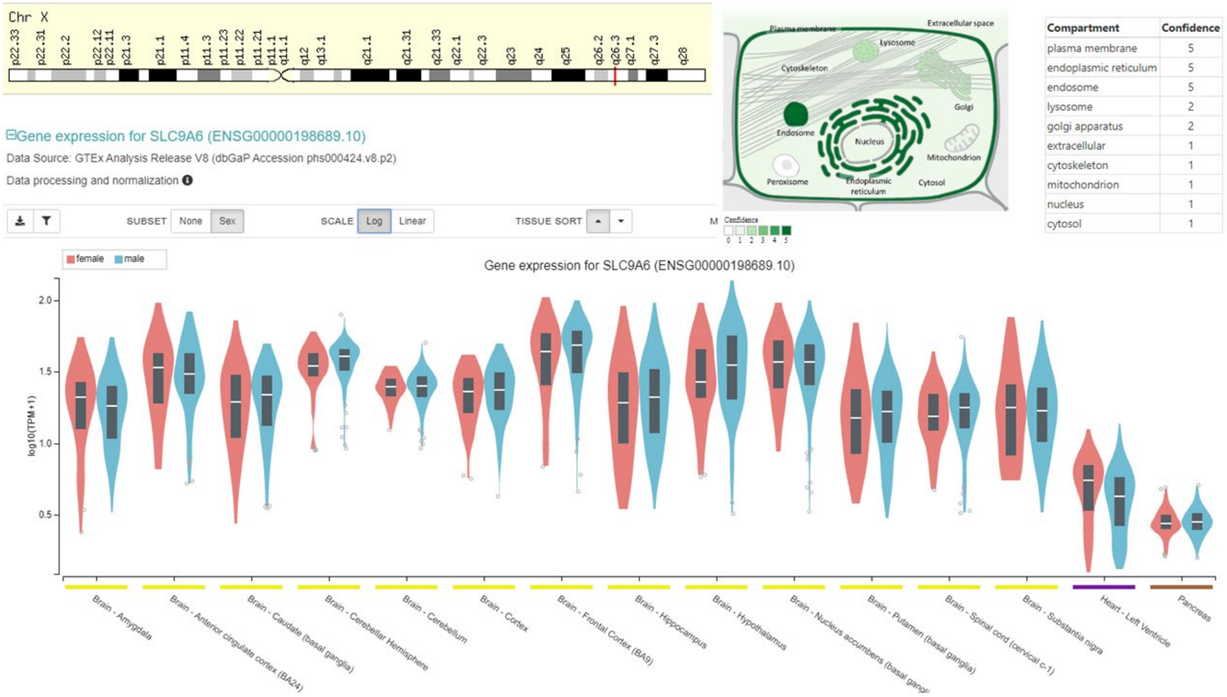


**Supplementary Figure 21. Information on gene SLC9A6 common to the SFARI Autism set and the X-Chromosome genes reported in the literature.** Christianson syndrome is an X-linked neurodevelopmental and progressive mental retardation syndrome characterized by microcephaly, impaired ocular movements, severe global developmental delay, developmental regression, hypotonia, abnormal movements, and early-onset seizures of variable types. Female carriers may be mildly affected [76, 77]. Additional features included microcephaly, absence of expressive verbal language, and slow regression of walking ability, and friendly demeanor.

#### Supplementary Figure 22 (OPHN1)

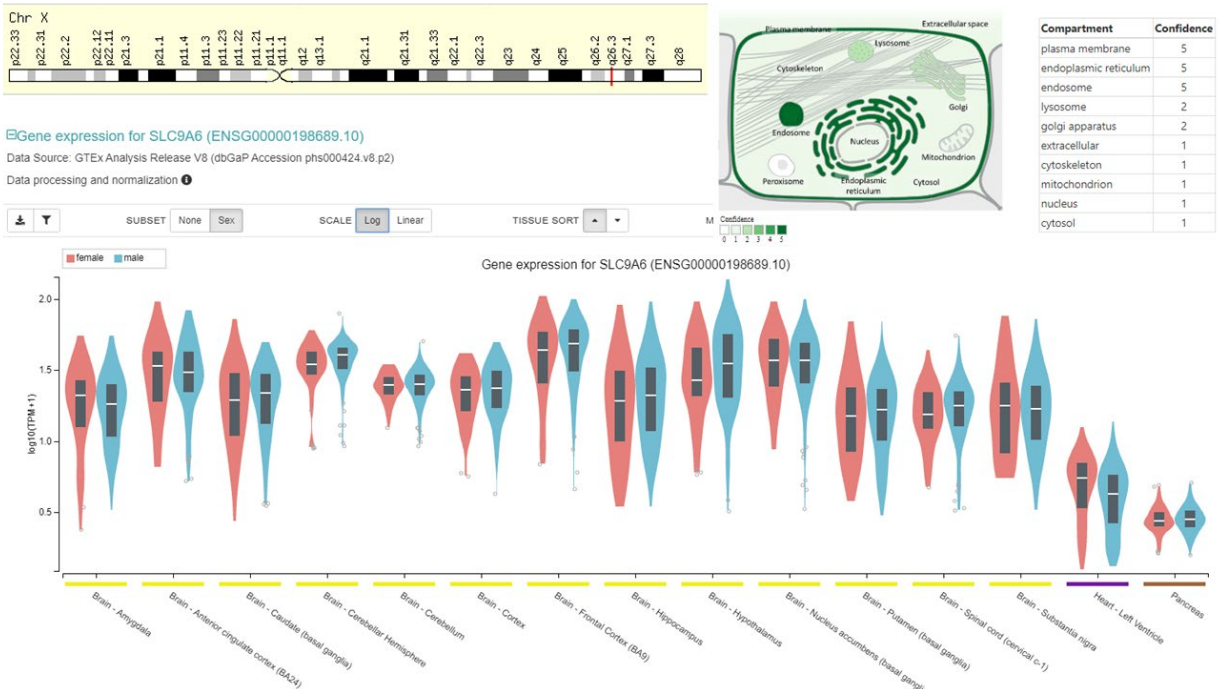


**Supplementary Figure 22. Information on gene OPHN1 common to the SFARI Autism set and the X-Chromosome genes reported in the literature.** Clinical features shared by affected individuals have been reported as generalized hypotonia and delayed psychomotor development in infancy, moderate mental retardation (IQ of 40) and autistic traits of minimal social interaction, eye contact and speech. In some cases, partial complex seizures have been reported. Physical examination has described dysmorphic facial features with long semi-triangular face, deep-set eyes, strabismus, broad high nasal root, peaked prominent nose, and prominent chin. Cerebellar signs included mild ataxia and intention tremor on finger-nose pointing. Also, at an early age, neonatal hypotonia with motor delay, marked strabismus, early-onset complex partial seizures, and moderate to severe mental retardation. Other features included hypogenitalism with cryptorchidism, hypoplastic scrotum, and microphallus. Brain MRI has showed cerebellar hypoplasia and ventriculomegaly, complex cerebellar dysgenesis including incomplete sulcation of the anterior and posterior vermis with the most prominent defect in lobules VI and VII, nonspecific cerebral corticosubcortical atrophy, expansion of the cisterna magna, and retrocerebellar cysts, with enlargement of the lateral ventricles. Various sources [78-83]
